# Heavy isotope labeling and mass spectrometry reveal unexpected remodeling of bacterial cell wall expansion in response to drugs

**DOI:** 10.1101/2021.08.06.454924

**Authors:** Heiner Atze, Filippo Rusconi, Michel Arthur

## Abstract

Antibiotics of the β-lactam (penicillin) family inactivate target enzymes called D,D-transpeptidases or penicillin-binding proteins (PBPs) that catalyze the last cross-linking step of peptidoglycan synthesis. The resulting net-like macromolecule is the essential component of bacterial cell walls that sustains the osmotic pressure of the cytoplasm. In *Escherichia coli*, bypass of PBPs by the YcbB L,D-transpeptidase leads to resistance to these drugs. We developed a new method based on heavy isotope labeling and mass spectrometry to elucidate PBP- and YcbB-mediated peptidoglycan polymerization. PBPs and YcbB similarly participated in single-strand insertion of glycan chains into the expanding bacterial side wall. This absence of any transpeptidase-specific signature suggests that the peptidoglycan expansion mode is determined by other components of polymerization complexes. YcbB did mediate β-lactam resistance by insertion of multiple strands that were exclusively cross-linked to existing tripeptide-containing acceptors. We propose that this unprecedented mode of polymerization depends upon accumulation of linear glycan chains due to PBP inactivation, formation of tripeptides due to cleavage of existing cross-links by a β-lactam-insensitive endopeptidase, and concerted cross-linking by YcbB.

## INTRODUCTION

The peptidoglycan is an essential component of the bacterial cell wall, which provides a mechanical barrier against the turgor pressure of the cytoplasm thereby preventing cells from bursting and lysing (Vollmer and Bertsche, 2008). The peptidoglycan also determines the shape of bacterial cells and is intimately integrated into the cell division process since the barrier to the osmotic pressure needs to be maintained during the entire cell cycle. These functions depend upon the net-like structure of the peptidoglycan macromolecule that is made of glycan strands cross-linked by short peptides (Fig. 1A). It is assembled from disaccharide-peptide subunits consisting of β,1→4 linked *N*-acetylglucosamine (GlcNAc) and *N*-acetyl muramic acid (MurNAc) and a stem peptide linked to the D-lactoyl group of MurNAc (Fig. 1B) (Mengin-Lecreulx et al., 1982). Polymerization of the subunit is mediated by glycosyltransferases for the elongation of glycan chains and by transpeptidases for cross-linking stem peptides carried by adjacent glycan chains.

**Figure 1.**
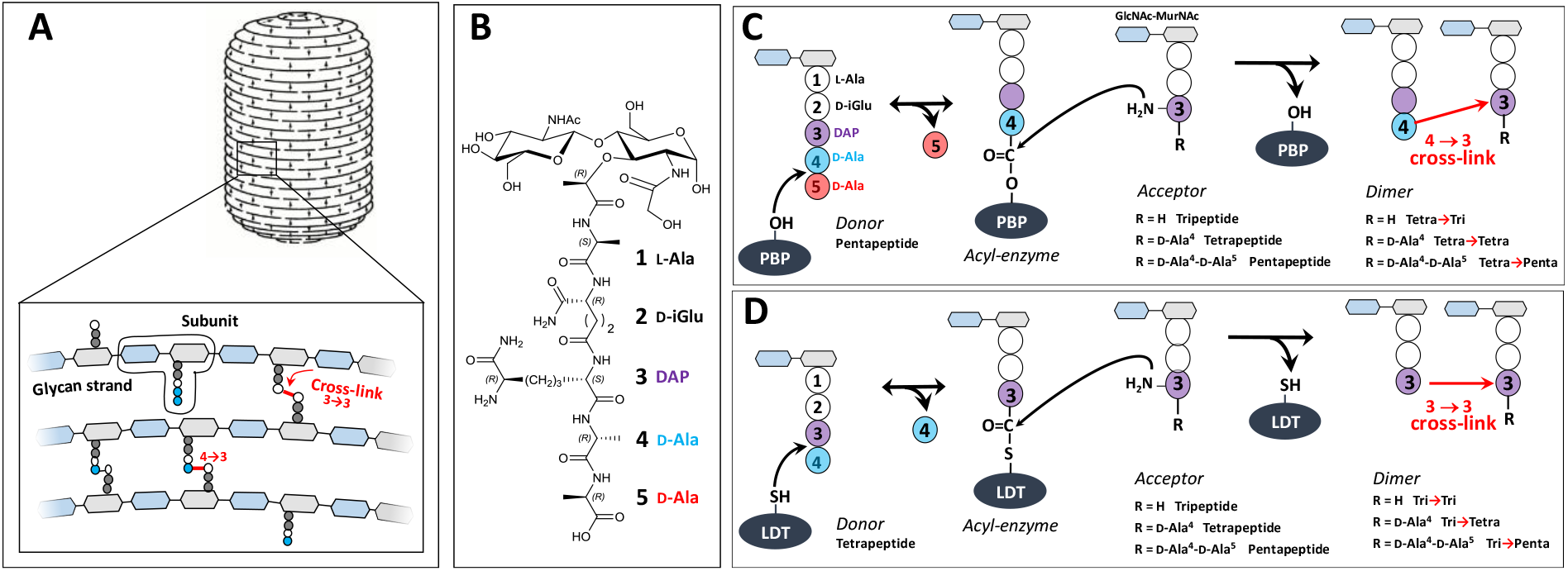
Structure and biosynthesis of *E. coli* peptidoglycan. (**A**) Structure of peptidoglycan. The net-like macromolecule is made of glycan strands cross-linked by short peptides. The polymerization of peptidoglycan involves glycosyltransferases that catalyze the elongation of glycan strands and transpeptidases that form 4→3 or 3→3 cross-links. (**B**) Structure of the peptidoglycan subunit. The stem peptide is assembled in the cytoplasm as a pentapeptide, L-Ala^1^-D-iGlu^2^-DAP^3^-D-Ala^4^-D-Ala^5^, in which D-*iso*-glutamic acid (D-iGlu) and diaminopimelic acid (DAP) are connected by an amide bond between the γ-carboxyl of D-iGlu and the L stereo-center of DAP. (**C**) Formation of 4→3 cross-links. Active-site Ser transpeptidases belonging to the penicillin-binding protein (PBP) family catalyze the formation of 4→3 cross-links connecting the carbonyl of D-Ala at the 4^th^ position of an acyl donor stem peptide to the side-chain amino group of DAP (D stereo-center) at the 3^rd^ position of an acyl acceptor stem peptide. A pentapeptide stem is essential in the donor substrate to form the acyl enzyme intermediate. The acyl acceptor potentially harbors a tripeptide, a tetrapeptide, or a pentapeptide stem leading to the formation of Tetra-Tri, Tetra-Tetra, or Tetra-Penta dimers, respectively. (**D**) Formation of 3→3 cross-links. Active-site Cys transpeptidases belonging to the L,D-transpeptidase (LDT) family catalyze the formation of 3→3 cross-links connecting two DAP residues. LDTs are specific of tetrapeptide-containing donors.

The last cross-linking step of peptidoglycan polymerization is catalyzed by two types of structurally unrelated transpeptidases (Biarrotte-Sorin et al., 2006), the L,D-transpeptidases (LDTs) (Mainardi et al., 2008) and the D,D-transpeptidases (Zapun et al., 2008), which belong to the penicillin-binding protein (PBP) family. PBPs (Fig. 1C) and LDTs (Fig. 1D) use different acyl donor substrates (pentapeptide *versus* tetrapeptide) resulting in the formation of 4→3 *versus* 3→3 cross-links, respectively (Hugonnet et al., 2016, Mainardi et al., 2005). PBPs are inactivated by all classes of β-lactams whereas LDTs are inactivated by carbapenems only (Mainardi et al., 2007). In wild type *E. coli*, LDTs are fully dispensable for growth, at least in laboratory conditions (Magnet et al., 2008). Accordingly, these enzymes have a minor contribution to peptidoglycan cross-linking in the exponential growth phase (Vollmer and Bertsche, 2008, Glauner, 1988). However, selection of mutants resistant to ceftriaxone, a β-lactam of the cephalosporin class, results in a full bypass of PBPs by the YcbB L,D-transpeptidase (Hugonnet et al., 2016). The bypass requires overproduction of the (p)ppGpp alarmone and of YcbB. The peptidoglycan of the mutant grown in the presence of ceftriaxone only contains 3→3 cross-links following PBP inhibition.

D,D-carboxypeptidase (DDC) and L,D-carboxypeptidase (LDC) activities sequentially remove D-Ala residues at the 5^th^ and 4^th^ positions of stem peptides containing a free carboxyl end, which are present in the acceptor position of dimers and in monomers. DDCs, which belong to the PBP family, hydrolyze the D-Ala^4^-D-Ala^5^ amide bond of stem pentapeptides to generate stem tetrapeptides in mature peptidoglycan (Sauvage et al., 2008). Since the reaction is nearly total in *E. coli* stem pentapeptides are found in very low abundance unless DDCs are inactivated by β-lactams. The LDC activity of LDTs hydrolyzes the DAP^3^-D-Ala^4^ amide bond generating tripeptide stems from tetrapeptide stems (Magnet et al., 2008). Since this reaction is incomplete the peptidoglycan contains combinations of tripeptide and tetrapeptide stems in monomers and in the acyl acceptor position of dimers. Removal of D-Ala residues by DDCs and LDTs occurs before or after cross-linking, leading to the polymorphism in the acceptor position of dimers depicted in Fig. 1C and 1D.

By removing D-Ala^5^ from pentapeptide stems, DDCs eliminate the essential pentapeptide donor of PBPs and generate the essential tetrapeptide donor of LDTs (Mainardi et al., 2002, Hugonnet et al., 2016). By removing D-Ala^4^ from tetrapeptide stems, LDCs eliminate the essential tetrapeptide donor of LDTs thereby potentially limiting the formation of 3→3 cross-links. Thus, DDC and LDC activities control both the extent of peptidoglycan cross-linking and the relative contributions of D,D-transpeptidases and L,D-transpeptidases to peptidoglycan cross-linking.

The interconversion between monomers and cross-linked dimers involves not only PBPs and LDTs in the biosynthetic direction but also hydrolytic enzymes, referred to as endopeptidases, which cleave the cross-links. Endopeptidases belonging to the PBP family are specific of 4→3 cross-links whereas other endopeptidases cleave both 4→3 and 3→3 cross-links (Voedts et al., 2021). Among eight endopeptidases with partially redundant functions, at least one enzyme is essential in the context of the formation of 4→3 cross-links whereas two endopeptidases, MepM and MepK, are required in the context of 3→3 cross-links (Voedts et al., 2021, Singh et al., 2012).

Morphogenesis of *E. coli* cells depends upon controlled expansion of the peptidoglycan and requires scaffolding proteins, actin (MreB) and tubulin (FtsZ) homologues (den Blaauwen et al., 2008). These proteins are involved in the recruitment of peptidoglycan polymerases and in the coordination of their activities with that of hydrolases (Vollmer and Holtje, 2001). The cell cycle involves two phases, namely the MreB-dependent elongation of the side wall at a constant diameter and the FtsZ-dependent formation of the septum at mid cell. These processes are thought to involve two distinct multienzyme complexes, the elongasome and the divisome, and specific enzymes, in particular class B PBP2 and PBP3, which mediate peptidoglycan cross-linking in the side wall and in the septum, respectively (Holtje, 1996). Accordingly, specific inactivation of PBP2 by mecillinam or of PBP3 by aztreonam results in the growth of *E. coli* as spheres or as filaments, respectively.

The average chemical composition of peptidoglycan is well known. Most enzymes involved in the formation and cleavage of glycosidic and amide bonds have been identified (at least one enzyme and generally several enzymes for each bond). However, the spatial arrangement of glycan strands is still a matter of debate. Most models propose a regular arrangement of glycan strands parallel to the cell surface and perpendicular to the long axis of the cell as depicted in Fig. 1A (Typas et al., 2011). Alternatively, it has been proposed that the glycan strands protrude perpendicularly to the surface of the cytoplasmic membrane (Meroueh et al., 2006, Dmitriev et al., 2003). The lack of direct experimental evidence originates from the heterogeneity of the peptidoglycan macromolecule that prevents the analysis of the polymer by radiocrystallography. NMR also failed to determine the structure of the polymer due to its heterogeneity and its important flexibility as measured by NMR relaxation (Kern et al., 2008). The mode of insertion of new glycan strands in the growing peptidoglycan is also not fully characterized. According to the ‘three-for-one’ model, peptidoglycan expansion involves the insertion of three neo-synthesized glycan strands to the detriment of the hydrolysis of one existing strand, which acts as a docking strand (Vollmer and Holtje, 2001). Alternatively, glycan strands may be inserted according either to a ‘one-at-a-time’ model or a ‘two-at-a-time’ model, the latter being supported by the dimeric nature of certain D,D-transpeptidases (Charpentier et al., 2002). All these models postulate that peptidoglycan expansion requires cleavage of cross-links by endopeptidases and obey to the ‘make-before-break’ principle, which proposes that cross-links in the existing peptidoglycan are hydrolyzed by endopeptidases only if the newly synthesized glycan strands are cross-linked and can sustain the osmotic pressure of the cytoplasm (Vollmer and Holtje, 2001).

At each generation, about half of the disaccharide-peptide units is released from the peptidoglycan by the combined action of endopeptidases and lytic glycosyltransferases (Goodell, 1985, Johnson et al., 2013). The latter enzymes catalyze a non-hydrolytic cleavage of the β,1→4 MurNAc-GlcNAc glycosydic bond and generate GlcNAc-anhydro-MurNAc-peptides fragments via MurNAc cyclisation at positions 1 and 6. These fragments are transported into the cytoplasm by the AmpG permease and recycled according to the pathway depicted in Supplementary Fig. S1. The key features of the recycling pathway are that the L-Ala-γ-D-Glu-DAP tripeptide is directly recycled whereas the glucosamine moiety of GlcNAc and MurNAc reenters into the peptidoglycan biosynthesis pathway at the level of glucosamine-1-phosphate. The relationships between peptidoglycan synthesis and recycling are poorly understood. The ‘three-for-one’ growth model predicts that the docking strand is recycled; conversely, the ‘one-at-a-time’ and ‘two-at-a-time’ models do not imply that recycling is a consequence of peptidoglycan synthesis.

The peptidoglycan expansion models described above were previously investigated by pulse-labeling of peptidoglycan with radioactive DAP. Indeed, the models imply specific acceptor-to-donor radioactivity ratios (ADRR) in cross-linked stem peptides (Supplementary Fig. S2) (de Jonge et al., 1989). The ADRR may potentially vary from zero to infinite for (i) exclusive cross-linking of neo-synthesized (radioactive) donor stems to existing (unlabeled) acceptor stems (new→old; ADRR = 0), (ii) the formation of cross-links between neo-synthesized (radioactive) donor and acceptor stems (new→new; ADRR = 1), and (iii) cross-linking of existing (unlabeled) donor stems to neo-synthesized (labeled) acceptor stems (old→new; infinite ADRR). This method is intrinsically inaccurate since neo-synthesized subunits issued from the recycling of stem peptides are mistakenly identified as material from the existing wall. Moreover, the method provides access to the average composition of dimers in labeled donor and acceptor stems rather than the labeling status of stem peptides that have directly participated in the same cross-linking reaction. Thus, an ADRR of 1 can originate both from dimers containing neo-synthesized stem peptides in the donor and acceptor positions (new→new) or, alternatively, from a combination of dimers containing a radioactive DAP either in the donor (new→old) or acceptor (old→new) positions in equimolar amounts. This limitation was disregarded in previous studies since dimers containing a single labeled DAP residue in the donor position are considered to be rare, at least in wild-type *E. coli* grown in the absence of β-lactam. This is arguably the case for dimers generated by PBPs since pentapeptide stems are rapidly converted to tetrapeptide stems by D,D-carboxypeptidases thereby precluding existing stems from participating as donors in the formation of 4→3 cross-links by D,D-transpeptidases. Obviously, this premise does not apply to LDTs since these enzymes use tetrapeptide stems as donors (Fig. 1). Here, we describe a new method based on full labeling of the entire peptidoglycan macromolecule with stable isotopes of carbon and nitrogen combined with high resolution mass spectrometry (MS). This method does not suffer from the limitations listed above and our results show that it enables a thorough investigation of peptidoglycan synthesis and recycling in a single experiment. We applied our method to the comparison of the modes of expansion of the peptidoglycan in bacterial strains relying on PBPs, and, for the first time, on L,D-transpeptidase YcbB, or on both types of enzymes for peptidoglycan cross-linking. We show how previously described β-lactam-induced malfunctioning of the bacterial cell wall synthesis machinery (Cho et al., 2014) affects peptidoglycan synthesis and recycling. We also report how L,D-transpeptidase YcbB rescues inhibition of PBPs by participating in an unprecedented mode of peptidoglycan polymerization. The results described in this report were obtained thanks to the richness of the structural information brought by MS and MS/MS analyses of all the labeled/unlabeled sugar and amino acid residues of peptidoglycan subunits, as opposed to the radioactive signal located in the DAP moiety only. The mass data analysis pipeline automating the identification of labeled muropeptides is described in this study and may be generally applied to the screening of the mode of action of antibacterial agents acting on peptidoglycan synthesis.

## RESULTS

### Muropeptide composition of the peptidoglycan from *E. coli* M1.5

Strain M1.5 was chosen for initial experiments because D,D-transpeptidases and L,D-transpeptidases both have major contributions to peptidoglycan polymerization offering the possibility to study 4→3 and 3→3 cross-linked dimers from the same peptidoglycan preparation. Strain M1.5 was grown in labeled M9 minimal medium containing [^15^N]NH4Cl and [^13^C]glucose as the sole sources of nitrogen and carbon, respectively. *E. coli* M1.5 was also grown in unlabeled M9 minimal medium. Peptidoglycan was extracted from exponentially growing cultures and digested by muramidases, which generate soluble disaccharide-peptides. The resulting muropeptides were reduced with NaBH4 and purified by *rp*HPLC. Mass spectrometric analyses (see below) were performed to determine the structure of the six predominant muropeptides. These muropeptides included two monomers containing either a tripeptide (L-Ala^1^-D-iGlu^2^-DAP^3^) or a tetrapeptide (L-Ala^1^-D-iGlu^2^-DAP^3^-D-Ala^4^) stem, both linked to a reduced GlcNAc-MurNAc disaccharide. The remaining four muropeptides were dimers containing a 3→3 or a 4→3 cross-link with a tripeptide or a tetrapeptide stem in the acceptor position in all four combinations. As expected, growth in the labeled *versus* unlabeled medium did not alter the muropeptide composition of peptidoglycan. The average abundance of the six predominant muropeptides is reported in Supplementary Table S1.

### Determination of the isotopic composition of the unlabeled monomers

The mass spectra of monomers from *E. coli* grown in unlabeled M9 minimal medium displayed the conventional isotopic clusters (Fig. 2 and Supplementary Fig. S3 for the disaccharide-tripeptide and - tetrapeptide, respectively), as expected for the natural abundances of carbon, hydrogen, nitrogen, and oxygen isotopes (See Supplementary Material and Methods for the method used to deduce the mass and abundance of isotopologues from the isotopic composition of the medium). For the disaccharide-tripeptide, the most abundant isotopologue (64.382%) only contained the light isotopes of the four chemical elements. The second peak of the isotopic cluster was composed of isotopologues containing only light isotopes except for either one ^13^C (23.497%), ^2^H (0.437%), ^15^N (1.411%), or ^17^O (0.492%) isotope, together representing 25.877% of the relative abundance of the most abundant isotopologues. The overlay of the experimental and simulated spectra (Fig. 2C) confirmed the expected isotopologue composition (See Supplementary Material and Methods for the method used to simulate mass spectra by using the deduced mass and abundance of isotopologues).

**Figure 2.**
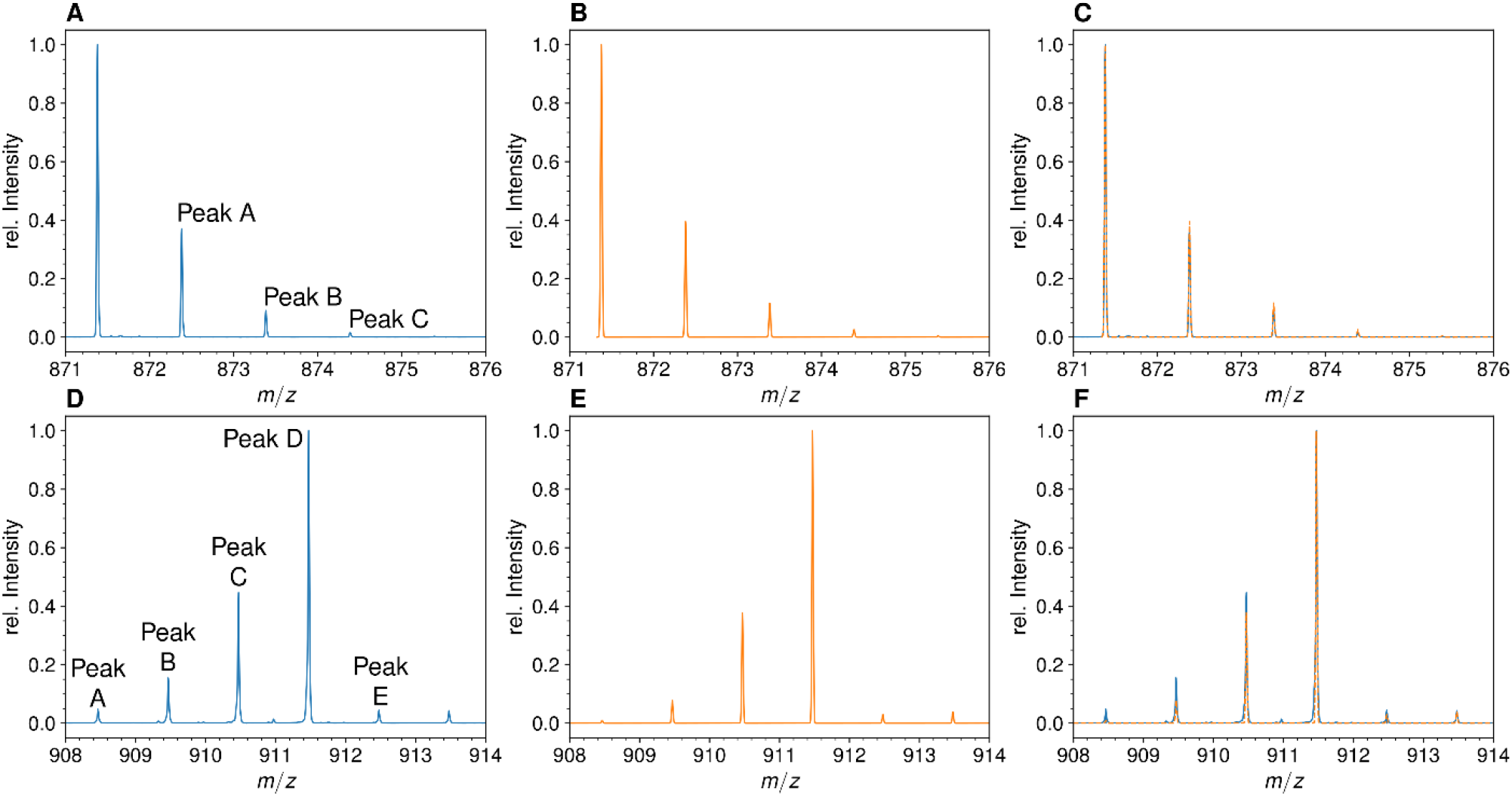
Mass spectra of the unlabeled and fully labeled disaccharide-tripeptide monomers. (**A**) Experimental mass spectrum of the mono-protonated ([M+H]^1+^) disaccharide-tripeptide (C34H58N6O20•H^+^) extracted from *E. coli* M1.5 grown in unlabeled M9 minimal medium. (**B**) Simulated spectrum obtained for the natural abundance of carbon and nitrogen isotopes. (**C**) Overlay of the spectra in A and B. (**D**) Observed mass spectrum of the GlcNAc-MurNAc-tripeptide purified from the peptidoglycan of *E. coli* M1.5 grown in the labeled M9 minimal medium. (**E**) Simulated mass spectrum obtained for 99% ^13^C and ^15^N labeling. (**F**) Overlay of spectra in D and E.

### Determination of the isotopic composition of the fully labeled monomers

Mass spectrometric analyses were performed on the reduced disaccharide-tripeptide monomer purified from the peptidoglycan of strain M1.5 grown in the labeled medium (Fig. 2D). The theoretical mass spectra were simulated as described in the Supplementary Material and Methods section (Fig. 2E) and overlaid to the experimental spectra (Fig. 2C), revealing a good match between the two types of spectra. For the labeled tripeptide, the most intense peak at *m/z* 911.473 (*z* = 1) (Fig. 2D) corresponds to cluster D described in the Supplementary Material and Methods, which is mainly accounted for by the uniformly labeled disaccharide-tripeptide (*m/z* value of 911.474). Additional cluster peaks at *m*/*z* values of 908.463, 909.467, and 910.469 (Fig. 2D), correspond to clusters A, B, and C (*m/z* values of 908.467, 909.469, and 910.472). These peaks correspond to uniformly labeled species except for the presence of combinations of 3, 2, and 1 light ^12^C or ^14^N nuclei, respectively. The presence of these peaks fits the simulated spectra that were calculated with the extent of [^15^N]NH_4_Cl and [^13^C]glucose labeling reported by the manufacturers (99% for each isotope). The same conclusion was drawn from the comparison of the simulated and experimental spectra of the labeled disaccharide-tetrapeptide (Supplementary Fig. S3).

Qualitatively, it is worth noting that the mass spectra of the unlabeled and labeled disaccharide– tripeptide presented in Fig. 2 are almost mirror images. This reflects the preponderance of the isotopologue containing only “light” isotopes in the unlabeled muropeptide followed (to the right) by peaks of decreasing intensity resulting from incorporation of a few rare heavy isotopes. In contrast, the peak of the fully labeled disaccharide-peptide is preceded (to the left) by peaks of decreasing masses due to incorporation of a few rare ^12^C and ^14^N nuclei. The same conclusions apply to the spectrum of the unlabeled disaccharide-tetrapeptide monomer (Supplementary Fig. S3).

The structure of the isotopologues was determined by tandem mass spectrometry. As an example, Table 2 presents the MS/MS data obtained for mono-protonated isotopologue ions of the labeled disaccharide-tetrapeptide (C_37_H_63_N_7_O_21_•H^+^) under isotopic cluster peaks at *m/z*_*obs*_ 986.519 and 985.516. These values match the calculated *m/z*_*cal*_ values for isotopologues either exclusively containing the ^13^C and ^15^N isotopes (*m*/*z* 986.518) or bearing a single light isotope of either element (*m/z*_*cal*_ 985.515 or 985.521 for a ^12^C or ^14^N isotope, respectively). Fragmentation of the isotopologues under the peak at *m/z*_*obs*_ of 986.519 resulted in the expected set of product ions for a uniformly labeled disaccharide-tetrapeptide, while two sets of product ions, differing by approximately one mass unit, were observed for the fragmentation of isotopologues bearing a single light isotope under the peak at *m/z*_*obs*_ of 985.516. The fragments in the lower-mass ion set had retained their single light nucleus (^12^C or ^14^N) while the fragments in the higher-mass ion set had lost their single light nucleus and were fully labeled (these higher-mass fragments matched exactly those obtained for the fully labeled disaccharide-tetrapeptide). Thus, each fragment was present in the spectrum as a pair of peaks that reflected the presence or the absence of the light isotope. Table 2 provides the relative intensity of these two peaks for each fragment of the isotopologues. The relative intensity of the peak containing the light isotope decreased with the mass of the fragment. This result is expected for a uniform distribution of the light isotope in both the sugar and peptide moieties of the disaccharide-tetrapeptide because a smaller number of nuclei (smaller mass) makes it more probable to lose the single light nucleus during fragmentation.

Taken together, these results show that we successfully achieved extensive peptidoglycan labeling that was only limited by the extent (99%) of the labeling of glucose and NH_4_Cl introduced in the labeled growth medium. Thus, isotopic dilution due to atmospheric N_2_ and CO_2_ was not an issue, as expected from the metabolic fluxes of *E. coli* grown in minimal medium. Note that the pre-culture and the culture were both performed in labeled M9 medium to avoid any contribution of the initial inoculum to the final isotopic composition of the peptidoglycan. Simulation of high-resolution mass spectra was successfully used to assign isotopologues of defined structure and isotopic composition to the corresponding peaks in experimental mass spectra (Fig. 2 and Supplementary Fig. S3). These assignments were confirmed by tandem mass spectrometry (shown for two isotopologues in Table 1).

**Table 1.**
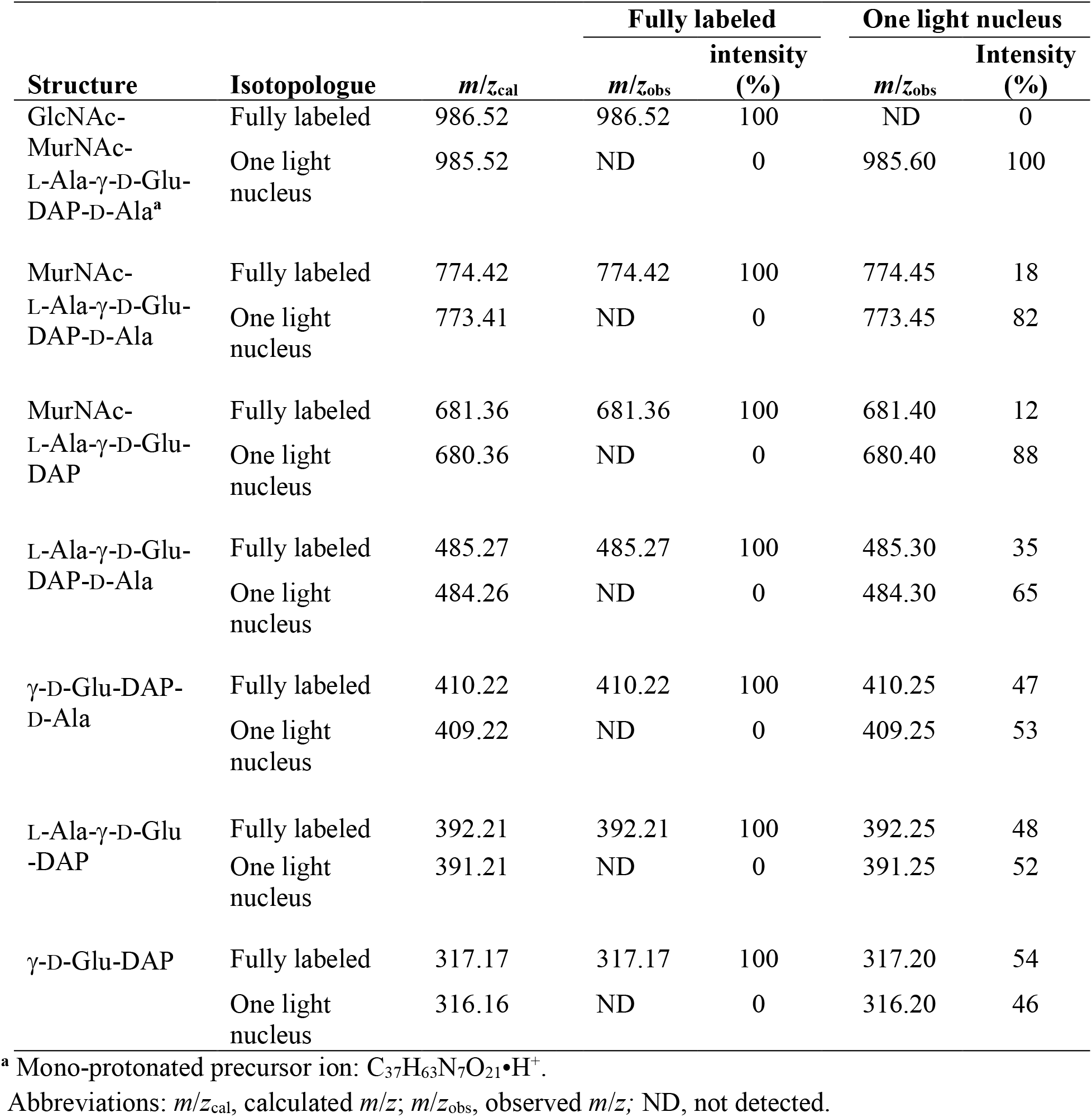
Tandem mass spectrometry analysis of isotopologues of the GlcNAc-MurNAc-tetrapeptide.

| Structure | Isotopologue | $m/z_{\text{cal}}$ | Fully labeled | | One light nucleus | |
| --- | --- | --- | --- | --- | --- | --- |
| | | | $m/z_{\text{obs}}$ | intensity (%) | $m/z_{\text{obs}}$ | Intensity (%) |
| GlcNAc-MurNAc-L-Ala-γ-D-Glu-DAP-D-Ala<sup>a</sup> | Fully labeled | 986.52 | 986.52 | 100 | ND | 0 |
|  | One light nucleus | 985.52 | ND | 0 | 985.60 | 100 |
| MurNAc-L-Ala-γ-D-Glu-DAP-D-Ala | Fully labeled | 774.42 | 774.42 | 100 | 774.45 | 18 |
|  | One light nucleus | 773.41 | ND | 0 | 773.45 | 82 |
| MurNAc-L-Ala-γ-D-Glu-DAP | Fully labeled | 681.36 | 681.36 | 100 | 681.40 | 12 |
|  | One light nucleus | 680.36 | ND | 0 | 680.40 | 88 |
| L-Ala-γ-D-Glu-DAP-D-Ala | Fully labeled | 485.27 | 485.27 | 100 | 485.30 | 35 |
|  | One light nucleus | 484.26 | ND | 0 | 484.30 | 65 |
| γ-D-Glu-DAP-D-Ala | Fully labeled | 410.22 | 410.22 | 100 | 410.25 | 47 |
|  | One light nucleus | 409.22 | ND | 0 | 409.25 | 53 |
| L-Ala-γ-D-Glu-DAP | Fully labeled | 392.21 | 392.21 | 100 | 392.25 | 48 |
|  | One light nucleus | 391.21 | ND | 0 | 391.25 | 52 |
| γ-D-Glu-DAP | Fully labeled | 317.17 | 317.17 | 100 | 317.20 | 54 |
|  | One light nucleus | 316.16 | ND | 0 | 316.20 | 46 |

<sup>a</sup> Mono-protonated precursor ion: C<sub>37</sub>H<sub>63</sub>N<sub>7</sub>O<sub>21</sub>•H<sup>+</sup>.
Abbreviations: $m/z_{\text{cal}}$ , calculated $m/z$ ; $m/z_{\text{obs}}$ , observed $m/z$ ; ND, not detected.

### Mass spectra of the four dimers

Peptidoglycan dimers do not harbor any symmetry axis as one peptide stem acts as an acyl donor in the transpeptidation reaction whereas the other acts as an acyl acceptor and retains the single free carboxyl extremity of the peptide moiety (Fig. 1). By convention, the structure of dimers is represented by placing the donor on the left separated from the acceptor by an arrow (donor→acceptor). In *E. coli* M1.5, the YcbB L,D-transpeptidase catalyzes the formation of DAP^3^→DAP^3^ cross-links connecting two diaminopimelyl residues (DAP^3^) located at the third position of a donor stem (a tripeptide) and of an acceptor stem (a tripeptide or a tetrapeptide), thus generating Tri→Tri and Tri→Tetra dimers, respectively. These dimers only differ by the presence or absence of D-Ala at the C-terminal end of the acceptor stem. The D,D-transpeptidases of *E. coli* M1.5 catalyze the formation of D-Ala^4^→DAP^3^ cross-links connecting a tetrapeptide donor stem to an acceptor stem containing either a tripeptide (Tetra→Tri dimer) or a tetrapeptide (Tetra→Tetra dimer). In these dimers, D-Ala at the fourth position of the donor stem is engaged in the D-Ala^4^→DAP^3^ cross-link and DAP^3^ or D-Ala^4^ occupies the C-terminal position of the acceptor stem. The Tri→Tetra and Tetra→Tri isomers formed by YcbB and the D,D-transpeptidases can be discriminated by tandem mass spectrometry based on the cleavage of the DAP^3^→DAP^3^ and D-Ala^4^→DAP^3^ cross-links, respectively, and on the loss of a C-terminal D-Ala only present in the acceptor stem of the dimer generated by YcbB (Supplementary Data). As detailed above for monomers, the comparison of the experimental and simulated mass spectra showed that the observed isotopic clusters of the dimers can be accounted for by the isotopic composition of the growth media (see Supplementary Fig. S4 for the Tetra→Tetra dimer). The structure of all dimers was confirmed by tandem mass spectrometry (Supplementary Data).

### Study design for the kinetics of incorporation of unlabeled subunits into the fully labeled peptidoglycan of strain M1.5

To investigate the mode of insertion of peptidoglycan precursors into sacculi, *E. coli* M1.5 was grown in minimal medium to an OD_600_ of 0.4. Bacteria were collected by centrifugation and resuspended in the unlabeled medium. Incubation was continued in the same conditions and samples were collected at 5, 30, 45, 65, and 85 min after the medium switch (Fig. 3A), representing a little less than a generation (90 min; Supplementary Table S2). Peptidoglycan was extracted and the six predominant muropeptides, two monomers and four dimers, were purified by *rp*HPLC (Fig. 3B to 3F) and identified by mass spectrometry (Fig. 3G). The relative abundance of these six muropeptides remained similar in all samples indicating, as expected, that the medium switch did not alter the overall peptidoglycan composition.

**Figure 3.**
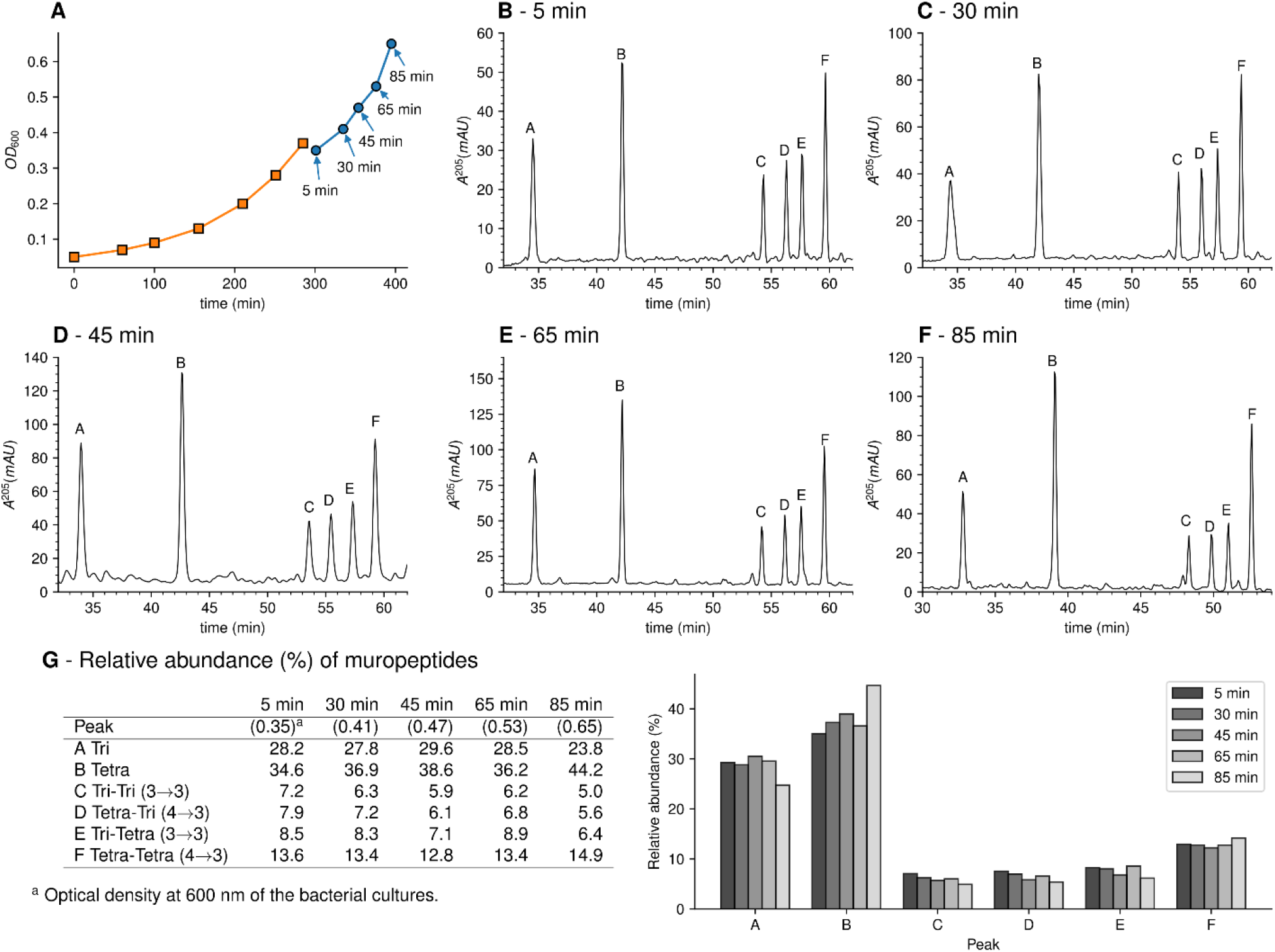
Muropeptide profile of peptidoglycan extracted from *E. coli* M1.5 after the medium switch. (**A**) *E. coli* M1.5 was inoculated in the labeled minimal medium and bacterial growth was monitored by determining the OD600 (orange curve, squares). When the optical density reached 0.4, bacteria were collected by centrifugation, resuspended in the unlabeled minimum medium, incubation was continued (blue curve), and samples were withdrawn at 5, 30, 45, 65, and 85 min (circles). (**B** to **F**) Peptidoglycan was extracted from these five culture samples, digested with muramidases, and the resulting muropeptides were separated by *rp*HPLC. (**G**) Identification and relative abundance of muropeptides in the major chromatographic peaks.

### Impact of the medium switch on the isotopic composition of monomers

Mass spectrometry revealed that growth in the unlabeled medium after the switch led to the appearance of disaccharide-tripeptide isotopologues of natural isotopic composition (Fig. 4A). These isotopologues were exclusively assembled from unlabeled glucose and NH_4_Cl after the medium switch. In addition, minor amounts of hybrids containing both labeled and unlabeled moieties were detected (named h1, h2, and h3 in Fig. 4A). Tandem mass spectrometry (Supplementary Data) indicated that hybrid h1 consists of two types of isotopologues containing a labeled hexosamine moiety either in the GlcNAc or MurNAc residue. Hybrid h2 contained a labeled stem peptide and an unlabeled disaccharide. Hybrid h3 was uniformly labeled except for GlcNAc. Hybrids h1, h2, and h3 could originate from the incorporation of recycled fragments of existing (labeled) peptidoglycan. This requires digestion of the existing peptidoglycan by hydrolases and import of the digested fragments by specialized permeases (AmpG and Opp) (Johnson et al., 2013). Alternatively, the hybrids could originate from an intracellular pool of labeled precursors present prior to the medium switch. The structures of h1 and h2 were fully accounted for by the known peptidoglycan recycling pathway of *E. coli* (Johnson et al., 2013) (Supplementary Fig. S1). Indeed, GlcNAc and MurNAc are recycled into glucosamine 6-phosphate, a common precursor of UDP-GlcNAc and UDP-MurNAc. This pathway could therefore account for the incorporation of a recycled (labeled) hexosamine into the GlcNAc or MurNAc residues of the h1 isotopologues. Regarding h2, the peptide moiety of peptidoglycan fragments is recycled as a tripeptide, which is added to UDP-MurNAc by the Mpl synthetase (Mengin-Lecreulx et al., 1996). This pathway could be responsible for formation of the h2 hybrid containing a labeled (recycled) tripeptide and an unlabeled GlcNAc-MurNAc disaccharide. In contrast, the formation of the h3 hybrid did not depend upon recycling since MurNAc-peptide moieties are not selectively recycled as single molecules. Thus, the h3 hybrid originated from moieties that were assembled after (unlabeled GlcNAc) or prior to (labeled MurNAc-peptide) the medium switch. Accordingly, the cytoplasm of *E. coli* contains an important pool of UDP-MurNAc-pentapeptide (an estimate of 120,000 molecules per bacterial cell), which represents *ca*. 3% of the total number of disaccharide-peptide units present in the cell wall (Mengin-Lecreulx et al., 1982). These observations indicate that h3 is the sole hybrid assembled independently from recycling.

**Figure 4.**
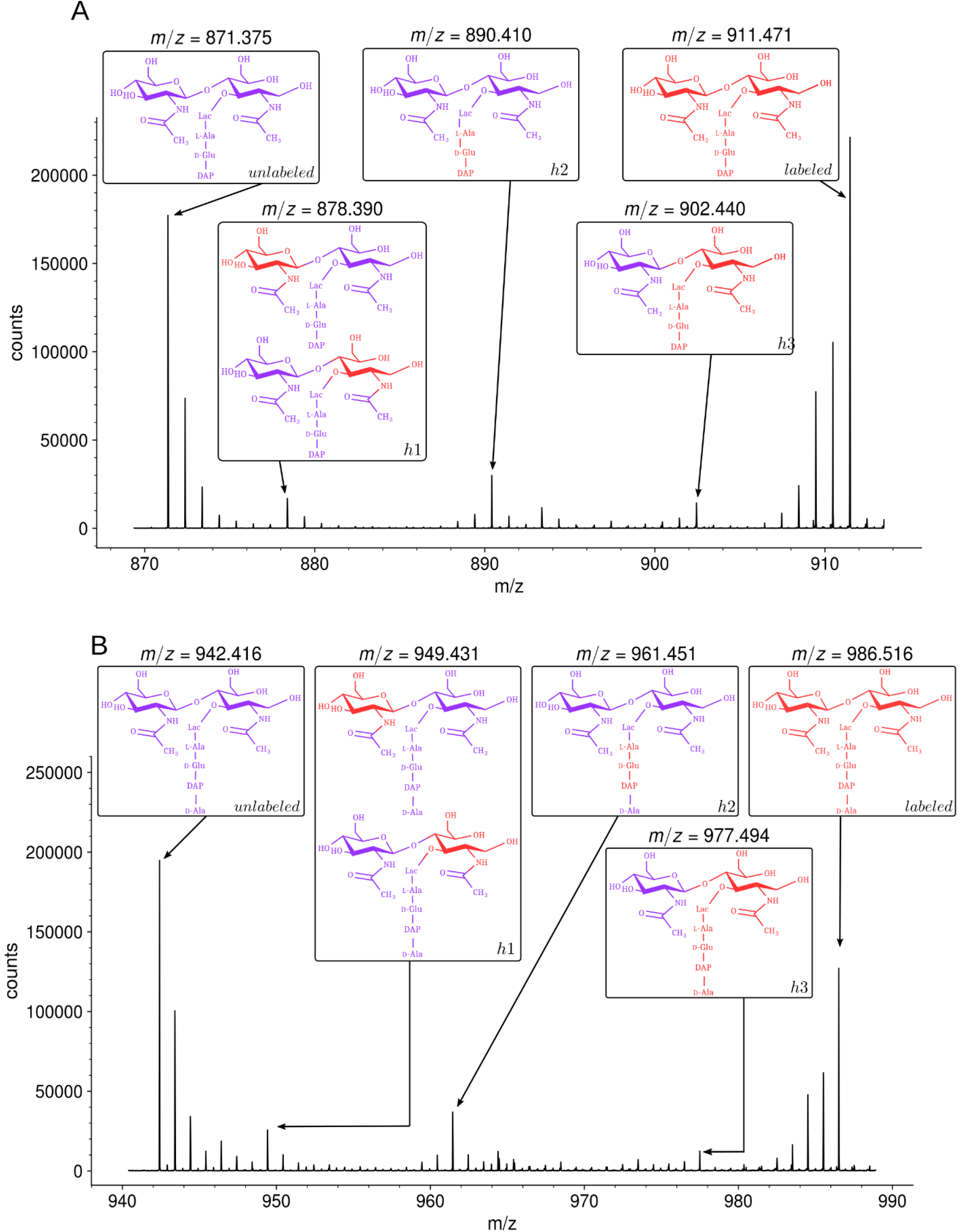
Isotopic composition of monomers isolated from the peptidoglycan of *E. coli* M1.5 after the medium switch. (**A**) Mass spectrum obtained for the GlcNAc-MurNAc-tripeptide monomer recovered in peak A of the chromatogram in Fig. 3D. (**B**) Mass spectrum obtained for the GlcNAc-MurNAc-tetrapeptide monomer recovered in peak B of the chromatogram in Fig. 3D. Purple and red colors indicate the unlabeled (^12^C and ^14^N) or labeled (^13^C and ^15^N) isotopic content of the amino acid and sugar residues of the monomers, respectively.

The role of recycling in the formation of the isotopologues was further investigated by constructing a derivative of *E. coli* M1.5 deficient in peptidoglycan recycling following deletion of the *ampG* permease gene. In this mutant, h1 hybrids were not detected indicating that they originated from incorporation of recycled glucosamine (Supplementary Fig. S5). Hybrids h2 and h3 were present in the peptidoglycan of the Δ*ampG* mutant as expected from the metabolic chart presented in Supplementary Fig. S1. Indeed, recycling of the tripeptide (hybrid h2) involves both the transport of anhydroMurNAc-peptides by AmpG and the transport of the tripeptide by the Opp permease (Johnson et al., 2013). Deletion of the *ampG* gene is therefore expected to reduce rather than abolish formation of the h2 hybrid. The h3 hybrid was proposed to originate from existing UDP-MurNAc-pentapeptide and *de novo* synthesized GlcNAc moieties (above). Accordingly, the *ampG* deletion did not prevent formation of the h3 hybrid.

Analysis of the disaccharide-tetrapeptide (Fig. 4B and Supplementary Data) revealed a pattern of incorporation of unlabeled moieties very similar to that described above for the disaccharide-tripeptide. After the medium switch, the bulk of the ^12^C and ^14^N nuclei was incorporated into uniformly unlabeled disaccharide-tetrapeptides. Hybrid molecules retained labeled glucosamine moieties in GlcNAc or MurNAc (h1), a labeled tripeptide moiety in the tetrapeptide stem (h2), or labeling of the entire molecule except for GlcNAc (h3). As expected, the h1 hybrid was not detected in the Δ*ampG* mutant (Supplementary Fig. S5). The hybrid tetrapeptide stem present in h2 is in full agreement with the *E. coli* recycling pathway as the tripeptide and the terminal D-Ala are expected to have different origins. Indeed, the terminal D-Ala of recycled tetrapeptide stems is cleaved off by a cytoplasmic L,D-carboxypeptidase to form a tripeptide that is added to MurNAc by the Mpl synthetase (Templin et al., 1999). In the following step, MurF, adds the D-Ala-D-Ala dipeptide to generate the complete pentapeptide stem.

The isotopologues described above generated the isotopic clusters expected from their isotopic composition. The basis for the mirror image of the isotopic clusters of muropeptides exclusively generated from either labeled or unlabeled glucose and NH_4_Cl has already been introduced in the text and in Fig. 2. The major h2 isotopologue mass peak had minor mass peaks both at lower and higher *m*/*z* values generating a seemingly symmetrical cluster (Fig. 4A and B; Supplementary Fig. S6 for simulated spectra). Peaks at lower *m*/*z* values mostly originated from the presence of rare ^12^C and ^14^N nuclei in the labeled moiety of the molecule. Peaks at higher *m*/*z* values mostly originated from the presence of rare ^13^C and ^15^N nuclei in the unlabeled moiety of the molecule. The contribution of these two effects shaped a nearly symmetrical isotopic cluster centered on the major isotopologue mass peak exclusively containing (i) ^12^C and ^14^N nuclei in the unlabeled moiety of the molecule, (ii) ^13^C and ^15^N nuclei in the labeled moiety, and (iii) ^1^H and ^16^O nuclei in the entire molecule.

### Kinetics of incorporation of unlabeled nuclei into muropeptide monomers

Our next objective was to compare the evolution of the ratios of uniformly labeled and unlabeled monomers following the culture medium switch. This analysis was based on the mass spectra of muropeptides isolated from culture samples withdrawn at various times after the medium switch as described in Fig. 3. The mass spectra obtained for the disaccharide-tripeptide and disaccharide-tetrapeptide are displayed in Fig. 5A and 5C, respectively. Variations in the relative abundance of the uniformly labeled and unlabeled isotopologues deduced from mass spectral ion intensities are presented in Fig. 5B and 5D. For the disaccharide-tripeptide, replacement of the uniformly labeled isotopologue by its unlabeled counterpart reached *ca*. 50 % in 85 min (Fig. 5B). In contrast, the replacement was more rapid for the disaccharide-tetrapeptide (*ca*. 94 % in 85 min) (Fig. 5D). Thus, the isotopic labeling method resolved different kinetics of accumulation of the neo-synthesized monomers (Fig. 5) although their relative proportions in the peptidoglycan remained similar (Fig. 3). The tetrapeptide and tripeptide stems may originate from sequential hydrolytic reactions involving the removal of D-Ala^5^ from free (uncross-linked) pentapeptide stems by D,D-carboxypeptidases, followed by the removal of D-Ala^4^ from the resulting tetrapeptide by the L,D-carboxypeptidase activity of YcbB (Magnet et al., 2008). The sequential nature of these reactions may explain, at least in part, the delayed accumulation of the disaccharide-tripeptide, as it derives from the disaccharide-tetrapeptide.

**Figure 5.**
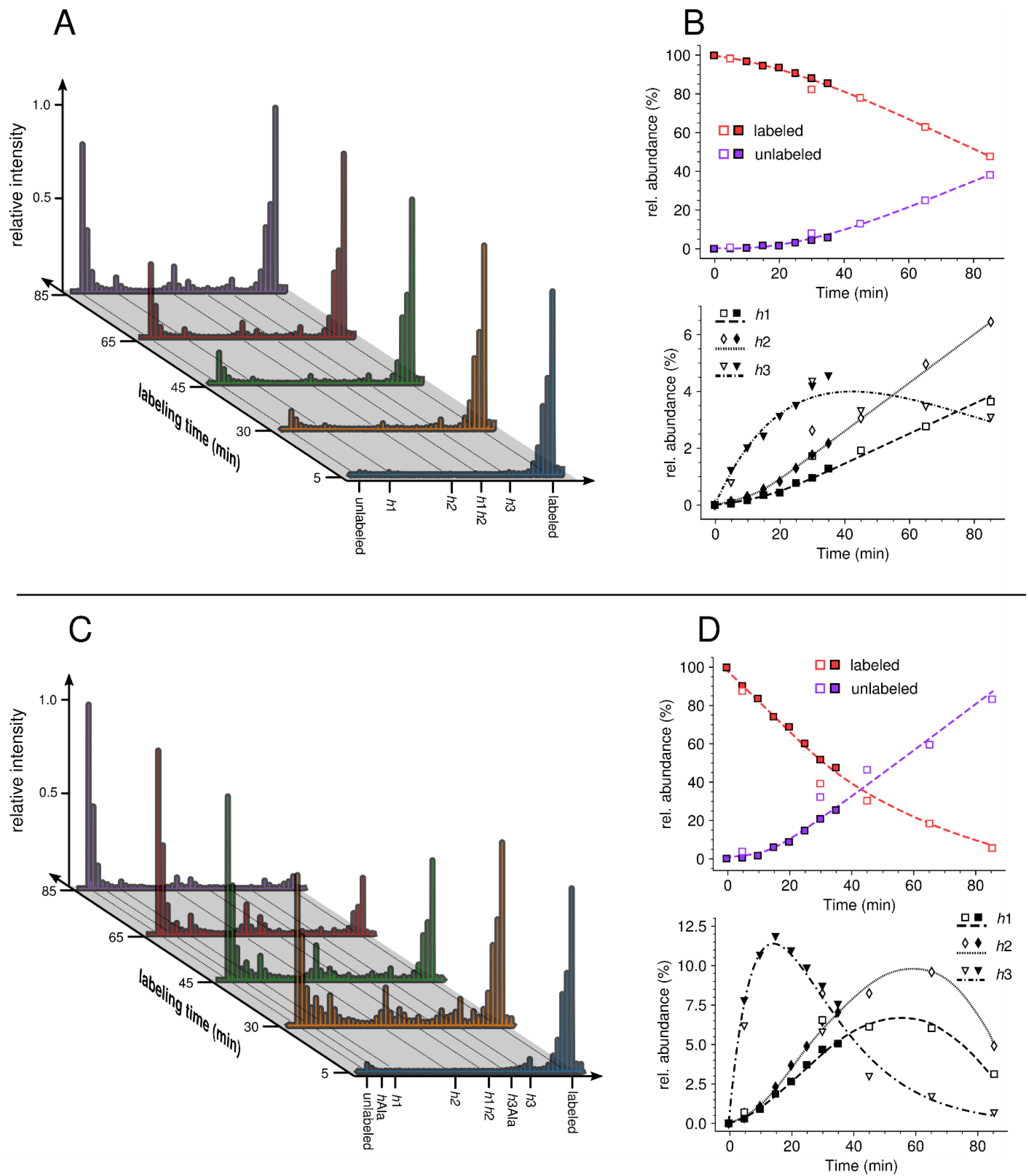
Kinetic analysis of the proportion of isotopologue monomers in the peptidoglycan of strain M1.5. (**A**) and (**C**), mass spectra of disaccharide-tripeptide and disaccharide-tetrapeptide from bacteria collected at 5, 30, 45, 65, and 85 min after the medium switch. (**B**) and (**D**), kinetic analysis of the ratios of isotopologues calculated from relative ion current intensities for the disaccharide-tripeptide and disaccharide-tetrapeptide, respectively. The upper panels show the evolution of the relative abundance of uniformly labeled and unlabeled monomers. The lower panels present the evolution of the relative abundance of hybrids relative to the complete set of monomers. The open and solid symbols are the results from two biological repeats. See Supplementary Data for the complete assignment of hybrid structures.

Kinetics of accumulations of hybrids revealed that hybrids h1 and h2, which originated from peptidoglycan recycling, were virtually absent at 5 min and increased between 5 and 85 min for the disaccharide-tripeptide and between 5 and 30 min for the disaccharide-tetrapeptide (Fig. 5). This is expected since recycled building blocks (glucosamine and tripeptide for h1 and h2, respectively) originated from a peptidoglycan that was fully labeled at the time of the medium switch and remained partially labeled during the time course of the experiment. In contrast, h3 hybrids were already present at 5 min, and reached a maximum at *ca*. 18 and 30 min for tetrapeptide and tripeptide, respectively, as expected for their synthesis from a cytoplasmic pool of labeled UDP-MurNAc-pentapeptide present at the time of the medium switch.

### Impact of the medium switch on the isotopic composition of dimers

Three main types of isotopologues were observed after the medium switch for each of the four dimers (Fig. 6 for the Tri→Tri dimer and Supplementary Fig. S7, S8, and S9 for the Tetra→Tri, Tri→Tetra, Tetra→Tetra dimer, respectively). The first type of isotopologues were uniformly labeled. These dimers were generated by the cross-linking of stems peptides assembled before the medium switch. Thus, they correspond to fragments of the existing peptidoglycan. These dimers are referred to as old→old isotopologue dimers. The second type of isotopologues was generated by the cross-linking of a donor stem assembled after the medium switch to an acceptor stem assembled prior to the medium switch (referred as new→old isotopologue dimers). These isotopologue dimers correspond to a *de novo* synthesized donor stem attached to the existing peptidoglycan, which provided the acceptor. The third type of dimers was generated by the cross-linking of two neo-synthesized stem peptides (new→new). The h1, h2, and h3 hybrid depicted in Fig. 4 were assigned to the neo-synthesized type of disaccharide-peptides since they contained unlabeled moieties implying their assembly after the medium switch. The absence of old→new dimers (see Supplementary data for MS/MS analyses) indicated that the peptide stems present in the existing peptidoglycan were not used as donors for cross-linking to neo-synthesized acceptors. For 4→3 cross-linked dimers, this result is expected since D,D-transpeptidases belonging to the PBP family use pentapeptide-containing donors, which are almost completely absent form mature peptidoglycan due to the hydrolysis of the D-Ala^4^-D-Ala^5^ amide bond by D,D-carboxypeptidases (See Introduction section and Fig. 1). Accordingly, subunits containing pentapeptide stems were not detected (Fig. 3 and Supplementary Table S1). For the 3→3 cross-linked Tri→Tetra dimer, tetrapeptide stems present in the existing peptidoglycan could potentially be used as donors by the YcbB L,D-transpeptidases although this was not observed. Thus, the absence of old→new 3→3 cross-linked Tri→Tetra dimer indicates that YcbB discriminates neo-synthesized tetrapeptide stems (used as donors) from existing tetrapeptide stems (used as acceptors) by an unknown mechanism that does not depend upon the structure of the disaccharide-peptide unit but upon the presence of the donor in a neo-synthesized glycan chain and of the acceptor in the existing peptidoglycan. The absence of old→new 3→3 cross-linked Tri→Tri dimers could be accounted for by the same mechanism.

**Figure 6.**
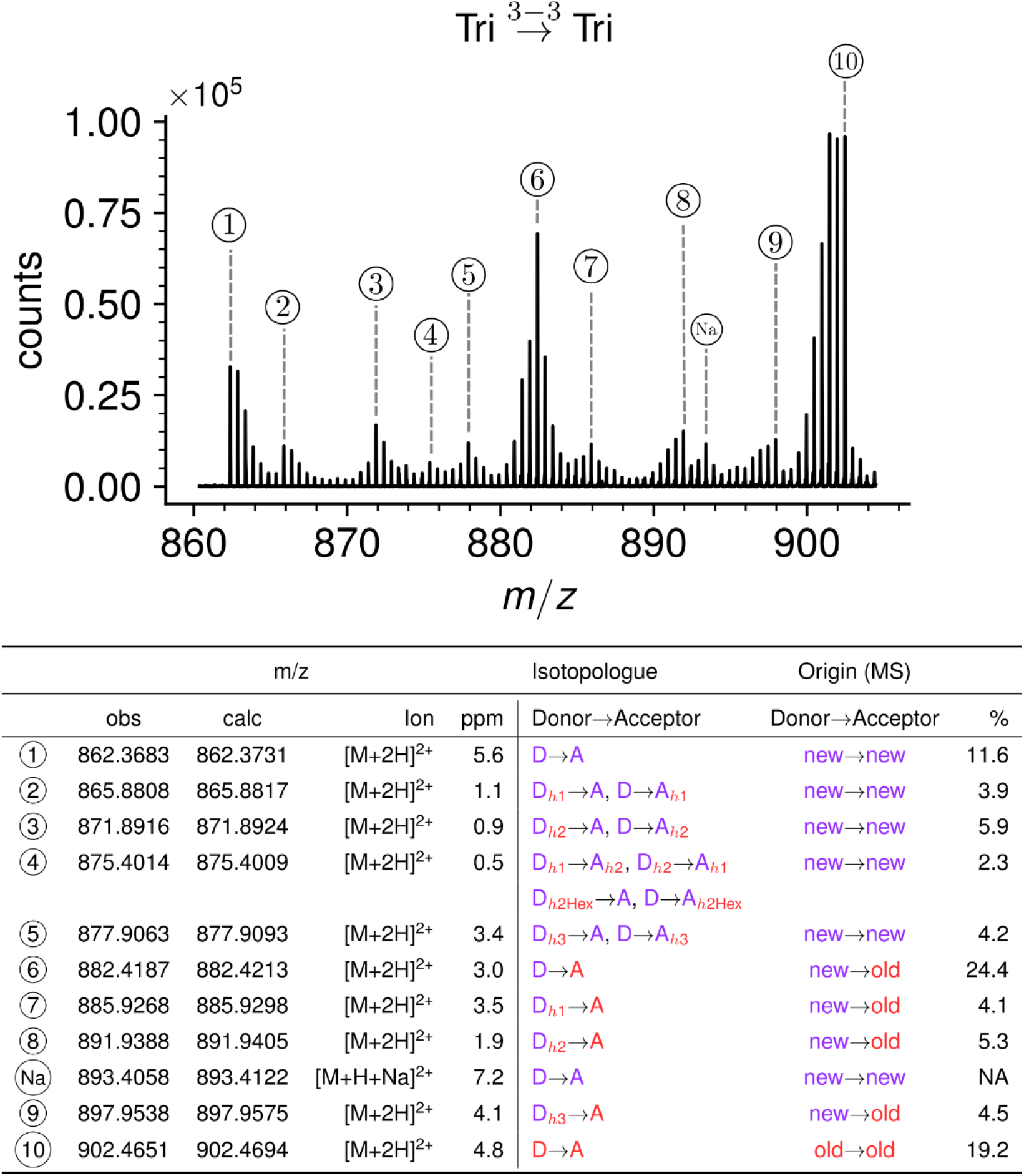
Structure of Tri→Tri isotopologues in the peptidoglycan of strain M1.4 grown in the absence of β-lactams. (**A**) Mass spectrum highlighting the 10 major isotopologues. (**B**) Structure of the 10 major isotopologues. The observed (obs) and calculated (cal) *m*/z values of the molecular ions (ion) and the difference in part per million (ppm) between these values are indicated. New (neo-synthesized) unlabeled moieties of isotopologues are indicated in purple. Old (existing) labeled moieties of isotopologues are indicated in red. h1, h2, and h3 (in red) refer to recycled moieties originating from existing labeled peptidoglycan (h1 and h2) or from the existing (labeled) UDP-MurNAc-pentapeptide pool as described in Fig. 4. The origin of the donor and acceptor participating in the cross-linking reaction, new (neo-synthesized) or old (existing in the cell wall) are indicated in purple and red, respectively. The relative abundance of the isotopologues (%) was deduced from the relative intensity of peaks corresponding to [M+2H]^2+^ ions as labeled in A. **Abbreviations:** h1, hybrid containing a recycled glucosamine moiety; h2, hybrid containing a recycled tripeptide moiety; h3, hybrid containing a labeled MurNAc-tripeptide moiety originating from the existing UDP-MurNAc-pentapeptide pool. NA, not applicable, as the sodium adduct was not taken into consideration.

Kinetic analyses revealed that the Tri→Tri and Tetra→Tri dimers formed by PBPs and YcbB, respectively, were almost exclusively of the new→old type (Fig. 7). In contrast, Tri→Tetra and Tetra→Tetra dimers formed by these enzymes were mixtures of new→new and new→old types of dimers. Strikingly, the kinetics of formation of dimers containing a tripeptide (Tri→Tri and Tetra→Tri) or a tetrapeptide (Tri→Tetra and Tetra→Tetra) in the acceptor position were similar for both types of transpeptidases. In contrast, distinct kinetics were observed between dimers containing a stem tripeptide or tetrapeptide in the acceptor position irrespective of the transpeptidase responsible for their formation. Thus, the mode of insertion of peptidoglycan subunits depended upon the structure of the acceptor stem (tripeptide *versus* tetrapeptide) rather than upon the type of transpeptidase (PBPs *versus* YcbB).

**Figure 7.**
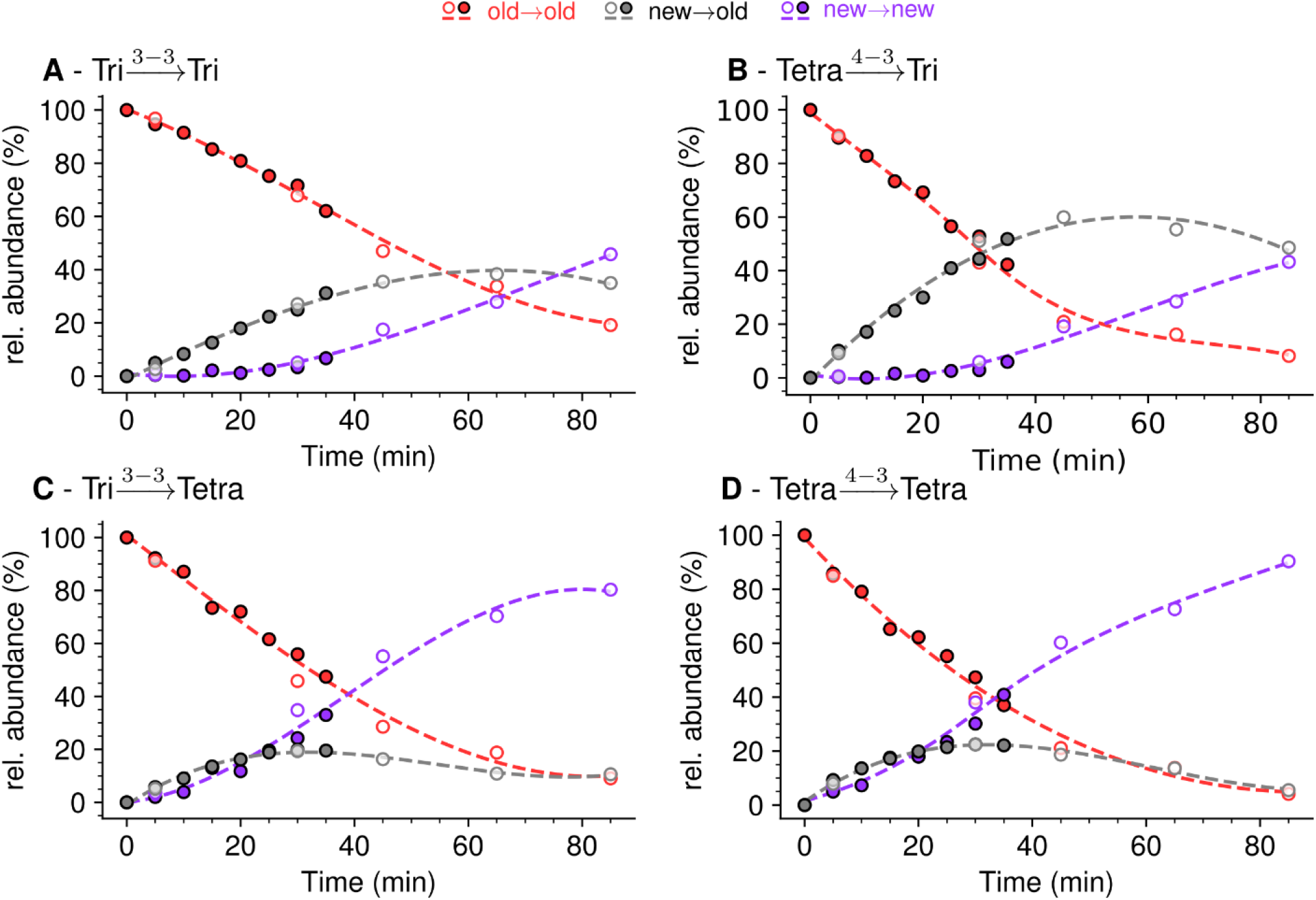
Mode of cross-linking of *de novo* synthesized peptidoglycan subunits in *E. coli* M1.5 grown in the absence of β-lactam. For each dimer (**A, B, C**, and **D**), the relative abundance (rel. abundance) was calculated for isotopologues originating from cross-liking of (i) two existing stems (old→old; red), (ii) a new donor stem (*i*.*e*. a stem synthesized after the medium switch) and an existing acceptor stem (new→old; grey), and (iii) two new stems (new→new; purple). The calculations take into account the contribution of *de novo* synthesized stems originating from recycled moieties and from the labeled UDP-MurNAc-pentapeptide pool as described in Figure 6 and in Supplementary Fig. S7, S8, and S9 for the Tri→Tri, Tetra→Tri, Tri→Tetra, and Tetra→Tetra dimers, respectively. The open and closed symbols correspond to data from two independent experiments.

### Mode of insertion of peptidoglycan subunits in the absence of any L,D-transpeptidase

Kinetic analyses of the insertion of peptidoglycan subunits was analyzed in *E. coli* Δ6*ldts*, a derivative of strain BW25113 that did not harbor any of the six *E. coli* L,D-transpeptidase genes. The absence of the corresponding enzymes greatly simplifies both the metabolic scheme for peptidoglycan cross-linking (Supplementary Fig. S10) and the muropeptide *rp*HPLC profile (Fig. 8). This profile contains only two main peaks, corresponding to the tetrapeptide monomer and the Tetra→Tetra dimer due to the absence of the formation of dimers containing 3→3 cross-links (L,D-transpeptidase activity) and of monomers or dimers containing tripeptide stems (L,D-carboxypeptidase activity). The medium switch led to the accumulation of the unlabeled tetrapeptide monomer and of hybrids h1, h2, and h3 (Fig. 8). The majority of the Tetra→Tetra dimers of the Δ6*ldts* strain were hybrids containing an unlabeled (*de novo* synthesized) tetrapeptide stem at the donor position and a labeled tetrapeptide stem (existing peptidoglycan) at the acceptor position. This pattern mainly fits the one-at-a time mode of insertion of newly synthesized glycan strands (Supplementary Fig. S2).

**Figure 8.**
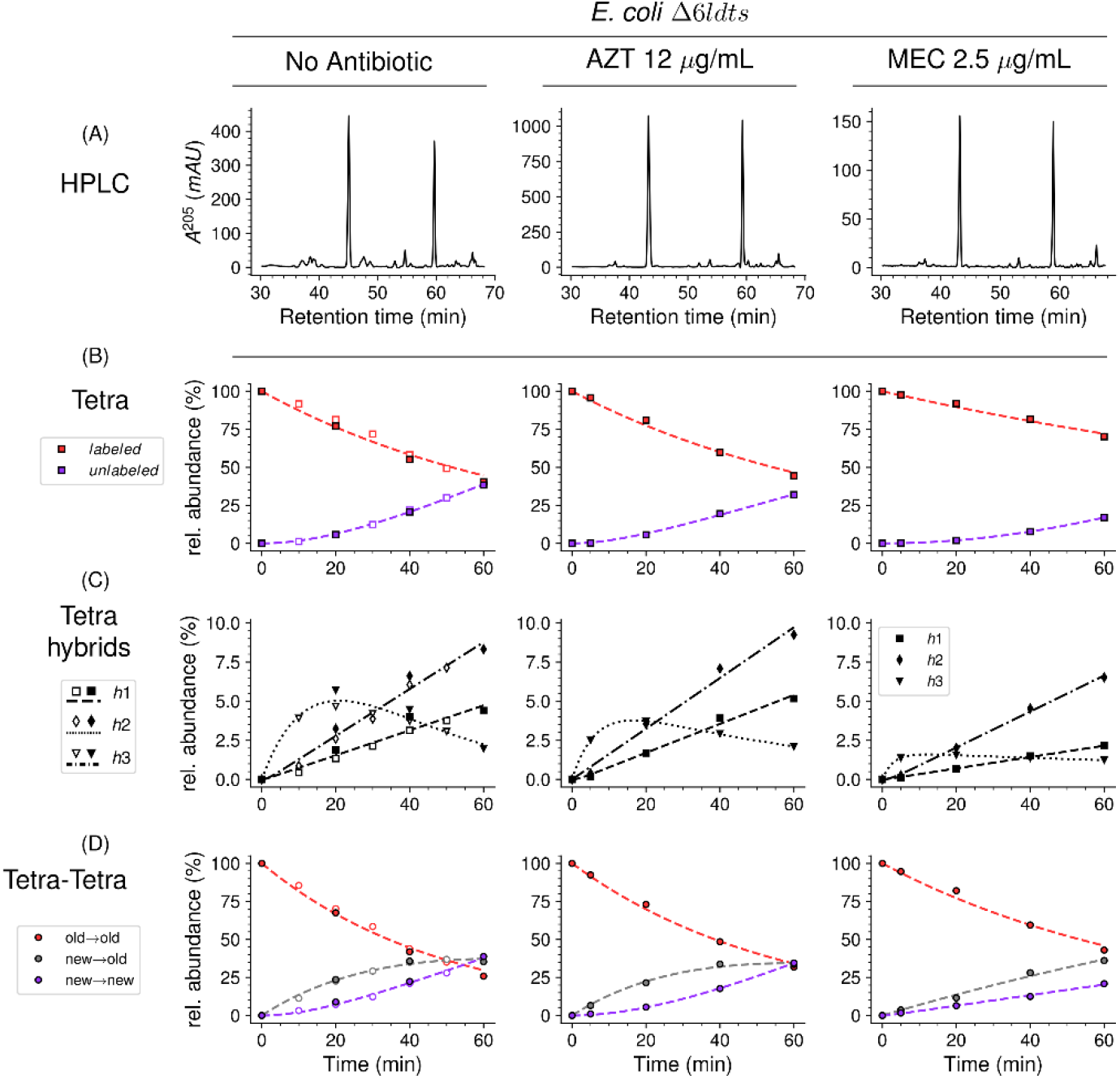
Mode of insertion of peptidoglycan subunits in the absence of L,D-transpeptidases and impact of the inhibition of PBPs by aztreonam and mecillinam. (**A**) Muropeptide *rp*HPLC profile of PG from *E. coli* strain Δ6*ldt* grown in the absence of β-lactam or in the presence of aztreonam (AZT) or mecillinam (MEC). Kinetic data were collected for fully labeled (red), fully unlabeled (purple) tetrapeptide monomers (Tetra) (**B**) and hybrids (**C**) defined in Supplementary Fig. S9. For the Tetra→Tetra dimer (**D**), the relative abundance (rel. abundance) was calculated for isotopologues originating from cross-liking of (i) two existing stems (old→old; red), (ii) a new donor stem (*i*.*e*. a stem synthesized after the medium switch) and an existing acceptor stem (new→old; grey), and (iii) two new stems (new→new; purple). The calculations take into account the contribution of *de novo* synthesized stems originating from recycled moieties and from the labeled UDP-MurNAc-pentapeptide pool. The open and closed symbols correspond to data from two independent experiments.

### Mode of insertion of peptidoglycan subunits into the septum and in the lateral wall

In *E. coli*, two β-lactams, mecillinam and aztreonam, specifically inhibit the synthesis of the lateral cell wall and of the septum by inactivating the D,D-transpeptidase activity of PBP2 and PBP3, respectively. Mecillinam (2.5 µg/mL) and aztreonam (12 µg/mL) were added to the labeled culture medium of strain Δ6*ldts* 5 min prior to the medium switch and to the unlabeled culture medium used to resuspend the bacteria at the medium switch. The kinetics of insertion of *de novo* synthesized subunits in the tetrapeptide monomers and in the Tetra→Tetra dimers were similar in the cultures containing aztreonam or no drug (Fig. 8). The pattern corresponded to the one-at-a time mode of insertion of newly synthesized glycan strands since the new-new dimer were not abundant at the beginning of the kinetics. In contrast, the new-new and new-old isotopologues dimers were both present in the cultures containing mecillinam indicating that formation of the septum involves insertion of multiple strands at a time.

### Mode of insertion of peptidoglycan subunits in conditions in which YcbB is the only functional transpeptidase

In the presence of ampicillin (16 µg/mL), the peptidoglycan of strain M1.5 was predominantly (*ca*. 99%) cross-linked by YcbB since the D,D-transpeptidase activity of PBPs was inhibited by the drug (Fig. 9 and Supplementary Table S1). Inhibition of D,D-carboxypeptidases belonging to the PBP family prevented full hydrolysis of D-Ala^5^ since pentapeptide stems were detected both in monomers and in the acceptor position of dimers. New→new rather than new→old isotopologues were predominantly detected among dimers containing a tetrapeptide or a pentapeptide stem in the acceptor position. Theses dimers originate from neo-synthesized glycan chains cross-linked to each other. In contrast, the Tri→Tetra dimers in the peptidoglycan of M1.5 grown in the absence of ampicillin mainly contained the new-old isotopologues (Fig. 7). This difference was not observed for the Tri→Tri dimers since the new-old isotopologues were prevalent for growth of M1.5 both in the presence or absence of ampicillin.

**Figure 9.**
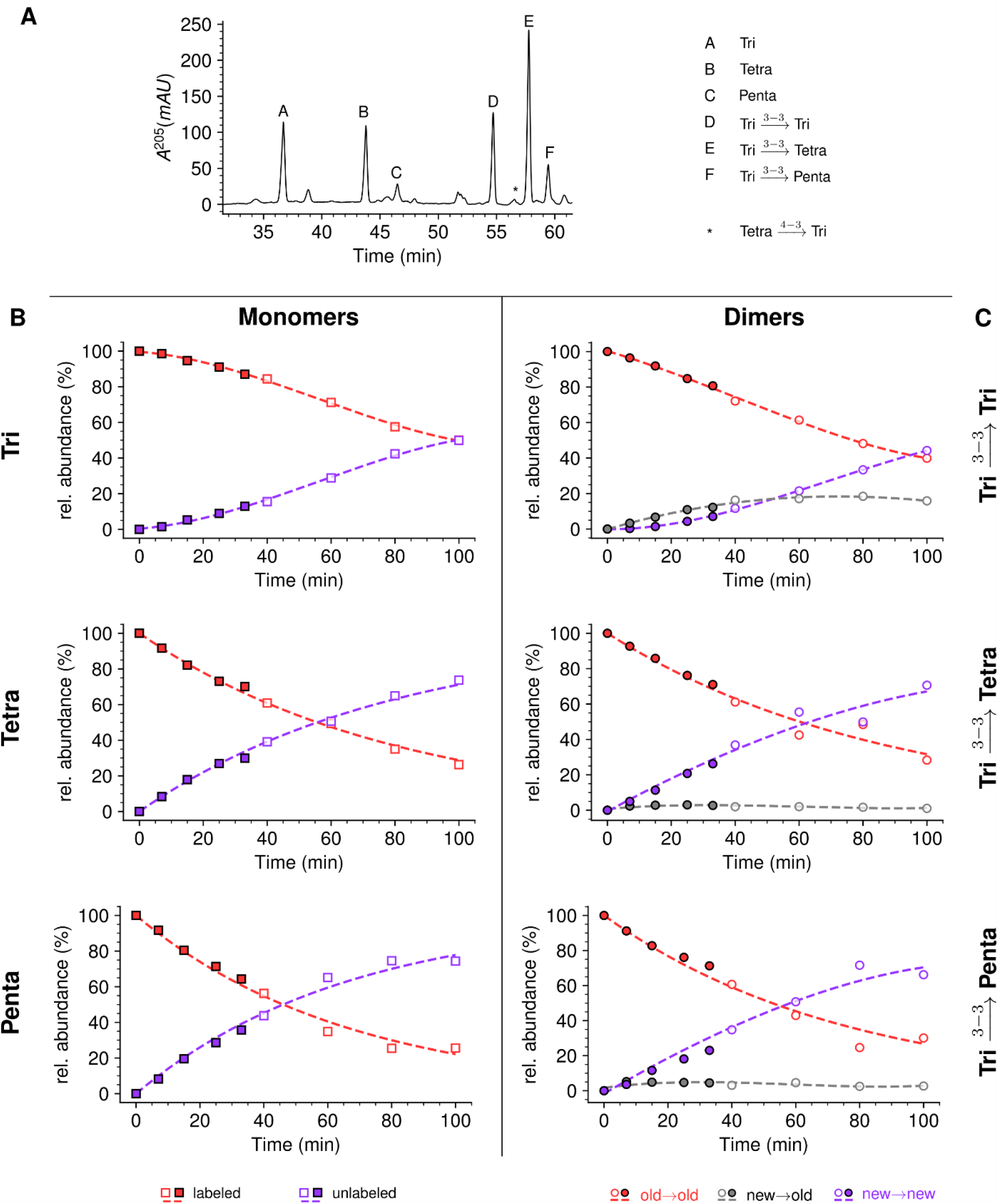
Mode of insertion of peptidoglycan in conditions in which YcbB is the only functional transpeptidase following inhibition of the D,D-transpeptidase activity of PBPs by ampicillin. (**A**) Muropeptide *rp*HPLC profile of PG from *E. coli* strain M1.5 grown in the presence of ampicillin (16 µg/mL). (**B**) Kinetic data were collected for neo-synthesized (new, red) and existing (old, purple) monomers. (**C**) Kinetic data were also collected for dimers originating from cross-liking of (i) two existing stems (old→old; red), (ii) a new donor stem (*i*.*e*. a stem synthesized after the medium switch) and an existing acceptor stem (new→old; grey), and (iii) two new stems (new→new; purple). The calculations take into account the contribution of *de novo* synthesized stems originating from recycled moieties and from the labeled UDP-MurNAc-pentapeptide pool. The open and closed symbols correspond to data from two independent experiments. Rel. abundance, relative abundance of isotopologues. Disaccharide-peptide subunits contained a tripeptide (Tri), a tetrapeptide (Tetra), or a pentapeptide (Penta) stem.

## DISCUSSION

Previous studies of the expansion of the peptidoglycan macromolecule relied on labeling with [^3^H] or [^14^C]DAP (de Jonge et al., 1989). Quantitatively, the number of glycan strands inserted at a time was estimated by determining the acceptor-to-donor radioactivity ratio (ADRR), *i*.*e*. the radioactivity ratio in the acceptor and donor positions of dimers (Supplementary Fig. S2) (Burman and Park, 1984). By analogy, we define here the acceptor-to-donor ratio in neo-synthesized stems (ADRNS). ADRR and ADRNS determinations rely on opposite labeling schemes involving pulse labeling with radioactive DAP *versus* incorporation of neo-synthesized (unlabeled) subunits into the fully labeled peptidoglycan, respectively. The latter strategy was chosen because of the ease of obtaining fully labeled cells with non-radioactive heavy isotopes. Switching the medium from labeled to unlabeled was chosen, rather than the unlabeled-to-labeled switch, since this facilitates the detection of neo-synthesized isotopologues present at a low abundance at early times after the medium switch. Indeed, the corresponding isotopic clusters do not appear in the region of the mass spectra containing the sodium adducts of the fully labeled isotopologues.

Pulse labeling with radioactive DAP suffers from four major limitations that are not encountered in our labeling strategy. First, optimization of radioactive DAP incorporation requires mutations that prevent endogenous DAP synthesis (*lysA*) and conversion of DAP to L-lysine (*dapF*) (Burman and Park, 1984, Glauner and Holtje, 1990, Goodell, 1985, Goodell and Schwarz, 1985) (Supplementary Fig. S1). Starvation prior to the addition of radiolabeled DAP was used to minimize the lag between the addition of radioactive DAP into the culture medium and its incorporation into peptidoglycan (Glauner and Holtje, 1990, Burman and Park, 1984). However, depleting the intracellular DAP pool compromised the integrity of the peptidoglycan layer (Burman et al., 1983b), affected cell morphology (Verwer and Nanninga, 1980), and reduced turnover (Burman et al., 1983b). Our labeling scheme is not affected by this confounding factors since it does not require any metabolic engineering. A second limitation of the DAP labeling procedures is their inability to discriminate between muropeptides originating from the existing peptidoglycan, from the recycled L-Ala-D-iGlu-DAP tripeptide, or from the existing UDP-MurNAc-pentapeptide pool, because all of them contain non-radioactive DAP. In contrast, our procedure easily discriminates between these three types of muropeptides since they correspond to different isotopologues, namely uniformly labeled subunits and hybrids h2 and h3, respectively (Fig. 4). The latter hybrids are likely to contribute to the lag phase observed for the incorporation of radioactive DAP into peptidoglycan in previous reports (Glauner and Holtje, 1990, Burman and Park, 1984). Accordingly, a lag phase was observed in our study when h2 and h3 hybrids containing [^13^C]- and [^15^N]-labeled DAP were excluded from the data set (Supplementary Fig. S11). A third limitation of the radioactive DAP labeling procedures for ADRR determination originates from the fact that they provide access only to the average radioactivity content of donor and acceptor stem peptides. Indeed, these procedures are based on purification of individual dimers, derivatization of their free amino group located in the acceptor stem, acid hydrolysis, and chromatography to separate DAP and derivatized DAP originating from the donor and acceptor stems, respectively (Burman et al., 1983a). Our procedure is richer in information since it provides direct access to the structure of dimers enabling identification of new→new, new→old, and old→new dimers based on differences in mass and fragmentation patterns. The fourth limitation stems from the fact that exploring peptidoglycan recycling by the radioactive DAP labeling procedures requires use of specific mutants and study designs. For example, Jacobs *et al*. used mutants deficient in the production of the AmpG and Opp permeases in which peptidoglycan fragments are not recycled in the cytoplasm but released in the culture medium (Jacobs et al., 1994). In contrast, our approach enables investigating peptidoglycan recycling and the mode of peptidoglycan expansion in a single experiment in the absence of any metabolic perturbation since it does not require altering the recycling pathway and is solely based on the determination of the structure of peptidoglycan fragments.

Strain BW25113 Δ6*ldt* offered the possibility to investigate the mode of insertion of new subunits by PBPs in the absence of any LDT. Aztreonam and mecillinam were used to specifically inhibit synthesis of septal and lateral peptidoglycan, respectively. In the presence of aztreonam, the ADRNS for side wall synthesis was 0.14 five minutes after the medium switch and increased to 0.20, 0.34, 0.50, and 0.64 at 20, 40, 60, and 80 minutes, respectively (Supplementary Fig. S12). At this time scale, existing disaccharide-peptide units are gradually replaced by neo-synthesized disaccharide-peptides accounting for the increase in new→new dimers and the increase in the ADRNS value. The extrapolation of the ADRNS at t=0 (time of the medium switch) is therefore relevant to the evaluation of the mode of insertion of new subunits into the peptidoglycan. The extrapolated ADRNS value of 0.08 (Supplementary Fig.

S12) was similar to the ADRR value of 0.10 reported by de Jonge and collaborators based on extrapolating data to a synchronized culture exclusively containing elongating cells (de Jonge et al., 1989). In the presence of mecillinam, the ADRNS values marginally increased over time, providing an extrapolated value of 0.31 for t=0. This value is lower than the ADRR of 0.64 obtained by extrapolating data to a synchronized culture exclusively containing constricting cells (de Jonge et al., 1989). The origin of this difference is unknown but could involve the fact that blocking cell elongation by mecillinam may not fully prevent incorporation of newly synthesized subunits into the peptidoglycan since this drug does not prevent the polymerization of glycan chains by the elongasome (Uehara and Park, 2008). Together, our results are in agreement with previous analyzes (de Jonge et al., 1989) showing single-strand insertion of neo-synthesized peptidoglycan into the side wall *versus* insertion of multiple strands for septal peptidoglycan synthesis. In the absence of mecillinam and aztreonam, the ADRNS value (0.14) was intermediate between the values observed in the presence of aztreonam (0.08) and mecillinam (0.31), as qualitatively expected for the combined contributions of cross-links located in the side wall and in the septum and the lower contribution of the latter to the cell surface.

In the absence of β-lactam, *E. coli* M1.5 relied both on the D,D-transpeptidase activity of PBPs and the L,D-transpeptidase activity of YcbB for peptidoglycan cross-linking, leading to a high peptidoglycan content in both 4→3 (56%) and 3→3 (44%) cross-linked dimers (Supplementary Table S1). The proportions of dimers containing two neo-synthesized stems (new→new) or one neo-synthesized and one existing stem (new→old) were similar for the two types of cross-links (Fig. 7). Thus, the D,D-transpeptidase activity of PBPs and the L,D-transpeptidase activity of YcbB were involved in similar modes of insertion of new strands into the existing peptidoglycan of strain M1.5. The ADRNS values were in the order of 0.17 and 0.22 for 4→3 and 3→3 cross-links, respectively. These values indicate that the neo-synthesized glycan strands were predominantly but not exclusively cross-linked to the existing peptidoglycan for both types of transpeptidation reactions. Peptidoglycan polymerization might involve single-strand insertion into the side-wall and insertion of multiple strands into the septum, as discussed above for strain Δ6*ldt*. The striking similarities between dimers generated by PBPs and YcbB raise the possibility that the mode of insertion of new subunits is not determined by the transpeptidases but by other components of the peptidoglycan polymerization complexes, such as the scaffolding proteins.

The peptidoglycan of strain M1.5 grown in the presence of ampicillin was mainly (99%) cross-linked by YcbB following the inhibition of the D,D-transpeptidase activity of PBPs (Fig. 9). Strikingly, the Tri→Tetra and Tri→Penta dimers were almost exclusively of the new→new type. This pattern was not found in strains M1.5 and Δ6*ldt* grown in the absence of β-lactam (Fig. 7 and 8). The prevalence of new→new dimers may be a consequence of the inhibition of the transpeptidase activity of PBPs by ampicillin that results in the uncoupling of the cross-linking and transglycosylation reactions (Uehara and Park, 2008, Cho et al., 2014). This in turn results in the accumulation of linear glycan strands that are eventually cross-linked to each other by YcbB because they are not effectively degraded due to a mutation present in M1.5 that prevents production of lytic transglycosylase Slt70 (Voedts et al., 2021, Cho et al., 2014). The resulting new→new cross-linked material was almost exclusively connected to the existing peptidoglycan by the formation of Tri→Tri dimers because they were the only new→old dimers detected in the peptidoglycan. This implies that the acceptors of the cross-linking reactions in the existing peptidoglycan were tripeptides, whereas existing tetrapeptides also acted as acceptors in the absence of ampicillin (Fig. 8). The nearly exclusive use of existing tripeptides as acceptors in the presence of ampicillin could be accounted for by the fact that these tripeptides originate from cleavage of 3→3 cross-links liberating the tripeptide stems present in the donor position of the existing dimers. This model implies a cooperation between YcbB and an endopeptidase that provides the tripeptide acceptor substrate of the reaction by cleaving existing 3→3 cross-links (Fig. 10). This enzyme could be MepK since this endopeptidase preferentially cleaves 3→3 cross-links *in vitro*, is essential for bacterial growth only in strains overproducing YcbB, and is not inhibited by ampicillin (Voedts et al., 2021). According to this model, the reactions catalyzed by MepK and YcbB are concerted rather than sequential. This mode of expansion of the peptidoglycan network differs from the ‘make-before-break’ models that postulate that neo-synthesized glycan strands are cross-linked to free stem peptides of the existing peptidoglycan prior to the cleavage of existing cross links. In the absence of ampicillin, the hydrolysis of 4→3 cross-links by endopeptidases belonging to the PBP family could generate tetrapeptide acceptors in addition to the tripeptide acceptors exclusively used in the presence of the drug.

**Figure 10.**
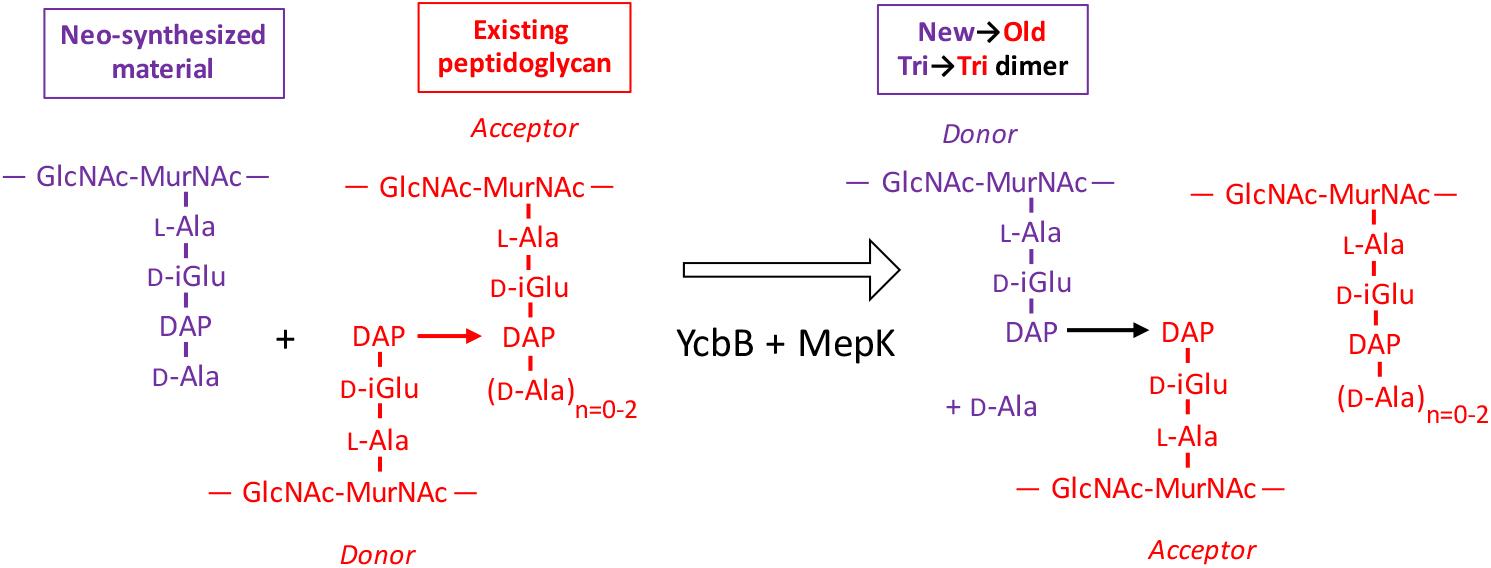
Model for the cross-linking of neo-synthesized material to existing peptidoglycan in strain M1.5 grown in the presence of ampicillin. The formation of New→Old cross-links results from cleavage of existing 3→3 cross-links by endopeptidase MepK thereby providing the tripeptide acceptor of the cross-linking reaction catalyzed by YcbB. Please note that the stem tripeptide in the donor position of dimers in the existing peptidoglycan is used as an acceptor in the neo-synthesized cross-links. This accounts for the fact that New→Old dimers are exclusively of the Tri→Tri type. The acceptor stem in the existing peptidoglycan may harbor 0, 1, or 2 D-Ala residue (n=0-2).

Cho et al. proposed that the antibacterial activity of β-lactams does not merely rely on a loss of function, *i*.*e*. the inactivation of the D,D-transpeptidase activity of PBPs by acylation of their catalytic Ser residue, but by inducing a toxic malfunctioning of the peptidoglycan biosynthesis machinery (Cho et al., 2014). This involves uncoupling of transglycosylation and transpeptidation, accumulation of polymerized glycan chains, and their degradation by lytic glycosyltransferase Slt70 (Banzhaf et al., 2012, Cho et al., 2014, Uehara and Park, 2008). The resulting futile cycle depletes resources and prevent bacterial growth. Exposure of Δ*sltY* cells to mecillinam was reported to result in the polymerization of an ‘aberrant peptidoglycan’ enriched in 3→3 cross-links. In comparison to this previous report (Cho et al., 2014), our analysis revealed that overproduction of YcbB results in substantially different modifications of peptidoglycan metabolism. First, M1.5 grown in the absence of β-lactam released a high proportion of its peptidoglycan per generation (56%) (Supplementary Fig. S13). This indicates that formation of 3→3 cross-links by YcbB is sufficient to stimulate peptidoglycan degradation in the absence of β-lactams. Under these conditions, Slt70 was not the main lytic enzyme since this activity is impaired in M1.5 (Voedts et al., 2021). Exposure to β-lactam had an additional stimulating effect on peptidoglycan degradation that reached 78% per generation for M1.5 grown in the presence of ampicillin (Supplementary Fig. S13). Second, overproduction of YcbB enabled growth of M1.5 in the presence of β-lactams indicating that the ‘aberrant peptidoglycan’ described by Cho *et al*. (Cho et al., 2014) can in fact sustain bacterial growth. This involved an unprecedented mode of peptidoglycan synthesis involving polymerization of linear glycan chains, their cross-linking by YcbB, and the incorporation of the resulting neo-synthesized material into the existing peptidoglycan by the concerted action of L,D-transpeptidase YcbB and of β-lactam-insensitive endopeptidase MepK.

## CONCLUSIONS

In conclusion, we report an innovative method for exploring peptidoglycan metabolism that does not depend upon metabolic engineering for labeling and is therefore applicable to various genetic backgrounds. The choice of uniform labeling with ^13^C and ^15^N enabled us investigating peptidoglycan metabolism at a very fine level of detail since most isotopologues predicted to occur according to known recycling and biosynthesis pathways were detected and kinetically characterized. In addition, high resolution MS and MS/MS identified the donors and acceptors that participated in individual cross-linking reactions rather than the average composition of dimers. This revealed an unexpected diversity in the modes of insertion of glycan strands into the expanding peptidoglycan macromolecule in the presence and absence of β-lactams. In particular, our results suggest that the reactions catalyzed by YcbB and by 3→3 specific endopeptidases in the presence of β-lactams are concerted rather than sequential as proposed by the ‘make-before-break’ models of peptidoglycan expansion. As previously discussed (Cho et al., 2014), efforts in drug discovery should focus on molecules that induce a lethal malfunctioning of multiple cellular targets rather simple inhibitors This implies that candidate molecules should be tested for their mode of action. The mass data analysis pipeline described in this study should facilitate such screening efforts by providing specific signatures for molecules acting on peptidoglycan synthesis.

## MATERIALS AND METHODS

### Sample preparation and analyses

#### Strains

*E. coli* M1.5 (Hugonnet et al., 2016, Voedts et al., 2021) is a derivative of *E. coli* BW25113 (Datsenko and Wanner, 2000) harboring (i) plasmid pJEH12 for IPTG-inducible expression of the *ycbB* L,D-transpeptidase gene, (ii) deletions of the *erfK, ynhG, ycfS*, and *ybiS* L,D-transpeptidase genes, (iii) a 13-bp deletion lowering the translation level of the gene encoding the isoleucyl-tRNA synthase (IleRS) thereby increasing synthesis of the (p)ppGpp alarmone, and (iv) an insertion of IS*1* in gene *sltY* encoding lytic transglycosylase Slt70 thereby promoting growth in the presence of ampicillin in liquid media (Voedts et al., 2021). Upon induction of *ycbB* by IPTG, *E. coli* M1.5 expresses high-level resistance to ampicillin and ceftriaxone following production of YcbB. Deletion of the *erfK, ycfS*, and *ybiS* genes abolishes the anchoring of the Braun lipoprotein and simplifies the *rp*HPLC muropeptide profile since stem peptides substituted by the Lys-Arg moiety of the lipoprotein are absent (Magnet et al., 2007).

The *ampG* permease gene (Johnson et al., 2013) of *E. coli* BW25113 M1.5 was deleted using the two-step procedure described by Datzenko and Wanner (Datsenko and Wanner, 2000). The kanamycin resistance gene used for allelic exchange was amplified with oligonucleotides 5’-CCAAAAACTATCGTGCCAGC and 5’-AGAGTGACAACTGGGTGATG. *E. coli* Δ6*ldts* is a derivative of strain BW25113 obtained by deletion of the six L,D-transpeptidase genes present in *E. coli* (*erfK, ynhG, ycfS, ybiS, yafK*, and *ycbB*) (Magnet et al., 2007, Magnet et al., 2008, Monton Silva et al., 2018, More et al., 2019).

#### Growth conditions and determination of the generation time

*E. coli* strains were grown in M9 minimal medium composed of 2 mM MgSO_4_, 0.00025% FeSO_4_, 12.8 g/L Na_2_HPO_4_, 3 g/L KH_2_PO_4_, 0.5 g/L NaCl, 1 g/L NH_4_Cl, 1 µg/L thiamine, 1 g/L glucose (unlabeled M9 minimal medium). The labeled M9 minimal medium had the same composition except for the replacement of NH_4_Cl and glucose by [^15^N]NH_4_Cl and uniformly labeled [^13^C]glucose at the same concentrations (99% labeling for both compounds; Cambridge Isotope Laboratories). The growth media were sterilized by filtration (Millex-LG 0.2 µm, Merck Millipore).

Bacteria were grown on brain heart infusion (BHI) agar plates (Difco) to initiate the experiment with an isolated colony that was inoculated in 25 mL of labeled minimal medium. The pre-culture was incubated for 18 h at 37 °C with aeration (180 rpm). The entire pre-culture (25 mL) was inoculated into one liter of labeled M9 medium and the resulting culture was incubated until the optical density at 600 nm (OD_600_) reached 0.4. Bacteria were collected by centrifugation (8,200 x *g*, 7 min, 22 °C), resuspended in 1 liter of prewarmed (37 °C) unlabeled M9 medium, and incubation was continued in the same conditions. Samples (200 mL) were withdrawn at various times and bacteria were collected by centrifugation (17,000 x *g*, 5 min, 4 °C) and stored at -55 °C. The generation time was deduced from the inverse of the slope of semi-logarithmic plots (log of OD_600_ as a function of time).

#### Peptidoglycan preparation and purification of muropeptides

Peptidoglycan was extracted by the hot SDS procedure and treated with pronase and trypsin (Arbeloa et al., 2004). Muropeptides were solubilized by digestion with lysozyme and mutanolysin, reduced with NaBH_4_, and separated by *rp*HPLC on a C_18_ column (Hypersil GOLD aQ 250 × 4.6, 3 µm; ThermoFisher) at a flow rate of 0.7 mL/min. A linear gradient (0% to 100%) was applied between 11.1 min and 105.2 min at room temperature (buffer A: 0.1% TFA; buffer B: 0.1% TFA, 20% acetonitrile; v/v). The absorbance was monitored at 205 nm and peaks corresponding to the major monomers and dimers were individually collected, lyophilized, solubilized in 30 µL of water, and stored at -20 °C.

#### Calculation of the molar ratios of muropeptides

*rp*HPLC analysis of sacculi digested by muramidases provided relative areas for peaks detected by the absorbance at 205 nm. In order to evaluate the relative molar abundance of muropeptides we used the correction factors reported by Glauner that take into account the number of disaccharide units, amide bonds, 1,6-anhydro-ends, and Lys and Arg residues (Glauner, 1988). Since the relative amount of the muropeptides did not vary upon growth (*i*.*e*. between samples collected at various times following the medium switch), or between samples from independent experiments, (*i*.*e*. between biological repeats obtained for the same strain in the same growth conditions) an average molar composition of the sacculi was deduced from the *rp*HPLC chromatograms and the correction factors reported by Glauner (Glauner, 1988).

#### Mass spectrometry

For routine identification of muropeptides, samples were diluted (10 μl plus 50 μl of water) and 2 μl were injected into the mass spectrometer (Maxis II ETD, Bruker, France) at a flow rate of 0.1 mL/min (50% acetonitrile, 50% water; v/v acidified with 0.1% formic acid; v/v). Mass spectra were acquired in the positive mode with a capillary voltage of 3,500V, a pre-storage pulse of 18 μs, ion funnel 1 RF 300 Vp-p. The *m*/*z* scan range was from 300 to 1,850 at a speed of 2 Hz. Transfer time stepping was enabled with the following parameters: RF values 400 and 1,200 Vp-p, transfer times 30 μs and 90 μs, timing 50% and 50%. MS/MS spectra were obtained using a collision energy of 50 eV in the *m*/*z* range of 150 to 1,000 for muropeptide monomers and 150 to 2,000 for muropeptide dimers with isolation width of 1. The applied collision energy was varied between 77 to 100% of the 50 eV setting with timings of 33% and 67%, respectively.

The structural characterization of the labeled GlcNAc-MurNAc-tetrapeptide isotopologues reported in Table 1 was performed on a ThermoFisher Orbitrap Fusion Lumos instrument operated in nanospray mode. The 50% acetonitrile in water analyte solution was acidified to 1% formic acid and introduced in the instrument using conventional nanospray glass emitter tips. The source voltage was set to 3,000-3,500 V and the sample was introduced without pneumatic assistance. The ions under each one of the peaks that made the isotopic cluster were fragmented individually by tune-mode quadrupole-selection of the corresponding ions in a 0.4 *m*/*z*-width selection window centered on the peak apex. That narrow selection window proved to be selective enough to only fragment the ions under each isotopic cluster peak of interest (from most intense to less intense: *m*/*z* 986.52, *m*/*z* 985.51, *m*/*z* 984.51). The fragmentation was obtained for a collision-induced dissociation energy of 10 eV. The MS/MS data were acquired in 200-1,000 *m*/*z*-range spectra (at least 50 MS/MS scans were acquired). Raw MS/MS data files were converted to the standard mzML file format using ProteoWizard’s msconvert tool (Adusumilli and Mallick, 2017). The obtained data files were scrutinized using the mineXpert2 software program (Langella and Rusconi, 2021).

The software developments required to predict and analyze the labeled/unlabeled muropeptide ions isotopic clusters either in MS or MS/MS experiments are hosted at https://gitlab.com/kantundpeterpan/masseltof and published under a Free Software license.

## Funding

*Agence National de la Recherche* (ANR-16-CE11-0030-12, TransPepNMR). MS data were obtained in the *Plateau Technique de Spectrométrie de Masse Bio-organique* of the *Muséum National d’Histoire Naturelle*.

## Notes

### Competing Interest Statement

The authors have declared no competing interest.

