## Supplementary Data for "Heavy isotope labeling and mass spectrometry reveal unexpected remodeling of bacterial cell wall expansion in response to drugs"


Tri - all heavy . . . . . 2
Tri - h1 . . . . . 4
Tri - h2 . . . . . 6
Tri - h3 . . . . . 8
Tetra - all heavy . . . . . 10
Tetra - h1 . . . . . 12
Tetra - h2 . . . . . 14
Tetra - h3 . . . . . 16
Tetra - hAla . . . . . 18
Tetra - h1h2 . . . . . 20
Tetra - h3Ala . . . . . 22
Additional monomer hybrids . . . . . 24
Tri-Tri - labeled . . . . . 25
Tri-Tri - unlabeled . . . . . 27
Tri-Tri - Donor all light . . . . . 29
Tetra-Tri - labeled . . . . . 31
Tetra-Tri - unlabeled . . . . . 33
Tetra-Tri - Donor all light . . . . . 35
Tri-Tetra - labeled . . . . . 37
Tri-Tetra - unlabeled . . . . . 39
Tri-Tetra - Donor all light . . . . . 41
Tetra-Tetra - labeled . . . . . 43
Tetra-Tetra - unlabeled . . . . . 45
Tetra-Tetra - Donor all light . . . . . 47

### Monomers

**Supplementary Data 1.1: Tandem mass spectrometry analysis of the labeled disaccharide-tripeptide.** The observed $m/z_{\mathrm{obs}}$ and calculated $m/z_{\mathrm{cal}}$ values of the parental ion, $[\mathrm{M+H}]^+$, as determined in the absence of fragmentation (MS1), are indicated. The $m/z_{\mathrm{calc}}$ value was used to select the ion for fragmentation. The difference between the $m/z_{\mathrm{obs}}$ and $m/z_{\mathrm{calc}}$ values is indicated in ppm. The GlcNAc residue is indicated in two moieties, glucosamine (GlcN) and the acetyl group (Ac), since these moieties are potentially differentially labeled. For the same reason, the reduced (Red) MurNAc residue is indicated in three moieties, reduced glucosamine ($\mathrm{GlcN^{Red}}$), the acetyl group (Ac), and the D-lactoyl group. Labeled and unlabeled moieties are represented in red and purple, respectively. The peaks corresponding to the list of fragments appearing in the table are highlighted by orange dots in the mass spectrum. In the mass spectrum, peaks differing by the loss of $H_20$ are connected by a dashed line. The tables only contain the $m/z$ values for the fragments containing $H_20$.
An interactive report of the $MS^2$ analysis is available in Supplementary File F1.1 .

| Precursor ion (MS1) | $m/z_{obs}$ | $m/z_{calc}$ | ppm |
|---|---|---|---|
| GlcN(-Ac)-$\mathrm{GlcN^{Red}}$(-Ac)-Lac-Ala-Glu-DAP | 911.475 | 911.475 | 0.3 |

| Product ion | $m/z_{obs}$ | $m/z_{calc}$ | ppm | Intensity (a.u.) | Isotopologue |
|---|---|---|---|---|---|
| $\mathrm{GlcN^{Red}}$(-Ac)-Lac-Ala-Glu-DAP | 699.371 | 699.371 | 0.7 | 7650 | all heavy |
| $\mathrm{GlcN^{Red}}$(-Ac)-Lac-Ala-Glu | 500.257 | 500.259 | 2.7 | 570 | all heavy |
| Lac-Ala-Glu-DAP | 485.254 | 485.253 | -2.7 | 220 | all heavy |
| Ala-Glu-DAP | 410.220 | 410.221 | 3.7 | 3226 | all heavy |
| $\mathrm{GlcN^{Red}}$(-Ac)-Lac-Ala | 365.201 | 365.202 | 2.5 | 4356 | all heavy |
| Glu-DAP | 335.177 | 335.177 | 0.3 | 9504 | all heavy |
| $\mathrm{GlcN^{Red}}$(-Ac)-Lac | 290.158 | 290.158 | -1.0 | 1835 | all heavy |
| $\mathrm{GlcN^{Red}}$(-Ac) | 215.127 | 215.127 | 0.5 | 229 | all heavy |
| GlcN(-Ac) | 213.110 | 213.111 | 3.7 | 108 | all heavy |
| DAP | 200.122 | 200.121 | -4.1 | 3273 | all heavy |

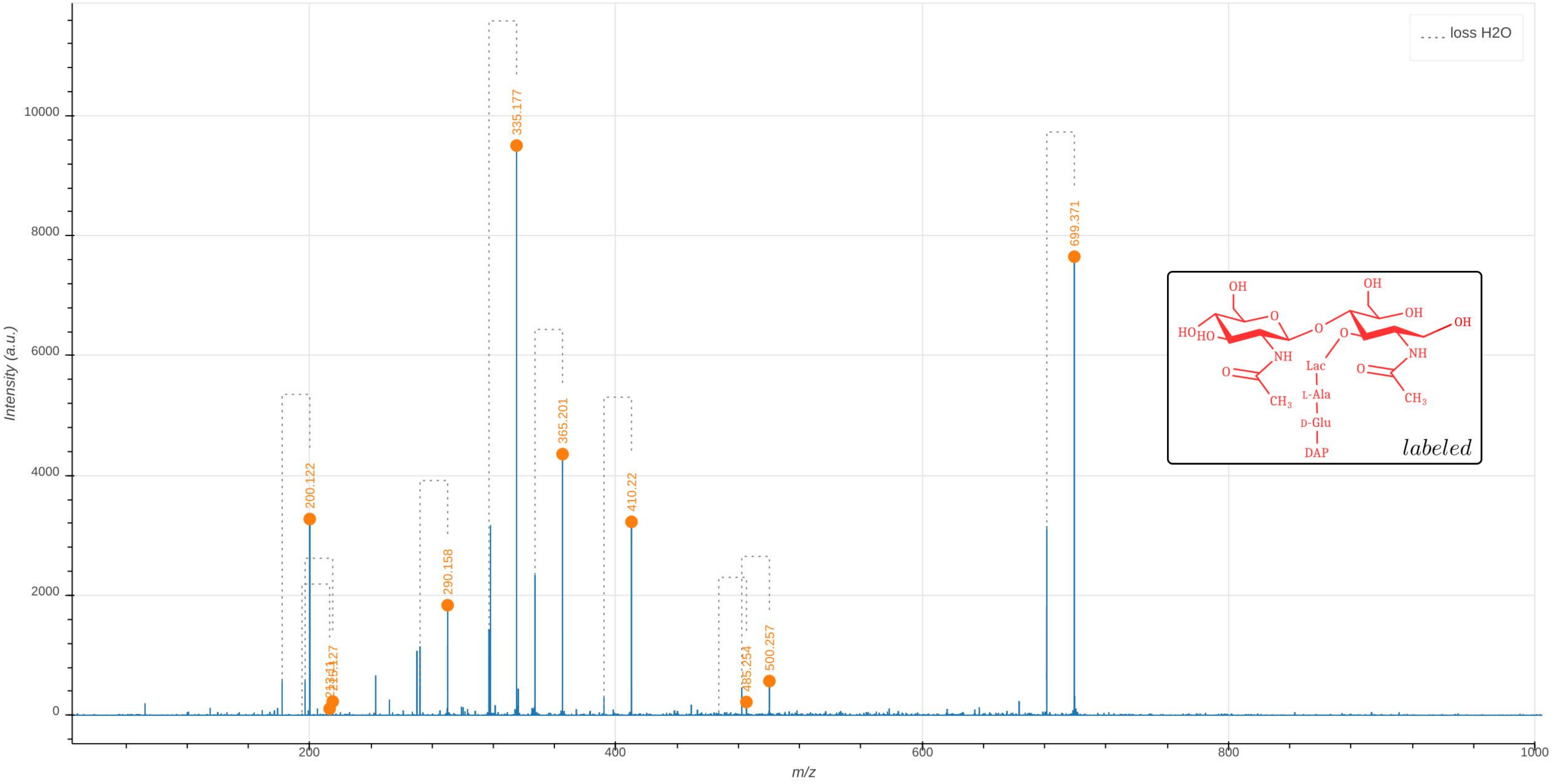

loss H2O
Intensity (a.u.)
m/z
0
2000
4000
6000
8000
10000
200
400
600
800
1000
200.122
215.127
290.158
335.177
365.201
410.22
485.254
500.257
699.371
OH
HO
HO
O
NH
CH3
Lac
L-Ala
D-Glu
DAP
labeled

**Supplementary Data 1.2: Tandem mass spectrometry analysis of the h1-type hybrid of the disaccharide-tripeptide.** The molecule is fully unlabeled except for one of the two glucosamine moieties present in GlcNAc or MurNAc leading to the presence of two isotopomers. The observed $m/z_{\mathrm{obs}}$ and calculated $m/z_{\mathrm{calc}}$ values of the parental ion, $[\mathrm{M{+}H}]^{+}$, as determined in the absence of fragmentation ($\mathrm{MS}^1$), are indicated. The $m/z_{\mathrm{cal}}$ value was used to select the ion for fragmentation. The difference between the $m/z_{\mathrm{obs}}$ and $m/z_{\mathrm{cal}}$ values is indicated in ppm. The GlcNAc residue is indicated in two moieties, glucosamine (GlcN) and the acetyl group (Ac), since these moieties are potentially differentially labeled. For the same reason, the reduced (Red) MurNAc residue is indicated in three moieties, reduced glucosamine ($\mathrm{GlcN}^{\mathrm{Red}}$), the acetyl group (Ac), and the D-lactoyl group. Labeled and unlabeled moieties are represented in red and purple, respectively. The fragmentation led to two series of fragments specific of the isotopomers containing the unlabeled glucosamine moiety in the GlcNAc (h1G) or $\mathrm{MurNAc}^{\mathrm{Red}}$ (h1M) residues, respectively (discriminatory fragments). The fragments specific of h1G and of h1M are highlighted by orange and green dots in the mass spectrum. The other fragments are common to the fragmentation patterns of h1G and of h1M. The corresponding peaks are highlighted by blue dots in the mass spectrum. The mass spectrum indicates the presence of both isotopomers as expected from the recycling and synthesis pathways since UDP-MurNAc exclusively derives from UDP-GlcNAc. In the mass spectrum, peaks differing by the loss of $H_20$ are connected by a dashed line. The tables only contain the $m/z$ values for the fragments containing $H_20$.An interactive report of the $\mathrm{MS}^2$ analysis is available in Supplementary File F1.2
.

| Precursor Ion ($\mathrm{MS}^1$) | $m/z_{obs}$ | $m/z_{calc}$ | ppm |
|---|---|---|---|
| GlcN(-Ac)-GlcN$^{Red}$(-Ac)-Lac-Ala-Glu-DAP | 878.392 | 878.396 | -4.0 |
| GlcN(-Ac)-GlcN$^{Red}$(-Ac)-Lac-Ala-Glu-DAP | 878.392 | 878.396 | -4.0 |

| Discriminatory product ion | $m/z_{obs}$ | $m/z_{calc}$ | ppm | Intensity (a.u.) | Isotopomer |
|---|---|---|---|---|---|
| GlcN$^{Red}$(-Ac)-Lac-Ala-Glu-DAP | 675.318 | 675.316 | -2.3 | 10563 | h1M |
| GlcN$^{Red}$(-Ac)-Lac-Ala-Glu-DAP | 668.299 | 668.299 | 0.6 | 8694 | h1G |
| GlcN$^{Red}$(-Ac)-Lac-Ala-Glu | 485.221 | 485.221 | 0.3 | 952 | h1M |
| GlcN$^{Red}$(-Ac)-Lac-Ala-Glu | 478.202 | 478.204 | 3.4 | 662 | h1G |
| GlcN$^{Red}$(-Ac)-Lac-Ala | 356.177 | 356.178 | 3.6 | 7236 | h1M |
| GlcN$^{Red}$(-Ac)-Lac-Ala | 349.161 | 349.161 | 1.1 | 6203 | h1G |
| GlcN$^{Red}$(-Ac)-Lac | 285.140 | 285.141 | 4.9 | 3223 | h1M |
| GlcN$^{Red}$(-Ac)-Lac | 278.123 | 278.124 | 4.0 | 3059 | h1G |
| GlcN$^{Red}$(-Ac) | 213.118 | 213.120 | 8.2 | 247 | h1M |
| GlcN(-Ac) | 211.106 | 211.104 | -6.8 | 238 | h1G |
| GlcN$^{Red}$(-Ac) | 206.103 | 206.103 | -0.5 | 148 | h1G |
| GlcN(-Ac) | 204.084 | 204.087 | 15.1 | 419 | h1M |

| Common product ion | $m/z_{obs}$ | $m/z_{calc}$ | ppm | Intensity (a.u.) |
|---|---|---|---|---|
| GlcN(-Ac)-GlcN$^{Red}$(-Ac)-Lac-Ala-Glu-DAP | 878.397 | 878.396 | -2.1 | 178 |
| GlcN(-Ac)-GlcN$^{Red}$(-Ac)-Lac-Ala-Glu-DAP | | | | |
| GlcN(-Ac)-GlcN$^{Red}$(-Ac)-Lac-Ala | 559.257 | 559.258 | 0.8 | 112 |
| GlcN(-Ac)-GlcN$^{Red}$(-Ac)-Lac-Ala | | | | |
| Lac-Ala-Glu-DAP | 463.203 | 463.204 | 1.8 | 613 |
| Ala-Glu-DAP | 391.182 | 391.183 | 2.0 | 9574 |
| Glu-DAP | 320.146 | 320.146 | 0.3 | 26374 |
| Ala-Glu | 201.085 | 201.088 | 13.5 | 227 |
| DAP | 191.103 | 191.103 | 0.6 | 6654 |
| Lac-Ala | 144.068 | 144.066 | -10.4 | 154 |

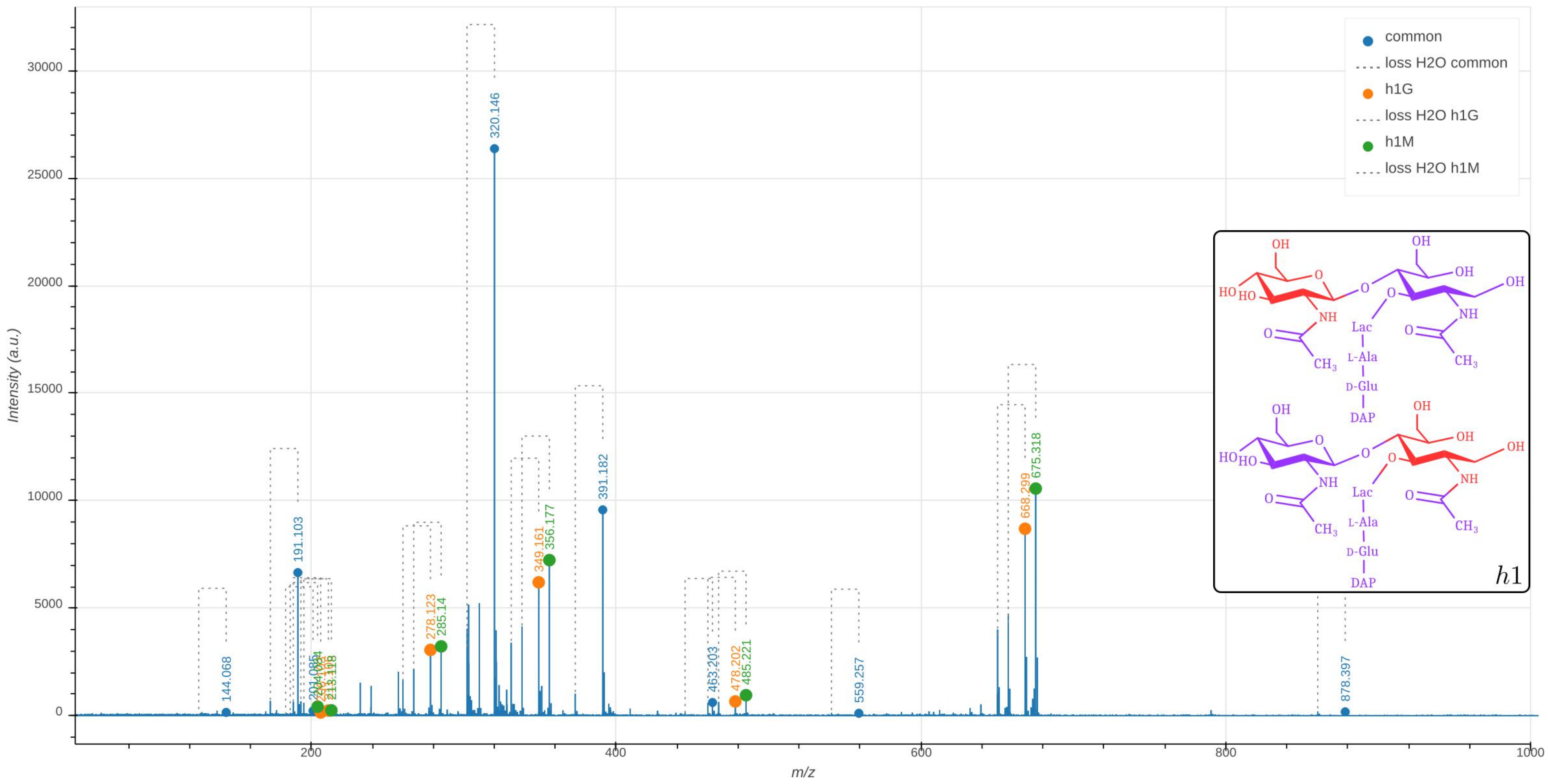

common
loss H2O common
h1G
loss H2O h1G
h1M
loss H2O h1M
Intensity (a.u.)
m/z
144.068
191.103
320.146
391.182
463.203
559.257
878.397
278.123
349.161
478.202
668.299
285.14
356.177
485.221
675.318
h1
Lac
L-Ala
D-Glu
DAP

**Supplementary Data 1.3: Tandem mass spectrometry analysis of the h2-type hybrid of the disaccharide-tripeptide.** The observed $m/z_{\text{obs}}$ and calculated $m/z_{\text{cal}}$ values of the parental ion, $[M+H]^+$, as determined in the absence of fragmentation (MS1), are indicated. The $m/z_{\text{calc}}$ value was used to select the ion for fragmentation. The difference between the $m/z_{\text{obs}}$ and $m/z_{\text{calc}}$ values is indicated in ppm. The GlcNAc residue is indicated in two moieties, glucosamine (GlcN) and the acetyl group (Ac), since these moieties are potentially differentially labeled. For the same reason, the reduced (Red) MurNAc residue is indicated in three moieties, reduced glucosamine ($\text{GlcN}^{\text{Red}}$), the acetyl group (Ac), and the D-lactoyl group. Labeled and unlabeled moieties are represented in red and purple, respectively. The peaks corresponding to the list of fragments appearing in the table are highlighted by orange dots in the mass spectrum. In the mass spectrum, peaks differing by the loss of $H_20$ are connected by a dashed line. The tables only contain the $m/z$ values for the fragments containing $H_20$.
An interactive report of the $MS^2$ analysis is available in Supplementary File F1.3 .

| Precursor ion (MS1) | $m/z_{obs}$ | $m/z_{calc}$ | ppm |
|---|---|---|---|
| GlcN(-Ac)-$\text{GlcN}^{\text{Red}}$(-Ac)-Lac-Ala-Glu-DAP | 890.412 | 890.417 | -5.7 |

| Product ion | $m/z_{obs}$ | $m/z_{calc}$ | ppm | Intensity (a.u.) | Isotopologue |
|---|---|---|---|---|---|
| $\text{GlcN}^{\text{Red}}$(-Ac)-Lac-Ala-Glu-DAP | 687.336 | 687.338 | 2.2 | 7465 | h2 |
| $\text{GlcN}^{\text{Red}}$(-Ac)-Lac-Ala-Glu | 488.223 | 488.225 | 3.1 | 404 | h2 |
| Lac-Ala-Glu-DAP | 482.239 | 482.242 | 7.1 | 296 | h2 |
| Ala-Glu-DAP | 410.219 | 410.221 | 5.5 | 3431 | h2 |
| $\text{GlcN}^{\text{Red}}$(-Ac)-Lac-Ala | 353.167 | 353.168 | 4.7 | 4771 | h2 |
| Glu-DAP | 335.178 | 335.177 | -2.4 | 9561 | h2 |
| $\text{GlcN}^{\text{Red}}$(-Ac)-Lac | 278.123 | 278.124 | 2.9 | 1727 | h2 |
| GlcN(-Ac) | 204.086 | 204.087 | 5.9 | 190 | h2 |
| DAP | 200.120 | 200.121 | 4.2 | 2800 | h2 |

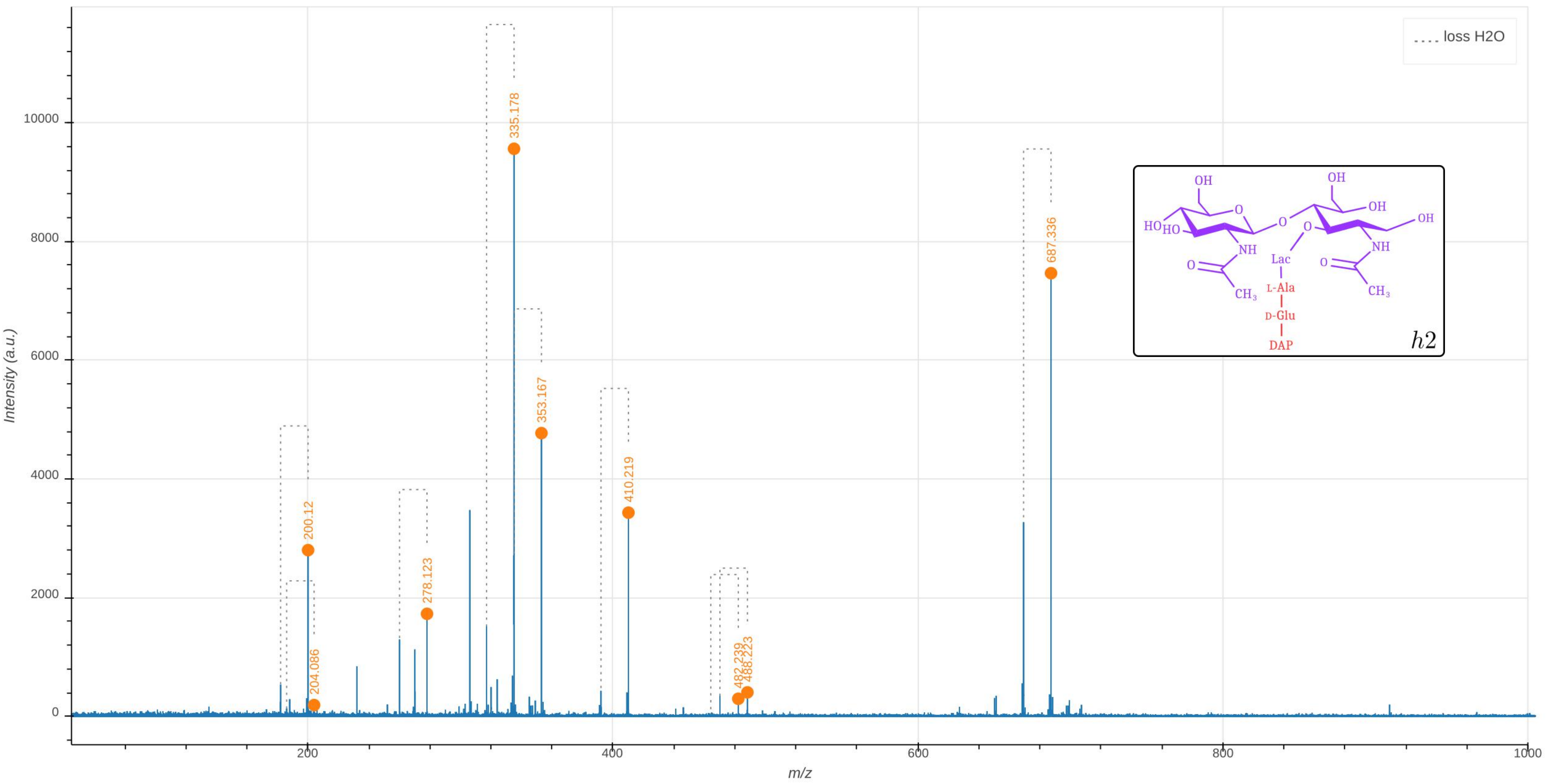

loss H2O
335.178
353.167
410.219
200.12
204.086
278.123
482.239
488.223
687.336
Intensity (a.u.)
m/z
0
2000
4000
6000
8000
10000
200
400
600
800
1000
OH
HO
HO
O
O
NH
O
CH3
OH
OH
OH
O
NH
O
CH3
Lac
L-Ala
D-Glu
DAP
h2

**Supplementary Data 1.4: Tandem mass spectrometry analysis of the h3-type hybrid of the disaccharide-tripeptide.** The observed $m/z_{\text{obs}}$ and calculated $m/z_{\text{cal}}$ values of the parental ion, $[M+H]^+$, as determined in the absence of fragmentation (MS1), are indicated. The $m/z_{\text{calc}}$ value was used to select the ion for fragmentation. The difference between the $m/z_{\text{obs}}$ and $m/z_{\text{calc}}$ values is indicated in ppm. The GlcNAc residue is indicated in two moieties, glucosamine (GlcN) and the acetyl group (Ac), since these moieties are potentially differentially labeled. For the same reason, the reduced (Red) MurNAc residue is indicated in three moieties, reduced glucosamine ($\text{GlcN}^{\text{Red}}$), the acetyl group (Ac), and the D-lactoyl group. Labeled and unlabeled moieties are represented in red and purple, respectively. The peaks corresponding to the list of fragments appearing in the table are highlighted by orange dots in the mass spectrum. In the mass spectrum, peaks differing by the loss of $H_20$ are connected by a dashed line. The tables only contain the $m/z$ values for the fragments containing $H_20$. The presence of the h3 isotopomer is expected since cells of *E. coli* contain the main PG precursor, UDP-MurNAc-pentapeptide in considerable amounts (about 2% compared to the total amount of dissaccharide peptides in the cell wall). An interactive report of the $MS^2$ analysis is available in Supplementary File F1.4 .

| Precursor ion (MS1) | $m/z_{obs}$ | $m/z_{calc}$ | ppm |
|---|---|---|---|
| GlcN(-Ac)-$\text{GlcN}^{\text{Red}}$(-Ac)-Lac-Ala-Glu-DAP | 902.447 | 902.451 | -3.8 |

| Product ion | $m/z_{obs}$ | $m/z_{calc}$ | ppm | Intensity (a.u.) | Isotopomer |
|---|---|---|---|---|---|
| $\text{GlcN}^{\text{Red}}$(-Ac)-Lac-Ala-Glu-DAP | 699.372 | 699.371 | -0.5 | 4005 | h3 |
| $\text{GlcN}^{\text{Red}}$(-Ac)-Lac-Ala-Glu | 500.259 | 500.259 | -0.8 | 329 | h3 |
| Lac-Ala-Glu-DAP | 485.251 | 485.253 | 3.1 | 308 | h3 |
| Ala-Glu-DAP | 410.219 | 410.221 | 5.1 | 1961 | h3 |
| $\text{GlcN}^{\text{Red}}$(-Ac)-Lac-Ala | 365.199 | 365.202 | 8.5 | 2535 | h3 |
| Glu-DAP | 335.178 | 335.177 | -2.9 | 5470 | h3 |
| $\text{GlcN}^{\text{Red}}$(-Ac)-Lac | 290.158 | 290.158 | 0.2 | 1047 | h3 |
| GlcN(-Ac) | 204.088 | 204.087 | -2.6 | 132 | h3 |
| DAP | 200.120 | 200.121 | 3.8 | 1331 | h3 |

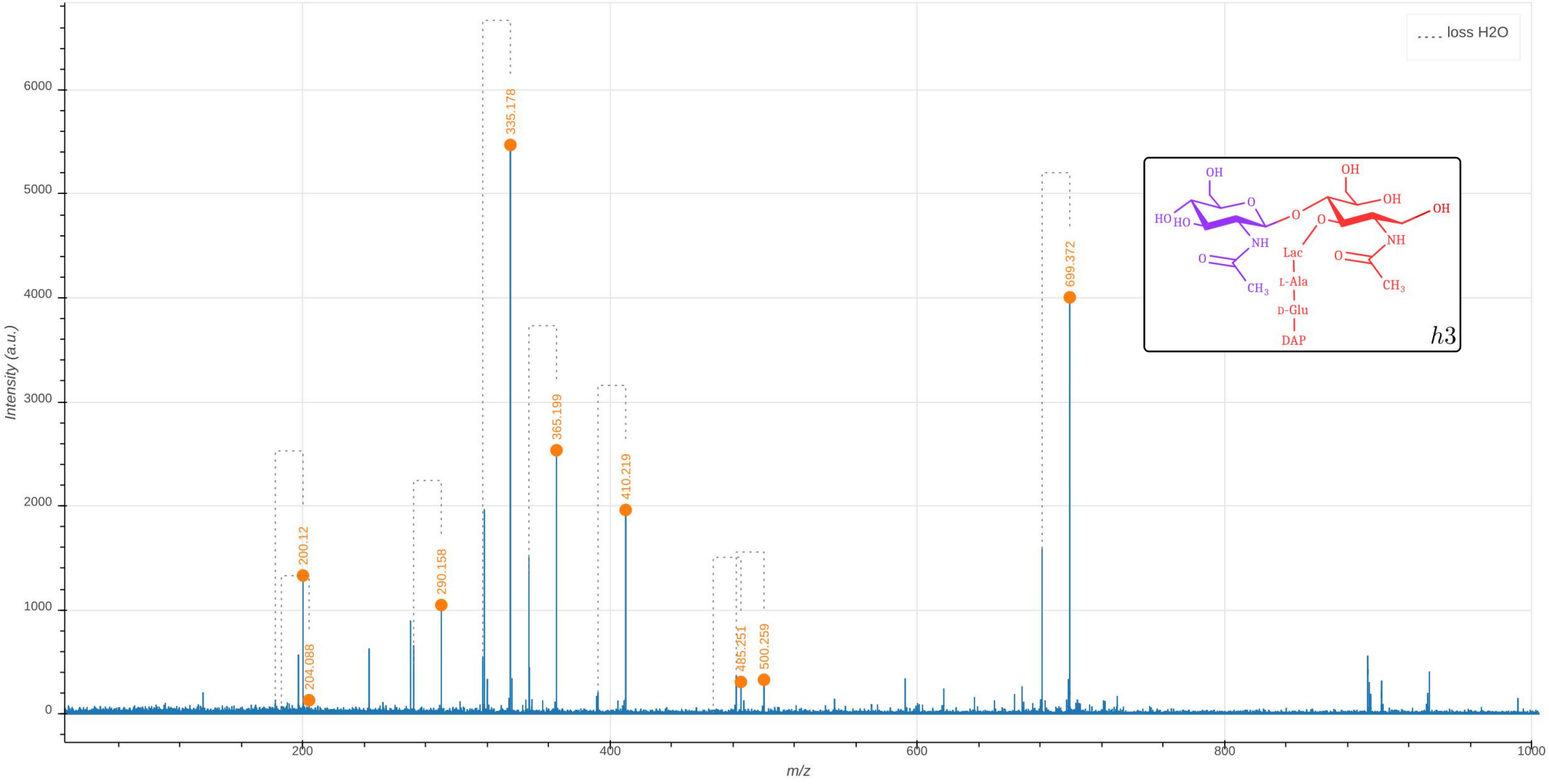

loss H2O
335.178
699.372
365.199
410.219
200.12
290.158
204.088
485.251
500.259
OH
O
HO
HO
NH
O
CH3
OH
OH
O
O
OH
NH
Lac
O
L-Ala
CH3
D-Glu
DAP
h3
Intensity (a.u.)
m/z
0
1000
2000
3000
4000
5000
6000
200
400
600
800
1000

**Supplementary Data 2.1: Supplementary Data D2.1 . Tandem mass spectrometry analysis of the labeled disaccharide-tetrapeptide.**

The observed $m/z_{\text{obs}}$ and calculated $m/z_{\text{cal}}$ values of the parental ion, $[M{+}H]^{+}$, as determined in the absence of fragmentation (MS1), are indicated. The $m/z_{\text{calc}}$ value was used to select the ion for fragmentation. The difference between the $m/z_{\text{obs}}$ and $m/z_{\text{calc}}$ values is indicated in ppm. The GlcNAc residue is indicated in two moieties, glucosamine (GlcN) and the acetyl group (Ac), since these moieties are potentially differentially labeled. For the same reason, the reduced (Red) MurNAc residue is indicated in three moieties, reduced glucosamine ($\text{GlcN}^{\text{Red}}$), the acetyl group (Ac), and the D-lactoyl group. Labeled and unlabeled moieties are represented in red and purple, respectively. The peaks corresponding to the list of fragments appearing in the table are highlighted by orange dots in the mass spectrum. In the mass spectrum, peaks differing by the loss of $H_20$ are connected by a dashed line. The tables only contain the $m/z$ values for the fragments containing $H_20$.

An interactive report of the MS2 analysis is available in Supplementary File F2.1.

| Precursor ion (MS1) | $m/z_{obs}$ | $m/z_{calc}$ | ppm |
|---|---|---|---|
| GlcN(-Ac)-$\text{GlcN}^{\text{Red}}$(-Ac)-Lac-Ala-Glu-DAP-Ala | 986.516 | 986.519 | -3.1 |

| Product ion | $m/z_{obs}$ | $m/z_{calc}$ | ppm | Intensity (a.u.) | Isotopologue |
|---|---|---|---|---|---|
| $\text{GlcN}^{\text{Red}}$(-Ac)-Lac-Ala-Glu-DAP-Ala | 774.423 | 774.416 | -9.3 | 5487 | all heavy |
| $\text{GlcN}^{\text{Red}}$(-Ac)-Lac-Ala-Glu-DAP | 681.364 | 681.361 | -5.1 | 494 | all heavy |
| Lac-Ala-Glu-DAP-Ala | 560.296 | 560.297 | 0.9 | 119 | all heavy |
| $\text{GlcN}^{\text{Red}}$(-Ac)-Lac-Ala-Glu | 500.259 | 500.259 | 0.0 | 194 | all heavy |
| Ala-Glu-DAP-Ala | 485.269 | 485.266 | -6.8 | 1147 | all heavy |
| Glu-DAP-Ala | 410.223 | 410.221 | -3.9 | 3963 | all heavy |
| Ala-Glu-DAP | 392.213 | 392.211 | -5.6 | 1512 | all heavy |
| $\text{GlcN}^{\text{Red}}$(-Ac)-Lac-Ala | 365.204 | 365.202 | -5.1 | 2434 | all heavy |
| Glu-DAP | 317.169 | 317.167 | -8.5 | 3861 | all heavy |
| $\text{GlcN}^{\text{Red}}$(-Ac)-Lac | 290.160 | 290.158 | -7.3 | 414 | all heavy |
| DAP-Ala | 275.167 | 275.165 | -6.2 | 3667 | all heavy |
| DAP | 182.112 | 182.110 | -8.1 | 611 | all heavy |

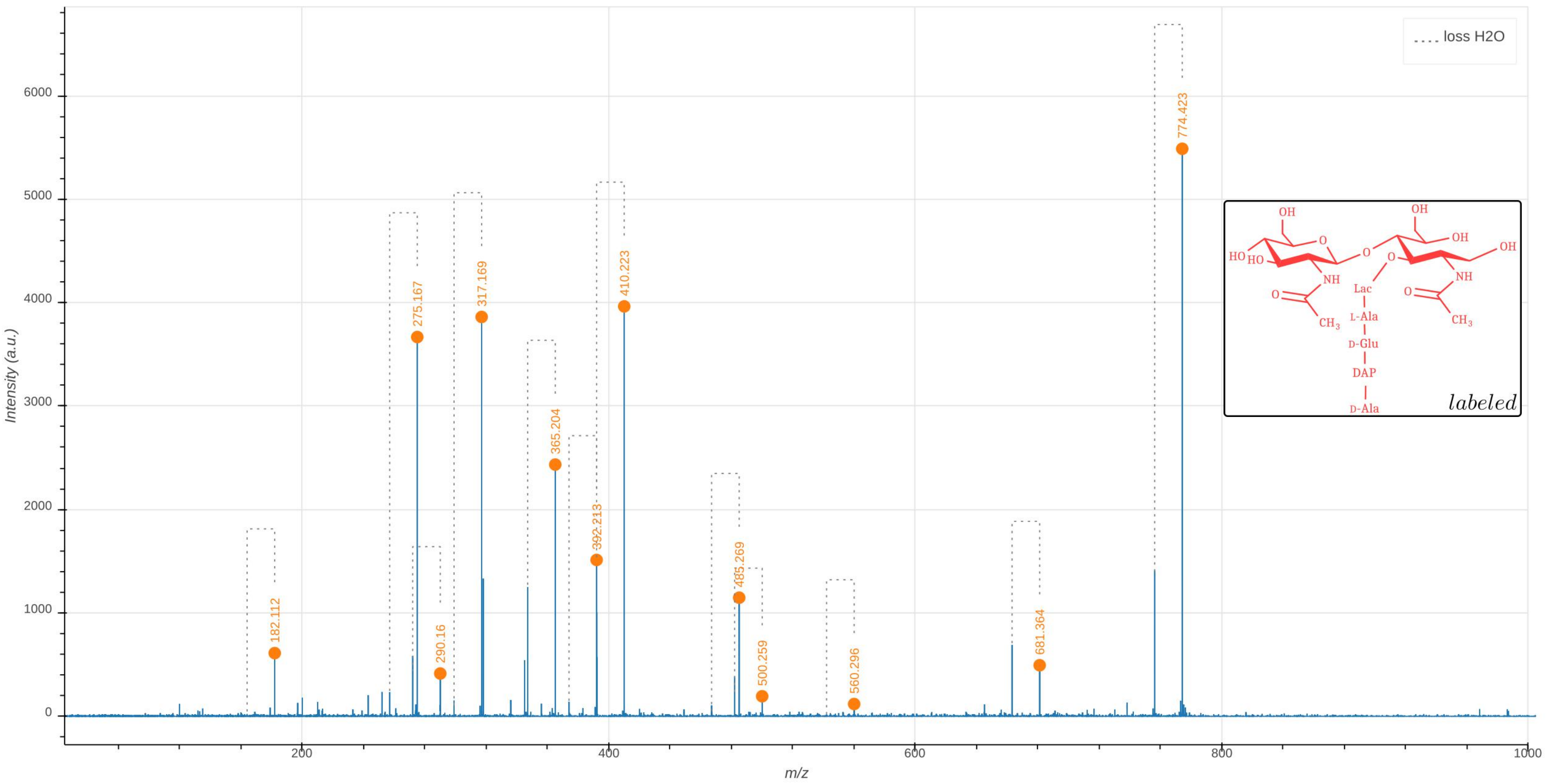

loss H2O
Intensity (a.u.)
m/z
0
1000
2000
3000
4000
5000
6000
200
400
600
800
1000
182.112
275.167
290.16
317.169
365.204
392.213
410.223
485.269
500.259
560.296
681.364
774.423
OH
HO
HO
O
NH
O
CH3
OH
OH
OH
O
O
NH
O
CH3
Lac
L-Ala
D-Glu
DAP
D-Ala
labeled

**Supplementary Data 2.2: Tandem mass spectrometry analysis of the h1-type hybrid of the disaccharide-tetrapeptide.** The molecule is fully unlabeled except for one of the two glucosamine moieties present in GlcNAc or MurNAc leading to the presence of two isotopomers. The observed $m/z_{\text{obs}}$ and calculated $m/z_{\text{calc}}$ values of the parental ion, $[M+H]^+$, as determined in the absence of fragmentation ($MS^1$), are indicated. The $m/z_{\text{cal}}$ value was used to select the ion for fragmentation. The difference between the $m/z_{\text{obs}}$ and $m/z_{\text{cal}}$ values is indicated in ppm. The GlcNAc residue is indicated in two moieties, glucosamine (GlcN) and the acetyl group (Ac), since these moieties are potentially differentially labeled. For the same reason, the reduced (Red) MurNAc residue is indicated in three moieties, reduced glucosamine ($\text{GlcN}^{\text{Red}}$), the acetyl group (Ac), and the D-lactoyl group. Labeled and unlabeled moieties are represented in red and purple, respectively. The fragmentation led to two series of fragments specific of the isotopomers containing the unlabeled glucosamine moiety in the GlcNAc (h1G) or $\text{MurNAc}^{\text{Red}}$ (h1M) residues, respectively (discriminatory fragments). The fragments specific of h1G and of h1M are highlighted by orange and green dots in the mass spectrum. The other fragments are common to the fragmentation patterns of h1G and of h1M. The corresponding peaks are highlighted by blue dots in the mass spectrum. The mass spectrum indicates the presence of both isotopomers as expected from the recycling and synthesis pathways since UDP-MurNAc exclusively derives from UDP-GlcNAc. In the mass spectrum, peaks differing by the loss of $H_20$ are connected by a dashed line. The tables only contain the $m/z$ values for the fragments containing $H_20$.An interactive report of the $MS^2$ analysis is available in Supplementary File F2.2 .

| Precursor ion (MS1) | $m/z_{obs}$ | $m/z_{calc}$ | ppm |
|---|---|---|---|
| GlcN(-Ac)-GlcN^Red(-Ac)-Lac-Ala-Glu-DAP-Ala | 949.429 | 949.433 | -4.0 |
| GlcN(-Ac)-GlcN^Red(-Ac)-Lac-Ala-Glu-DAP-Ala | 949.429 | 949.433 | -4.0 |

| Discriminatory product ions | $m/z_{obs}$ | $m/z_{calc}$ | ppm | Intensity (a.u.) | Isotopologue |
|---|---|---|---|---|---|
| GlcN^Red(-Ac)-Lac-Ala-Glu-DAP-Ala | 746.351 | 746.353 | 3.3 | 5188 | h1M |
| GlcN^Red(-Ac)-Lac-Ala-Glu-DAP-Ala | 739.333 | 739.336 | 4.7 | 3690 | h1G |
| GlcN^Red(-Ac)-Lac-Ala-Glu-DAP | 657.302 | 657.306 | 5.2 | 477 | h1M |
| GlcN^Red(-Ac)-Lac-Ala-Glu-DAP | 650.287 | 650.288 | 2.7 | 427 | h1G |
| GlcN^Red(-Ac)-Lac-Ala-Glu | 485.216 | 485.221 | 10.3 | 417 | h1M |
| GlcN^Red(-Ac)-Lac-Ala-Glu | 478.202 | 478.204 | 3.3 | 210 | h1G |
| GlcN^Red(-Ac)-Lac-Ala | 356.177 | 356.178 | 3.6 | 1850 | h1M |
| GlcN^Red(-Ac)-Lac-Ala | 349.158 | 349.161 | 8.1 | 2036 | h1G |
| GlcN^Red(-Ac)-Lac | 285.140 | 285.141 | 4.9 | 586 | h1M |
| GlcN^Red(-Ac)-Lac | 278.123 | 278.124 | 4.0 | 622 | h1G |
| GlcN(-Ac) | 204.086 | 204.087 | 7.2 | 137 | h1M |

| Common product ion | $m/z_{obs}$ | $m/z_{calc}$ | ppm | Intensity (a.u.) |
|---|---|---|---|---|
| GlcN(-Ac)-GlcN^Red(-Ac)-Lac-Ala-Glu-DAP-Ala | 949.433 | 949.433 | 0.1 | 241 |
| GlcN(-Ac)-GlcN^Red(-Ac)-Lac-Ala-Glu-DAP-Ala | | | | |
| Ala-Glu-DAP-Ala | 462.217 | 462.220 | 6.1 | 1977 |
| Glu-DAP-Ala | 391.180 | 391.183 | 6.2 | 7517 |
| Ala-Glu-DAP | 373.172 | 373.172 | 1.7 | 2756 |
| Glu-DAP | 302.135 | 302.135 | 0.9 | 7122 |
| DAP-Ala | 262.138 | 262.140 | 9.0 | 5564 |
| DAP | 173.093 | 173.093 | 0.3 | 796 |

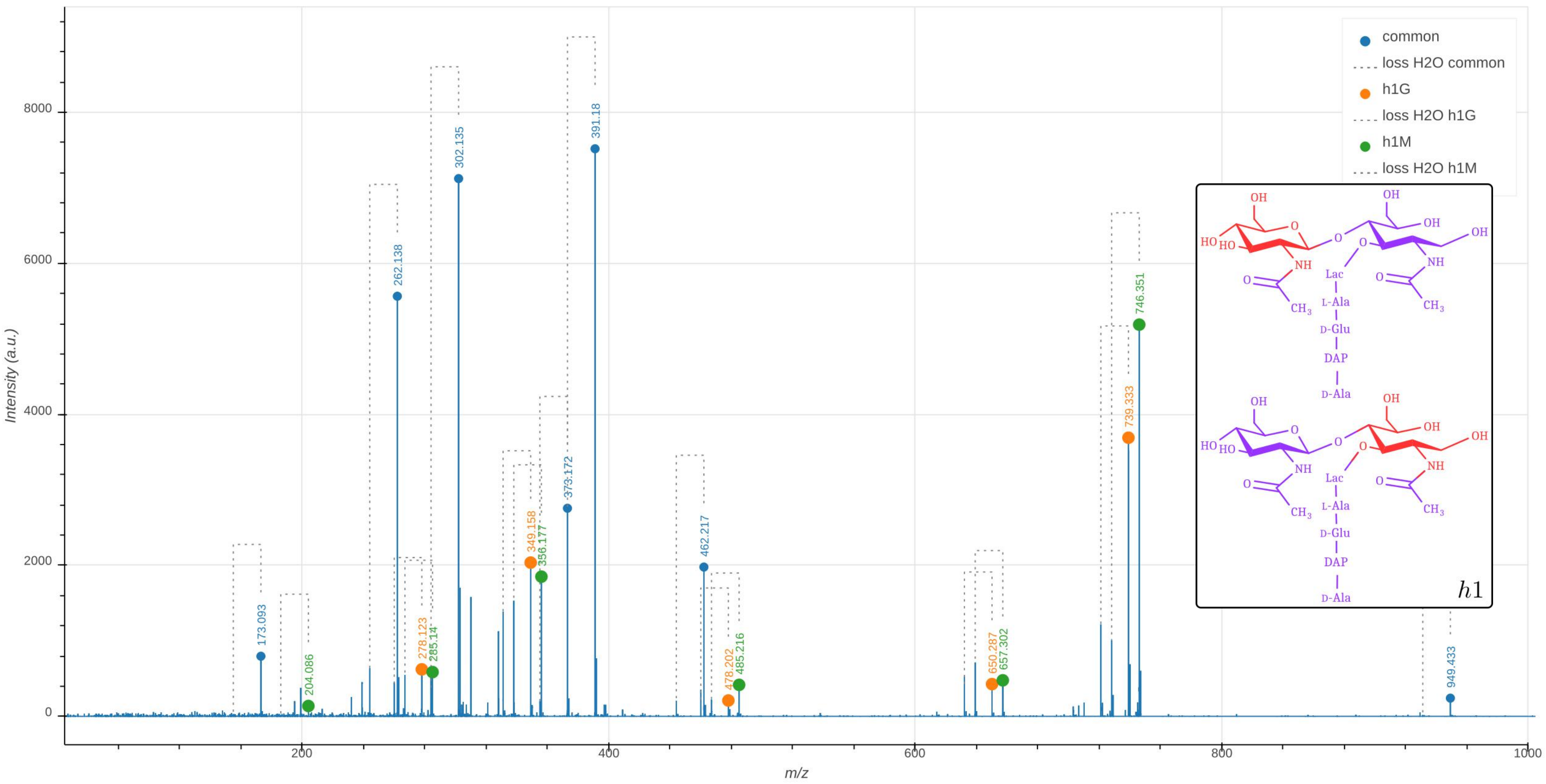

common
loss H2O common
h1G
loss H2O h1G
h1M
loss H2O h1M
173.093
204.086
262.138
278.123
285.14
302.135
349.158
356.177
373.172
391.18
462.217
478.202
485.216
650.287
657.302
739.333
746.351
949.433
Intensity (a.u.)
m/z
h1

**Supplementary Data 2.3: Tandem mass spectrometry analysis of the h2-type hybrid of the disaccharide-tetrapeptide.** The observed $m/z_{\text{obs}}$ and calculated $m/z_{\text{cal}}$ values of the parental ion, $[\text{M+H}]^+$, as determined in the absence of fragmentation (MS1), are indicated. The $m/z_{\text{calc}}$ value was used to select the ion for fragmentation. The difference between the $m/z_{\text{obs}}$ and $m/z_{\text{calc}}$ values is indicated in ppm. The GlcNAc residue is indicated in two moieties, glucosamine (GlcN) and the acetyl group (Ac), since these moieties are potentially differentially labeled. For the same reason, the reduced (Red) MurNAc residue is indicated in three moieties, reduced glucosamine ($\text{GlcN}^{\text{Red}}$), the acetyl group (Ac), and the D-lactoyl group. Labeled and unlabeled moieties are represented in red and purple, respectively. The peaks corresponding to the list of fragments appearing in the table are highlighted by orange dots in the mass spectrum. In the mass spectrum, peaks differing by the loss of $H_2 0$ are connected by a dashed line. The tables only contain the $m/z$ values for the fragments containing $H_2 0$.
An interactive report of the $\text{MS}^2$ analysis is available in Supplementary File F2.3 .

| Precursor ion (MS1) | $m/z_{obs}$ | $m/z_{calc}$ | ppm |
|---|---|---|---|
| GlcN(-Ac)-GlcN$^{\text{Red}}$(-Ac)-Lac-Ala-Glu-DAP-Ala | 961.451 | 961.454 | -2.7 |

| Product ion | $m/z_{obs}$ | $m/z_{calc}$ | ppm | Intensity (a.u.) | Isotopologue |
|---|---|---|---|---|---|
| GlcN(-Ac)-GlcN$^{\text{Red}}$(-Ac)-Lac-Ala-Glu-DAP-Ala | 961.456 | 961.454 | -2.4 | 250 | h2 |
| GlcN$^{\text{Red}}$(-Ac)-Lac-Ala-Glu-DAP-Ala | 758.374 | 758.375 | 1.0 | 15348 | h2 |
| GlcN$^{\text{Red}}$(-Ac)-Lac-Ala-Glu-DAP | 669.327 | 669.327 | -0.8 | 1466 | h2 |
| GlcN(-Ac)-GlcN$^{\text{Red}}$(-Ac)-Lac-Ala | 556.244 | 556.248 | 7.1 | 191 | h2 |
| Lac-Ala-Glu-DAP-Ala | 553.280 | 553.280 | -0.4 | 302 | h2 |
| GlcN$^{\text{Red}}$(-Ac)-Lac-Ala-Glu | 488.224 | 488.225 | 1.8 | 1041 | h2 |
| Ala-Glu-DAP-Ala | 481.259 | 481.258 | -1.0 | 3833 | h2 |
| Lac-Ala-Glu-DAP | 464.235 | 464.232 | -6.3 | 133 | h2 |
| Glu-DAP-Ala | 406.212 | 406.214 | 6.4 | 13380 | h2 |
| Ala-Glu-DAP | 392.208 | 392.211 | 6.8 | 4340 | h2 |
| GlcN$^{\text{Red}}$(-Ac)-Lac-Ala | 353.167 | 353.168 | 3.4 | 6673 | h2 |
| Glu-DAP | 317.165 | 317.167 | 3.8 | 12830 | h2 |
| GlcN$^{\text{Red}}$(-Ac)-Lac | 278.122 | 278.124 | 7.6 | 1991 | h2 |
| DAP-Ala | 271.156 | 271.158 | 7.2 | 10798 | h2 |
| GlcN$^{\text{Red}}$(-Ac) | 206.103 | 206.103 | -2.8 | 163 | h2 |
| GlcN(-Ac) | 204.085 | 204.087 | 12.7 | 229 | h2 |
| DAP | 182.108 | 182.110 | 9.5 | 1297 | h2 |

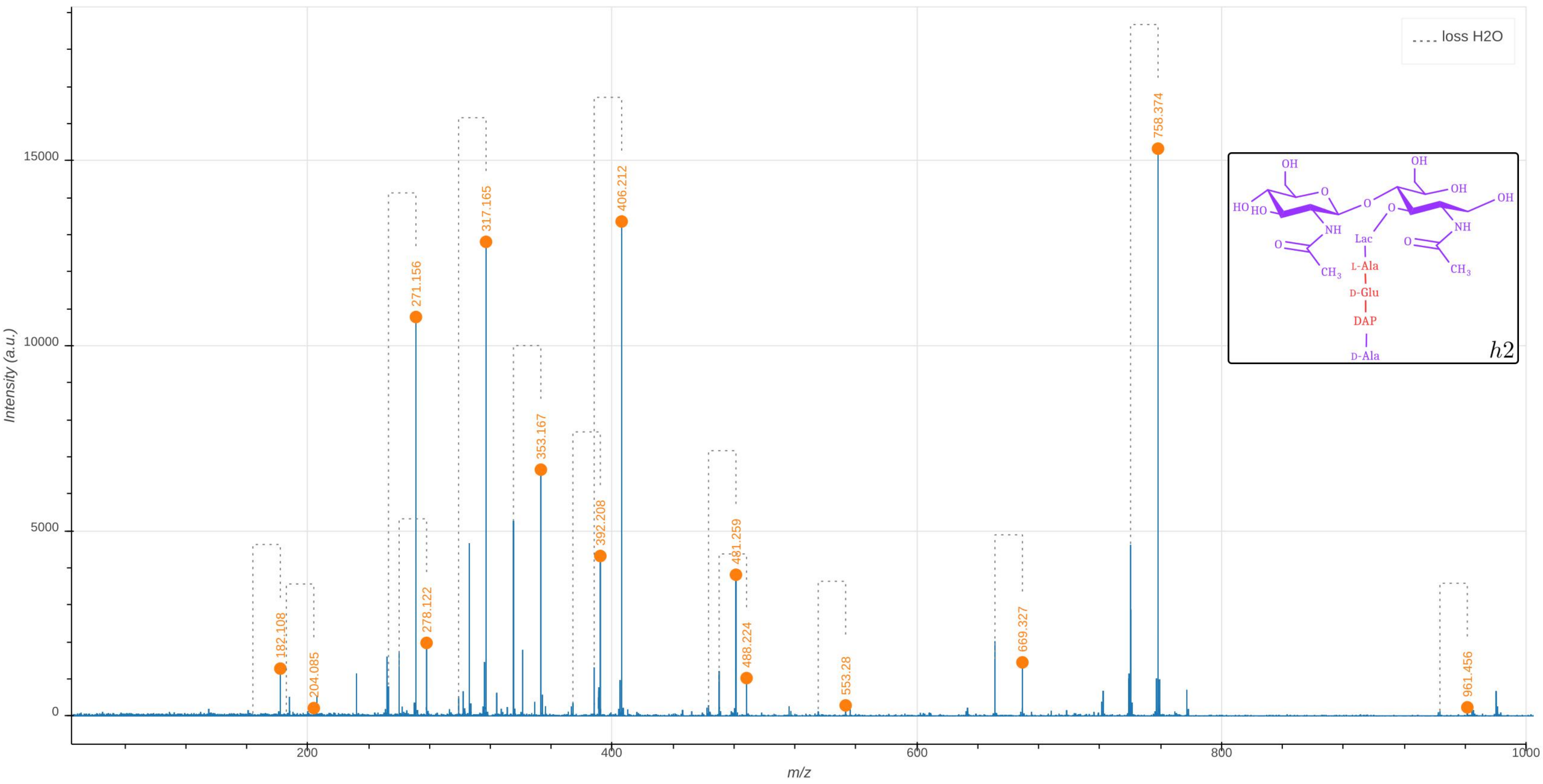

loss H2O
Intensity (a.u.)
m/z
182.108
204.085
271.156
278.122
317.165
353.167
392.208
406.212
481.259
488.224
553.28
669.327
758.374
961.456
OH
HO
HO
O
NH
O
CH3
Lac
L-Ala
D-Glu
DAP
D-Ala
OH
OH
NH
h2

**Supplementary Data 2.4: Tandem mass spectrometry analysis of the h3-type hybrid of the disaccharide-tetrapeptide.** The observed $m/z_{\mathrm{obs}}$ and calculated $m/z_{\mathrm{cal}}$ values of the parental ion, $[\mathrm{M+H}]^+$, as determined in the absence of fragmentation (MS1), are indicated. The $m/z_{\mathrm{calc}}$ value was used to select the ion for fragmentation. The difference between the $m/z_{\mathrm{obs}}$ and $m/z_{\mathrm{calc}}$ values is indicated in ppm. The GlcNAc residue is indicated in two moieties, glucosamine (GlcN) and the acetyl group (Ac), since these moieties are potentially differentially labeled. For the same reason, the reduced (Red) MurNAc residue is indicated in three moieties, reduced glucosamine ($\mathrm{GlcN^{Red}}$), the acetyl group (Ac), and the D-lactoyl group. Labeled and unlabeled moieties are represented in red and purple, respectively. The peaks corresponding to the list of fragments appearing in the table are highlighted by orange dots in the mass spectrum. In the mass spectrum, peaks differing by the loss of $H_20$ are connected by a dashed line. The tables only contain the $m/z$ values for the fragments containing $H_20$. The presence of the h3 isotopomer is expected since cells of *E. coli* contain the main PG precursor, UDP-MurNAc-pentapeptide in considerable amounts (about 2% compared to the total amount of dissaccharide peptides in the cell wall). An interactive report of the $\mathrm{MS}^2$ analysis is available in Supplementary File F2.4 .

| Precursor ion (MS1) | $m/z_{obs}$ | $m/z_{calc}$ | ppm |
|---|---|---|---|
| GlcN(-Ac)-$\mathrm{GlcN^{Red}}$(-Ac)-Lac-Ala-Glu-DAP-Ala | 977.490 | 977.49 | -5.0 |

| Product ion | $m/z_{obs}$ | $m/z_{calc}$ | ppm | Intensity (a.u.) | Isotopologue |
|---|---|---|---|---|---|
| $\mathrm{GlcN^{Red}}$(-Ac)-Lac-Ala-Glu-DAP-Ala | 774.413 | 774.416 | 3.0 | 3734 | h3 |
| $\mathrm{GlcN^{Red}}$(-Ac)-Lac-Ala-Glu-DAP | 681.368 | 681.361 | -9.8 | 306 | h3 |
| $\mathrm{GlcN^{Red}}$(-Ac)-Lac-Ala-Glu | 500.260 | 500.259 | -2.0 | 259 | h3 |
| Ala-Glu-DAP-Ala | 485.265 | 485.266 | 1.6 | 834 | h3 |
| Glu-DAP-Ala | 410.220 | 410.221 | 3.9 | 2916 | h3 |
| Ala-Glu-DAP | 392.208 | 392.211 | 6.4 | 1481 | h3 |
| $\mathrm{GlcN^{Red}}$(-Ac)-Lac-Ala | 365.199 | 365.202 | 7.3 | 1567 | h3 |
| Glu-DAP | 317.165 | 317.167 | 3.4 | 3046 | h3 |
| $\mathrm{GlcN^{Red}}$(-Ac)-Lac | 290.156 | 290.158 | 7.5 | 508 | h3 |
| DAP-Ala | 275.163 | 275.165 | 6.7 | 2174 | h3 |
| DAP | 182.110 | 182.110 | 0.1 | 354 | h3 |

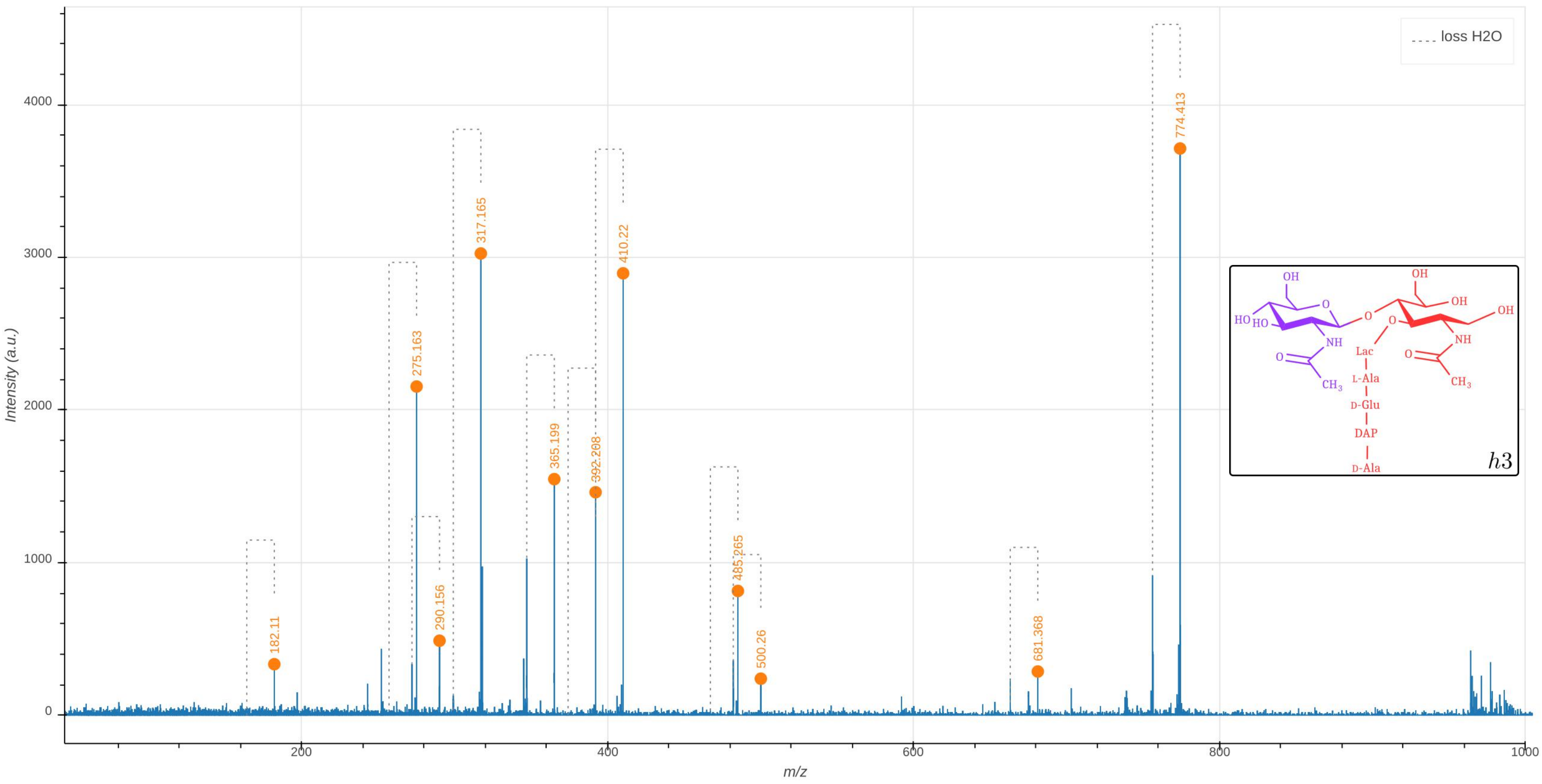

loss H2O
Intensity (a.u.)
m/z
182.11
275.163
290.156
317.165
365.199
392.208
410.22
485.265
500.26
681.368
774.413
h3
Lac
L-Ala
D-Glu
DAP
D-Ala

### Additional Monomer hybrids

**Supplementary Data 2.5: Tandem mass spectrometry analysis of the hAla-type hybrid of the disaccharide-tetrapeptide.** The observed $m/z_{\rm obs}$ and calculated $m/z_{\rm cal}$ values of the parental ion, $[M+H]^+$, as determined in the absence of fragmentation (MS1), are indicated. The $m/z_{\rm calc}$ value was used to select the ion for fragmentation. The difference between the $m/z_{\rm obs}$ and $m/z_{\rm calc}$ values is indicated in ppm. The GlcNAc residue is indicated in two moieties, glucosamine (GlcN) and the acetyl group (Ac), since these moieties are potentially differentially labeled. For the same reason, the reduced (Red) MurNAc residue is indicated in three moieties, reduced glucosamine ($\rm GlcN^{Red}$), the acetyl group (Ac), and the D-lactoyl group. Labeled and unlabeled moieties are represented in red and purple, respectively. The peaks corresponding to the list of fragments appearing in the table are highlighted by orange dots in the mass spectrum. In the mass spectrum, peaks differing by the loss of $H_20$ are connected by a dashed line. The tables only contain the $m/z$ values for the fragments containing $H_20$.
An interactive report of the MS2 analysis is available in Supplementary File F2.5.

| Parent (MS1) | $m/z_{obs}$ | $m/z_{calc}$ | ppm |
|---|---|---|---|
| GlcN(-Ac)-GlcN$^{Red}$(-Ac)-Lac-Ala-Glu-DAP-Ala | 946.420 | 946.423 | -2.9 |

| Fragment | $m/z_{obs}$ | $m/z_{calc}$ | ppm | Intensity (a.u.) | Isotopologue |
|---|---|---|---|---|---|
| GlcN(-Ac)-GlcN$^{Red}$(-Ac)-Lac-Ala-Glu-DAP-Ala | 946.414 | 946.423 | 8.8 | 227 | hAla4 |
| GlcN$^{Red}$(-Ac)-Lac-Ala-Glu-DAP-Ala | 743.340 | 743.343 | 4.3 | 1657 | hAla4 |
| Ala-Glu-DAP-Ala | 466.232 | 466.227 | -9.5 | 194 | hAla4 |
| Glu-DAP-Ala | 395.191 | 395.190 | -3.2 | 616 | hAla4 |
| Ala-Glu-DAP | 373.172 | 373.172 | 0.3 | 84 | hAla4 |
| GlcN$^{Red}$(-Ac)-Lac-Ala | 349.162 | 349.161 | -2.2 | 131 | hAla4 |
| Glu-DAP | 302.133 | 302.135 | 7.2 | 279 | hAla4 |
| GlcN$^{Red}$(-Ac)-Lac | 278.125 | 278.124 | -2.5 | 60 | hAla4 |
| DAP-Ala | 266.146 | 266.147 | 4.3 | 73 | hAla4 |
| DAP | 173.091 | 173.093 | 10.9 | 150 | hAla4 |

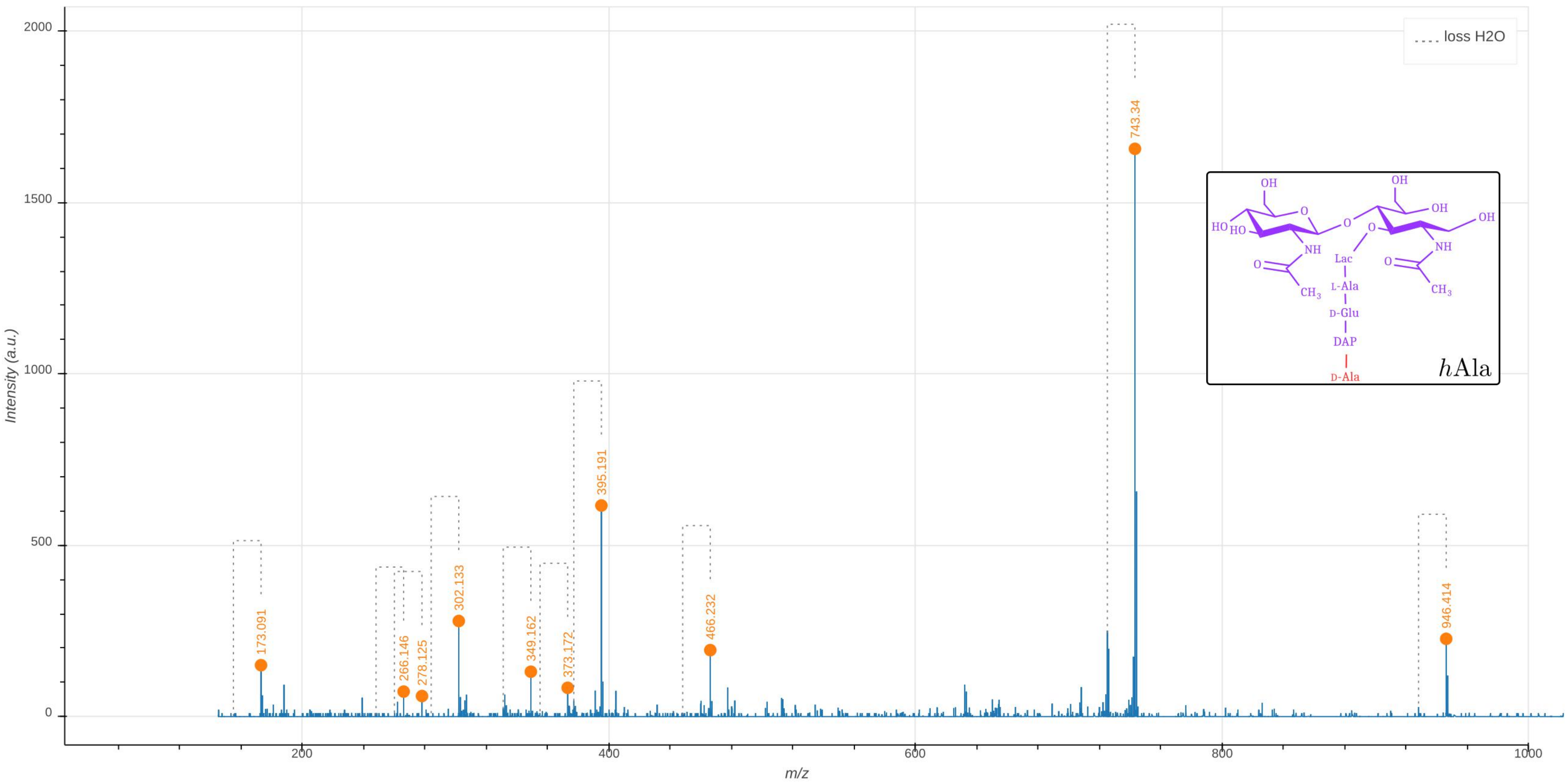

loss H2O
173.091
266.146
278.125
302.133
349.162
373.172
395.191
466.232
743.34
946.414
hAla
OH
HO
NH
O
CH3
Lac
L-Ala
D-Glu
DAP
D-Ala
Intensity (a.u.)
m/z
0
500
1000
1500
2000
200
400
600
800
1000

**Supplementary Data 2.6: Tandem mass spectrometry analysis of the h1h2-type hybrid of the disaccharide-tetrapeptide.** The molecule is fully unlabeled except for one of the two glucosamine moieties present in GlcNAc or MurNAc leading to the presence of two isotopomers.

The observed $m/z_{\text{obs}}$ and calculated $m/z_{\text{calc}}$ values of the parental ion, $[M+H]^+$, as determined in the absence of fragmentation ($MS^1$), are indicated. The $m/z_{\text{cal}}$ value was used to select the ion for fragmentation. The difference between the $m/z_{\text{obs}}$ and $m/z_{\text{cal}}$ values is indicated in ppm. The GlcNAc residue is indicated in two moieties, glucosamine (GlcN) and the acetyl group (Ac), since these moieties are potentially differentially labeled. For the same reason, the reduced (Red) MurNAc residue is indicated in three moieties, reduced glucosamine ($GlcN^{Red}$), the acetyl group (Ac), and the D-lactoyl group. Labeled and unlabeled moieties are represented in red and purple, respectively. The fragmentation led to two series of fragments specific of the isotopomers containing the unlabeled glucosamine moiety in the GlcNAc (h1Gh2) or $MurNAc^{Red}$ (h1Mh2) residues, respectively (discriminatory fragments). The fragments specific of h1Gh2 and of h1Mh2 are highlighted by orange and green dots in the mass spectrum. The other fragments are common to the fragmentation patterns of h1Gh2 and of h1Mh2. The corresponding peaks are highlighted by blue dots in the mass spectrum. The mass spectrum indicates the presence of both isotopomers as expected from the recycling and synthesis pathways since UDP-MurNAc exclusively derives from UDP-GlcNAc. In the mass spectrum, peaks differing by the loss of $H_20$ are connected by a dashed line. The tables only contain the $m/z$ values for the fragments containing $H_20$. An interactive report of the MS2 analysis is available in Supplementary File F2.6.

| Precursor ion (MS1) | $m/z_{obs}$ | $m/z_{calc}$ | ppm |
|---|---|---|---|
| GlcN(-Ac)-$GlcN^{Red}$(-Ac)-Lac-Ala-Glu-DAP-Ala | 968.471 | 968.471 | 0.2 |
| GlcN(-Ac)-$GlcN^{Red}$(-Ac)-Lac-Ala-Glu-DAP-Ala | 968.471 | 968.471 | 0.2 |

| Discriminatory product ions | $m/z_{obs}$ | $m/z_{calc}$ | ppm | Intensity (a.u.) | Isotopologue |
|---|---|---|---|---|---|
| $GlcN^{Red}$(-Ac)-Lac-Ala-Glu-DAP-Ala | 765.390 | 765.392 | 2.2 | 469 | h1Mh2 |
| $GlcN^{Red}$(-Ac)-Lac-Ala-Glu-DAP-Ala | 758.370 | 758.375 | 6.2 | 309 | h1Gh2 |

| Common product ions | $m/z_{obs}$ | $m/z_{calc}$ | ppm | Intensity (a.u.) |
|---|---|---|---|---|
| Glu-DAP-Ala | 406.212 | 406.214 | 5.8 | 151 |
| Glu-DAP | 317.163 | 317.167 | 12.6 | 121 |
| DAP | 182.108 | 182.110 | 11.6 | 60 |

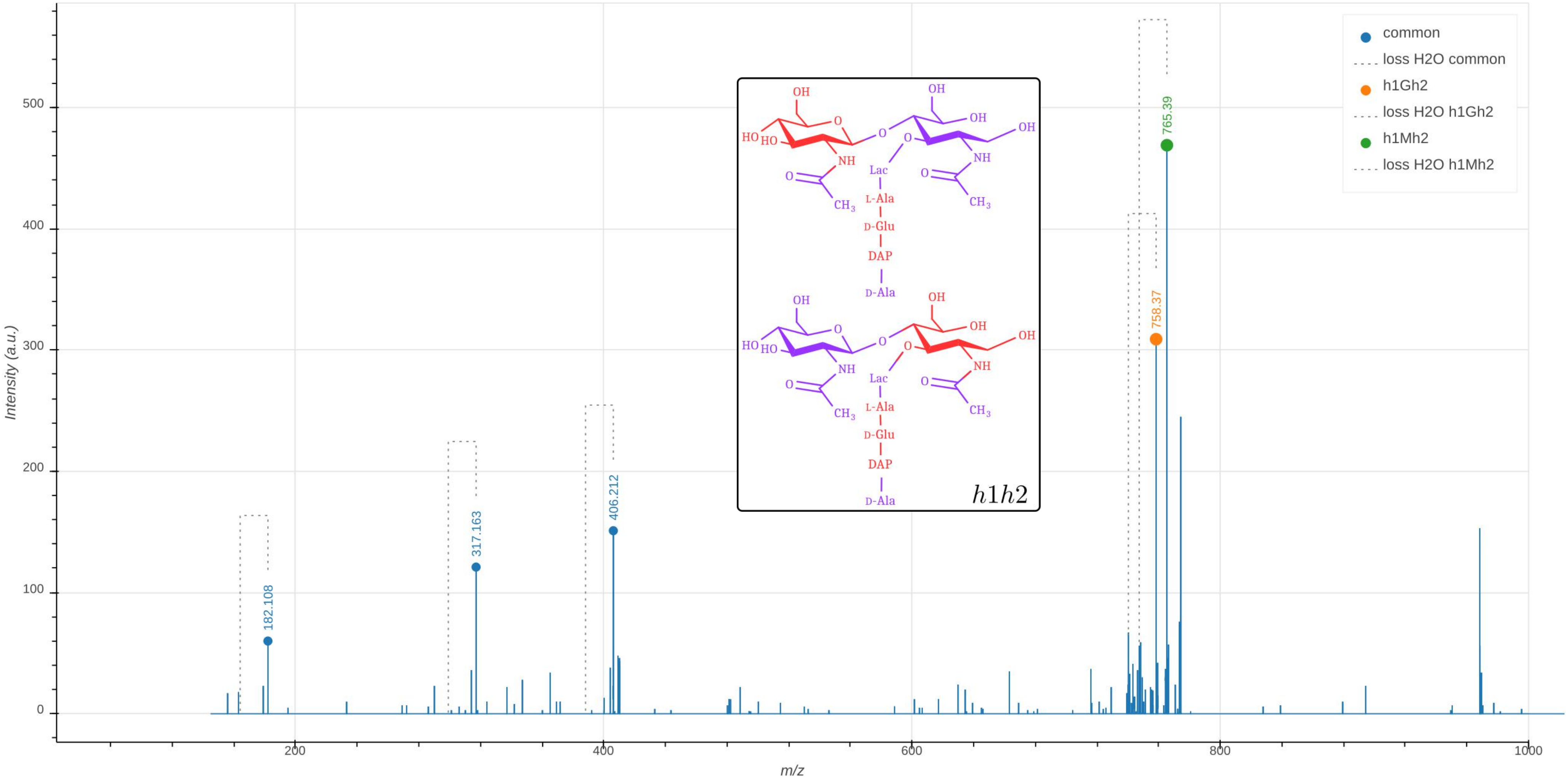

common
loss H2O common
h1Gh2
loss H2O h1Gh2
h1Mh2
loss H2O h1Mh2
182.108
317.163
406.212
758.37
765.39
Intensity (a.u.)
m/z
0
100
200
300
400
500
200
400
600
800
1000
OH
O
HO
HO
NH
O
CH3
OH
OH
OH
NH
CH3
Lac
L-Ala
D-Glu
DAP
D-Ala
h1h2

**Supplementary Data 2.7: Tandem mass spectrometry analysis of the h3Ala-type hybrid of the disaccharide-tetrapeptide.** The observed $m/z_{\text{obs}}$ and calculated $m/z_{\text{cal}}$ values of the parental ion, $[\text{M+H}]^+$, as determined in the absence of fragmentation (MS1), are indicated. The $m/z_{\text{calc}}$ value was used to select the ion for fragmentation. The difference between the $m/z_{\text{obs}}$ and $m/z_{\text{calc}}$ values is indicated in ppm. The GlcNAc residue is indicated in two moieties, glucosamine (GlcN) and the acetyl group (Ac), since these moieties are potentially differentially labeled. For the same reason, the reduced (Red) MurNAc residue is indicated in three moieties, reduced glucosamine ($\text{GlcN}^{\text{Red}}$), the acetyl group (Ac), and the D-lactoyl group. Labeled and unlabeled moieties are represented in red and purple, respectively. The peaks corresponding to the list of fragments appearing in the table are highlighted by orange dots in the mass spectrum. In the mass spectrum, peaks differing by the loss of $H_20$ are connected by a dashed line. The tables only contain the $m/z$ values for the fragments containing $H_20$.
An interactive report of the MS2 analysis is available in Supplementary File F2.7.

| Precursor Ion (MS1) | $m/z_{obs}$ | $m/z_{calc}$ | ppm |
|---|---|---|---|
| GlcN(-Ac)-GlcN$^{\text{Red}}$(-Ac)-Lac-Ala-Glu-DAP-Ala | 973.484 | 973.488 | -3.9 |

| Product ion | $m/z_{obs}$ | $m/z_{calc}$ | ppm | Intensity (a.u.) |
|---|---|---|---|---|
| GlcN(-Ac)-GlcN$^{\text{Red}}$(-Ac)-Lac-Ala-Glu-DAP-Ala | 973.492 | 973.488 | -3.7 | 161 |
| GlcN$^{\text{Red}}$(-Ac)-Lac-Ala-Glu-DAP-Ala | 770.407 | 770.409 | 2.1 | 2018 |
| GlcN$^{\text{Red}}$(-Ac)-Lac-Ala-Glu-DAP | 681.362 | 681.361 | -1.4 | 91 |
| GlcN$^{\text{Red}}$(-Ac)-Lac-Ala-Glu | 500.260 | 500.259 | -3.4 | 65 |
| Ala-Glu-DAP-Ala | 481.254 | 481.258 | 9.6 | 113 |
| Glu-DAP-Ala | 406.215 | 406.214 | -2.1 | 544 |
| Ala-Glu-DAP | 392.207 | 392.211 | 10.6 | 138 |
| GlcN$^{\text{Red}}$(-Ac)-Lac-Ala | 365.201 | 365.202 | 2.0 | 332 |
| Glu-DAP | 317.167 | 317.167 | -1.6 | 259 |
| GlcN$^{\text{Red}}$(-Ac)-Lac | 290.158 | 290.158 | -0.3 | 62 |
| DAP-Ala | 271.155 | 271.158 | 11.1 | 111 |
| DAP | 182.108 | 182.110 | 12.8 | 20 |

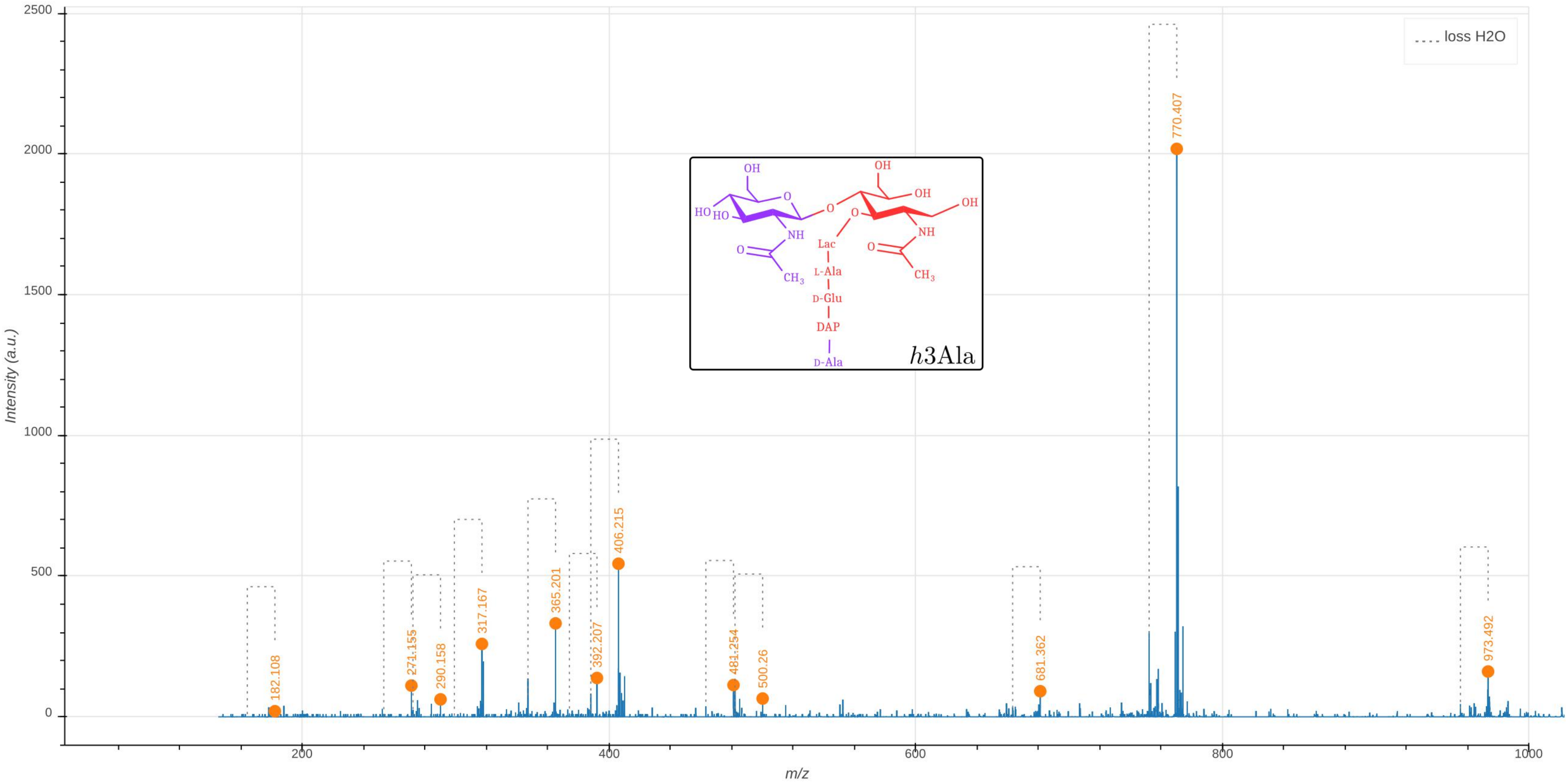

loss H2O
2500
2000
1500
1000
500
0
Intensity (a.u.)
200
400
600
800
1000
m/z
182.108
271.155
290.158
317.167
365.201
392.207
406.215
481.254
500.26
681.362
770.407
973.492
OH
O
HO
HO
NH
O
CH3
O
O
OH
OH
OH
NH
O
CH3
Lac
L-Ala
D-Glu
DAP
D-Ala
h3Ala

**A** - GM-Tripeptide

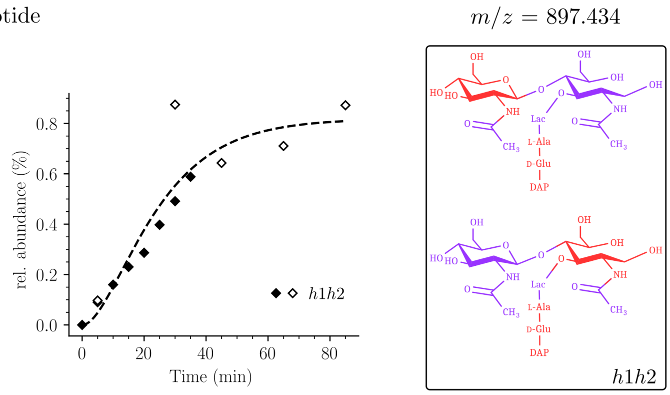


**B** - GM-Tetrapeptide

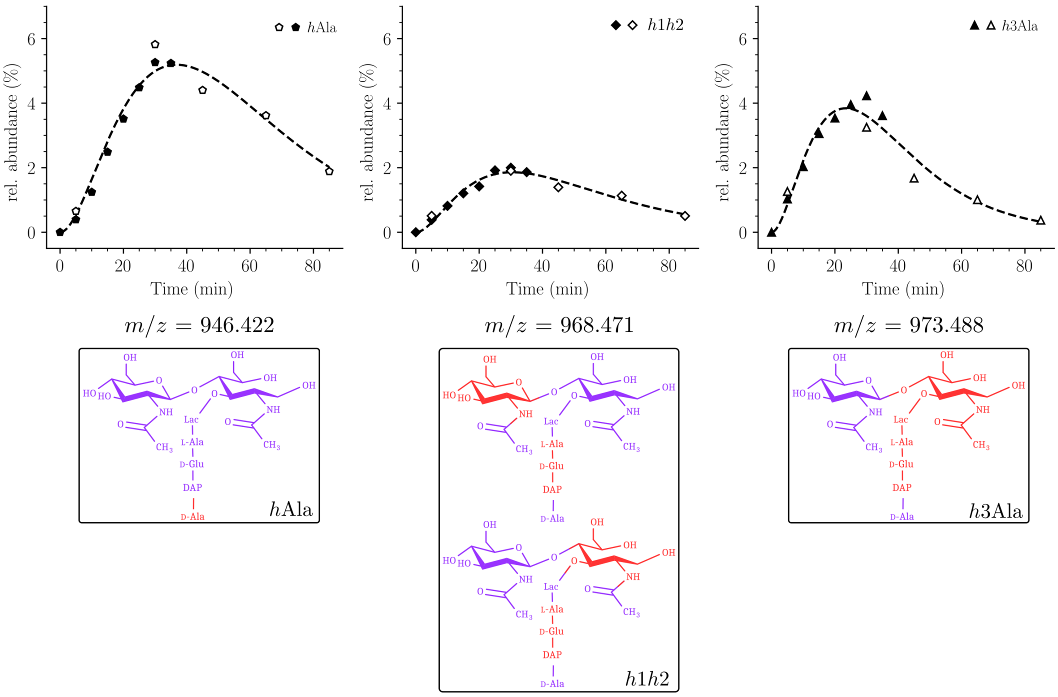


**Supplementary Data 2.8: Timecourses and structures of additional monomer hybrids.** (**A**) Structure and timecourse of the *h*1*h*2 hybrid of the disaccharide tripeptide monomer combining a recycled tripeptide stem (*h*2) with a recycled glucosamine moeity (*h*1). (**B**) Additional hybrids of the disaccharide tetrapeptide comprise *h*Ala (labeled C-terminal D-Ala$^4$), *h*1*h*2 (see (**A**)) and *h*3Ala (neo-synthesiszed GlcNAc and C-terminal D-Ala$^4$) Since disaccharide tripeptides are issued from disaccharide tetrapeptides by removal of the C-terminal D-Ala$^4$, *h*Ala and *h*3Ala are not detected for the tripeptide as these are converted into the uniformly unlabeled and *h*3 hybrid of the tripeptide, respectively. For structural characterization of the *h*Ala, *h*1*h*2, and *h*3Ala hybrids see Supplementary Data above.

### Dimers

#### Tri-Tri

**Supplementary Data 3.1: Tandem mass spectrometry analysis of the uniformly labeled Tri(3→3)Tri dimer.** The observed $m/z_{\mathrm{obs}}$ and calculated $m/z_{\mathrm{cal}}$ values of the parental ion, $[\mathrm{M+2H}]^{2+}$, as determined in the absence of fragmentation (MS1), are indicated. The $m/z_{\mathrm{calc}}$ value was used to select the ion for fragmentation. The difference between the $m/z_{\mathrm{obs}}$ and $m/z_{\mathrm{calc}}$ values is indicated in ppm. The GlcNAc residue is indicated in two moieties, glucosamine (GlcN) and the acetyl group (Ac), since these moieties are potentially differentially labeled. For the same reason, the reduced (Red) MurNAc residue is indicated in three moieties, reduced glucosamine ($\mathrm{GlcN^{Red}}$), the acetyl group (Ac), and the D-lactoyl group. Labeled and unlabeled moieties are represented in red and purple, respectively. The peaks corresponding to the list of fragments appearing in the table are highlighted by orange dots in the mass spectrum. In the mass spectrum, peaks differing by the loss of $H_20$ are connected by a dashed line. The tables only contain the $m/z$ values for the fragments containing $H_20$. The arrows indicate the position of the 3→3 and 4→3 crosslinks. Residues in the acceptor are underlined.
An interactive report of the MS2 analysis is available in Supplementary File F3.1.

| Precursor ion (MS1) | $m/z_{obs}$ [M+2H]$^{2+}$ | $m/z_{calc}$ [M+2H]$^{2+}$ | ppm |
|---|---|---|---|
| GlcN(-Ac)-GlcN$^{\mathrm{Red}}$(-Ac)-Lac-Ala-Glu-DAP→DAP-Glu-Ala-Lac-GlcN$^{\mathrm{Red}}$(-Ac)-GlcN(-Ac) | 902.464 | 902.469 | -5.9 |

| Product ion | $m/z_{obs}$ [M+H]$^{1+}$ | $m/z_{calc}$ [M+H]$^{1+}$ | ppm | Intensity (a.u.) |
|---|---|---|---|---|
| GlcN(-Ac)-GlcN$^{\mathrm{Red}}$(-Ac)-Lac-Ala-Glu-DAP→DAP-Glu-Ala-Lac-GlcN$^{\mathrm{Red}}$(-Ac) | 1591.833 | 1591.828 | -3.4 | 52 |
| GlcN$^{\mathrm{Red}}$(-Ac)-Lac-Ala-Glu-DAP→DAP-Glu-Ala-Lac-GlcN$^{\mathrm{Red}}$(-Ac) | 1379.721 | 1379.724 | 2.3 | 709 |
| GlcN$^{\mathrm{Red}}$(-Ac)-Lac-Ala-Glu-DAP→DAP-Glu-Ala | 1090.570 | 1090.574 | 3.7 | 53 |
| GlcN$^{\mathrm{Red}}$(-Ac)-Lac-Ala-Glu-DAP→DAP-Glu | 1015.523 | 1015.530 | 7.4 | 133 |
| DAP-Glu-Ala-Lac-GlcN$^{\mathrm{Red}}$(-Ac) | 699.371 | 699.371 | 1.1 | 71 |
| GlcN$^{\mathrm{Red}}$(-Ac)-Lac-Ala-Glu-DAP | 681.356 | 681.361 | 6.9 | 286 |
| GlcN$^{\mathrm{Red}}$(-Ac)-Lac-Ala | 365.200 | 365.202 | 5.6 | 77 |
| GlcN(-Ac) | 213.109 | 213.111 | 7.8 | 860 |

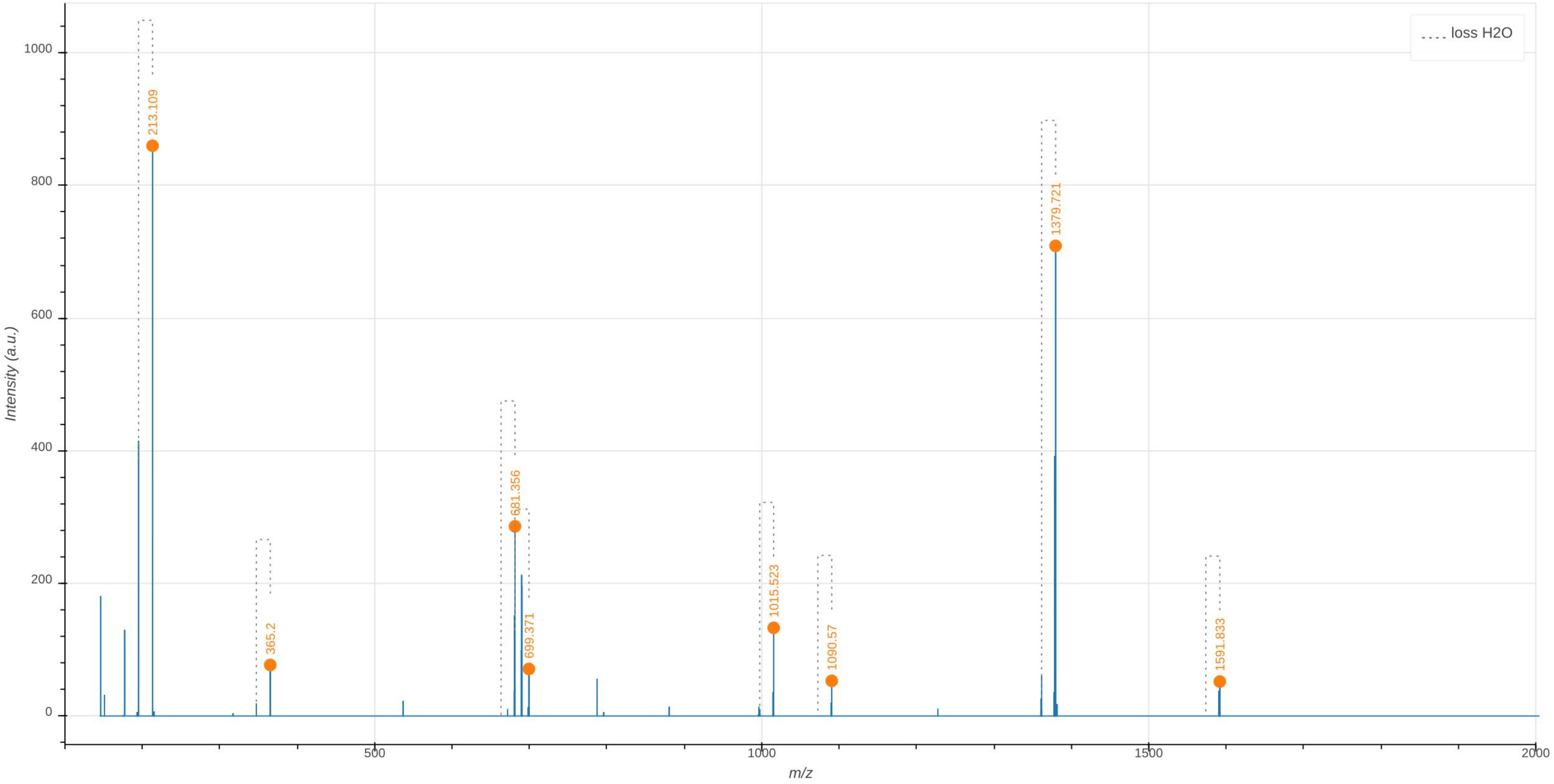

loss H2O
213.109
365.2
681.356
699.371
1015.523
1090.57
1379.721
1591.833
Intensity (a.u.)
m/z
0
200
400
600
800
1000
500
1000
1500
2000

**Supplementary Data 3.2: Tandem mass spectrometry analysis of the uniformly unlabeled Tri(3→3)Tri dimer.** The observed $m/z_{\mathrm{obs}}$ and calculated $m/z_{\mathrm{cal}}$ values of the parental ion, $[M{+}2H]^{2+}$, as determined in the absence of fragmentation (MS1), are indicated. The $m/z_{\mathrm{calc}}$ value was used to select the ion for fragmentation. The difference between the $m/z_{\mathrm{obs}}$ and $m/z_{\mathrm{calc}}$ values is indicated in ppm. The GlcNAc residue is indicated in two moieties, glucosamine (GlcN) and the acetyl group (Ac), since these moieties are potentially differentially labeled. For the same reason, the reduced (Red) MurNAc residue is indicated in three moieties, reduced glucosamine (GlcN$^{\mathrm{Red}}$), the acetyl group (Ac), and the D-lactoyl group. Labeled and unlabeled moieties are represented in red and purple, respectively. The peaks corresponding to the list of fragments appearing in the table are highlighted by orange dots in the mass spectrum. In the mass spectrum, peaks differing by the loss of $H_20$ are connected by a dashed line. The tables only contain the $m/z$ values for the fragments containing $H_20$.
The arrows indicate the position of the 3→3 and 4→3 crosslinks. Residues in the acceptor are underlined.
An interactive report of the MS2 analysis is available in Supplementary File F3.2.

| Precursor ion (MS1) | $m/z_{obs}$ $[M{+}2H]^{2+}$ | $m/z_{calc}$ $[M{+}2H]^{2+}$ | ppm |
|---|---|---|---|
| GlcN(-Ac)-GlcN$^{Red}$(-Ac)-Lac-Ala-Glu-DAP→DAP-Glu-Ala-Lac-GlcN$^{Red}$(-Ac)-GlcN(-Ac) | 862.373 | 862.373 | 0.0 |

| Product ion | $m/z_{obs}$ $[M{+}H]^{1+}$ | $m/z_{calc}$ $[M{+}H]^{1+}$ | ppm | Intensity (a.u.) |
|---|---|---|---|---|
| GlcN(-Ac)-GlcN$^{Red}$(-Ac)-Lac-Ala-Glu-DAP→DAP-Glu-Ala-Lac-GlcN$^{Red}$(-Ac) | 1520.658 | 1520.659 | 0.9 | 528 |
| GlcN$^{Red}$(-Ac)-Lac-Ala-Glu-DAP→DAP-Glu-Ala-Lac-GlcN$^{Red}$(-Ac) | 1317.581 | 1317.580 | -1.3 | 14608 |
| GlcN(-Ac)-GlcN$^{Red}$(-Ac)-Lac-Ala-Glu-DAP→DAP-Glu | 1172.522 | 1172.506 | -13.7 | 195 |
| GlcN$^{Red}$(-Ac)-Lac-Ala-Glu-DAP→DAP-Glu-Ala-Lac | 1112.491 | 1112.485 | -5.3 | 340 |
| GlcN$^{Red}$(-Ac)-Lac-Ala-Glu-DAP→DAP-Glu-Ala | 1040.466 | 1040.464 | -2.0 | 2012 |
| GlcN$^{Red}$(-Ac)-Lac-Ala-Glu-DAP→DAP-Glu | 969.428 | 969.426 | -1.3 | 10207 |
| Lac-Ala-Glu-DAP→DAP-Glu-Ala-Lac | 907.403 | 907.390 | -14.5 | 267 |
| GlcN$^{Red}$(-Ac)-Lac-Ala-Glu-DAP→DAP | 840.384 | 840.384 | -0.5 | 4637 |
| Lac-Ala-Glu-DAP→DAP-Glu-Ala | 835.377 | 835.369 | -10.5 | 208 |
| Lac-Ala-Glu-DAP→DAP-Glu | 764.336 | 764.331 | -5.8 | 517 |
| Ala-Glu-DAP→DAP-Glu-Ala | 763.357 | 763.347 | -12.3 | 114 |
| Ala-Glu-DAP→DAP-Glu | 692.315 | 692.310 | -6.1 | 860 |
| DAP-Glu-Ala-Lac-GlcN$^{Red}$(-Ac) | 668.298 | 668.299 | 1.6 | 6970 |
| GlcN$^{Red}$(-Ac)-Lac-Ala-Glu-DAP | 650.291 | 650.288 | -4.1 | 6495 |
| Lac-Ala-Glu-DAP→DAP | 635.289 | 635.289 | -0.5 | 324 |
| Glu-DAP→DAP-Glu | 621.278 | 621.273 | -7.6 | 1912 |
| Ala-Glu-DAP→DAP | 563.264 | 563.268 | 5.8 | 800 |
| Glu-DAP→DAP | 492.233 | 492.231 | -4.0 | 3051 |
| GlcN$^{Red}$(-Ac)-Lac-Ala-Glu | 478.208 | 478.204 | -9.4 | 449 |
| DAP-Glu-Ala-Lac | 463.205 | 463.204 | -2.7 | 521 |
| DAP-Glu-Ala | 391.185 | 391.183 | -5.5 | 608 |
| Ala-Glu-DAP | 373.173 | 373.172 | -0.8 | 1284 |
| DAP→DAP | 363.190 | 363.188 | -4.9 | 1374 |
| GlcN$^{Red}$(-Ac)-Lac-Ala | 349.162 | 349.161 | -1.6 | 2576 |
| DAP-Glu | 320.146 | 320.146 | -0.4 | 1973 |
| Glu-DAP | 302.136 | 302.135 | -1.6 | 3703 |
| GlcN$^{Red}$(-Ac)-Lac | 278.126 | 278.124 | -6.0 | 576 |

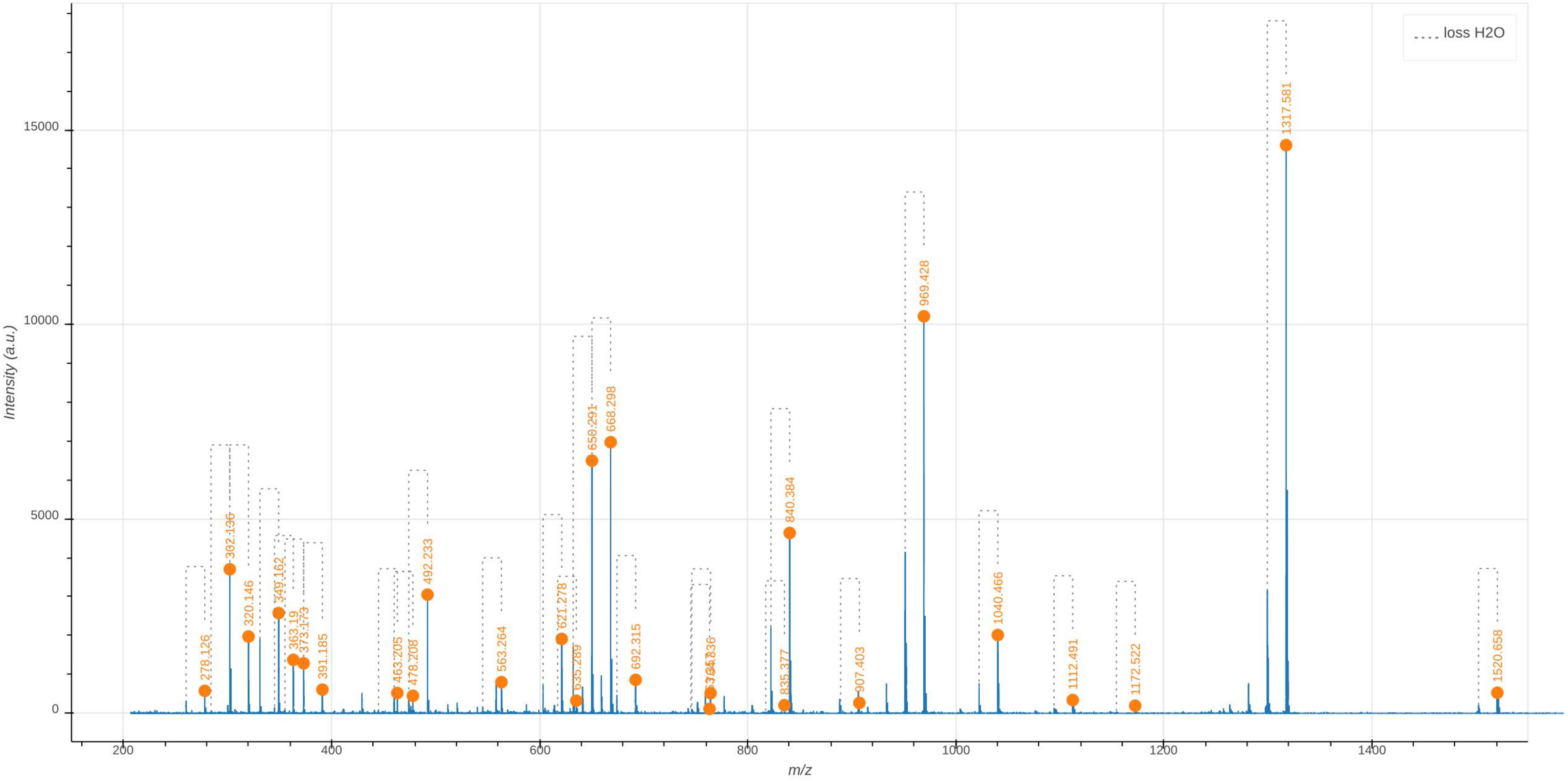
loss H2O
Intensity (a.u.)
m/z
0
5000
10000
15000
200
400
600
800
1000
1200
1400
278.126
302.136
320.146
349.162
363.19
373.173
391.185
463.205
478.208
492.233
563.264
621.278
635.289
650.291
668.298
692.315
835.377
840.384
907.403
969.428
1040.466
1112.491
1172.522
1317.581
1520.658

**Supplementary Data 3.3: Tandem mass spectrometry analysis of the Tri→Tri dimer containing a uniformly labeled disaccharide-tripeptide cross-linked to a uniformly unlabeled disaccharide-tripeptide (3→3 cross-link).** The observed $m/z_{\mathrm{obs}}$ and calculated $m/z_{\mathrm{cal}}$ values of the parental ion, $[\mathrm{M+2H}]^{2+}$, as determined in the absence of fragmentation (MS1), are indicated. The $m/z_{\mathrm{calc}}$ value was used to select the ion for fragmentation. The difference between the $m/z_{\mathrm{obs}}$ and $m/z_{\mathrm{calc}}$ values is indicated in ppm. The GlcNAc residue is indicated in two moieties, glucosamine (GlcN) and the acetyl group (Ac), since these moieties are potentially differentially labeled. For the same reason, the reduced (Red) MurNAc residue is indicated in three moieties, reduced glucosamine (GlcN$^{\mathrm{Red}}$), the acetyl group (Ac), and the D-lactoyl group. Labeled and unlabeled moieties are represented in red and purple, respectively. The fragmentation potentially leads to two series of fragments (discriminatory fragments) specific of the isotopomers containing the fully labeled disaccharide subunit in the donor (all heavy-all light) or acceptor position (all light-all heavy). These fragments are highlighted by green and orange dots in the mass spectrum, respectively. The other fragments are common to the fragmentation of the two isotopomers (highlighted by blue dots in the mass spectrum). The mass spectrum indicates the presence of the all light-all heavy isotopomers (orange dots). The peak at $m/z_{obs}$ 681.358 (green dot) can also be accounted for by the loss of $H_20$ from the peak at 699.367 indicating that all peaks may be accounted for by the presence of the all light-all heavy isotopomer.

In the mass spectrum, peaks differing by the loss of $H_20$ are connected by a dashed line. The arrows indicate the position of the 3→3 and 4→3 crosslinks. Residues in the acceptor are underlined.

An interactive report of the MS2 analysis is available in Supplementary File F3.3.

| Precursor ion (MS1) | $m/z_{obs}$ $[\mathrm{M+2H}]^{2+}$ | $m/z_{calc}$ $[\mathrm{M+2H}]^{2+}$ | ppm |
|---|---|---|---|
| GlcN(-Ac)-GlcN$^{Red}$(-Ac)-Lac-Ala-Glu-DAP→DAP-Glu-Ala-Lac-GlcN$^{Red}$(-Ac)-GlcN(-Ac) | 882.421 | 882.421 | -0.0 |
| GlcN(-Ac)-GlcN$^{Red}$(-Ac)-Lac-Ala-Glu-DAP→DAP-Glu-Ala-Lac-GlcN$^{Red}$(-Ac)-GlcN(-Ac) | 882.421 | 882.421 | -0.0 |

| Discriminatory product ions | $m/z_{obs}$ $[\mathrm{M+H}]^{1+}$ | $m/z_{calc}$ $[\mathrm{M+H}]^{1+}$ | ppm | Intensity (a.u.) | Isotopologue |
|---|---|---|---|---|---|
| DAP-Glu-Ala-Lac-GlcN$^{Red}$(-Ac) | 699.367 | 699.371 | 6.5 | 678 | all light-all heavy |
| GlcN$^{Red}$(-Ac)-Lac-Ala-Glu-DAP | 681.358 | 681.361 | 4.8 | 325 | all heavy-all light |
| DAP-Glu | 335.173 | 335.177 | 13.8 | 198 | all light-all heavy |
| Glu-DAP | 302.132 | 302.135 | 11.5 | 281 | all light-all heavy |

| Common product ions | $m/z_{obs}$ $[\mathrm{M+H}]^{1+}$ | $m/z_{calc}$ $[\mathrm{M+H}]^{1+}$ | ppm | Intensity (a.u.) |
|---|---|---|---|---|
| GlcN$^{Red}$(-Ac)-Lac-Ala-Glu-DAP→DAP-Glu-Ala-Lac-GlcN$^{Red}$(-Ac) | 1348.645 | 1348.652 | 5.6 | 912 |
| GlcN$^{Red}$(-Ac)-Lac-Ala-Glu-DAP→DAP-Glu-Ala-Lac-GlcN$^{Red}$(-Ac) | | | | |
| GlcN$^{Red}$(-Ac)-Lac-Ala-Glu-DAP→DAP-Glu | 1000.498 | 1000.499 | 0.6 | 242 |
| Glu-DAP→DAP-Glu-Ala-Lac-GlcN$^{Red}$(-Ac) | | | | |
| GlcN$^{Red}$(-Ac)-Lac-Ala-Glu-DAP→DAP-Glu | 984.450 | 984.458 | 7.6 | 428 |
| Glu-DAP→DAP-Glu-Ala-Lac-GlcN$^{Red}$(-Ac) | | | | |
| GlcN$^{Red}$(-Ac)-Lac-Ala-Glu-DAP→DAP | 871.456 | 871.456 | 0.5 | 183 |
| DAP→DAP-Glu-Ala-Lac-GlcN$^{Red}$(-Ac) | | | | |
| DAP→DAP-Glu-Ala-Lac-GlcN$^{Red}$(-Ac) | 849.398 | 849.401 | 3.8 | 136 |
| GlcN$^{Red}$(-Ac)-Lac-Ala-Glu-DAP→DAP | | | | |
| Glu-DAP→DAP-Glu | 636.300 | 636.305 | 7.0 | 163 |
| Glu-DAP→DAP-Glu | | | | |
| DAP→DAP-Glu | 507.260 | 507.262 | 3.0 | 190 |
| Glu-DAP→DAP | | | | |
| DAP→DAP | 372.198 | 372.206 | 19.4 | 125 |
| DAP→DAP | | | | |
| GlcN$^{Red}$(-Ac)-Lac-Ala | 365.199 | 365.202 | 9.2 | 182 |
| Ala-Lac-GlcN$^{Red}$(-Ac) | | | | |
| GlcN(-Ac) | 213.109 | 213.111 | 9.2 | 441 |
| GlcN(-Ac) | | | | |
| GlcN(-Ac) | 204.086 | 204.087 | 5.9 | 951 |
| GlcN(-Ac) | | | | |

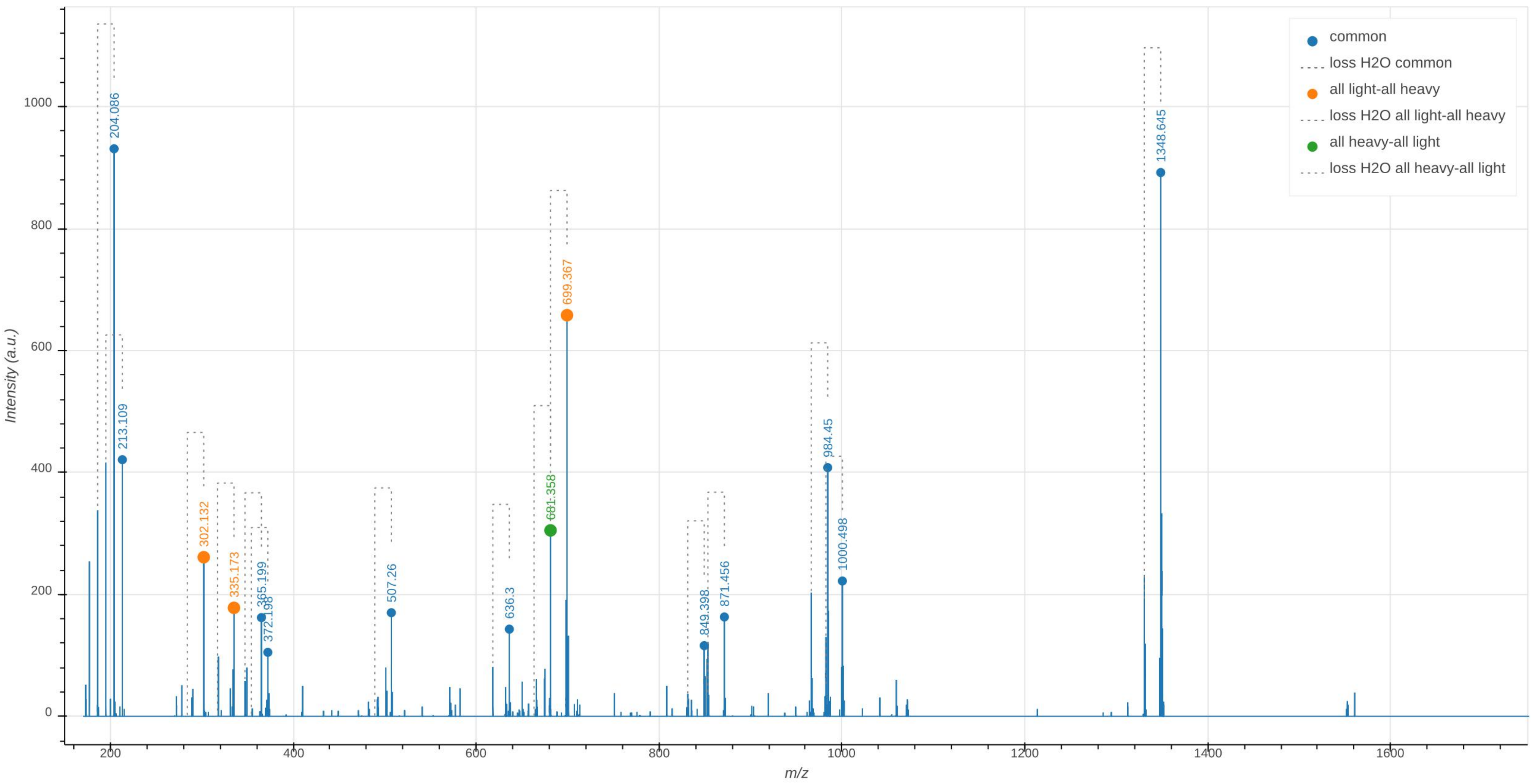

common
loss H2O common
all light-all heavy
loss H2O all light-all heavy
all heavy-all light
loss H2O all heavy-all light
204.086
213.109
302.132
335.173
365.199
372.198
507.26
636.3
681.358
699.367
849.398
871.456
984.45
1000.498
1348.645
Intensity (a.u.)
m/z

### Tetra-Tri

**Supplementary Data 4.1: Tandem mass spectrometry analysis of the uniformly labeled Tetra(4→3)Tri dimer.** The observed $m/z_{\text{obs}}$ and calculated $m/z_{\text{cal}}$ values of the parental ion, $[\text{M+2H}]^{2+}$, as determined in the absence of fragmentation (MS1), are indicated. The $m/z_{\text{calc}}$ value was used to select the ion for fragmentation. The difference between the $m/z_{\text{obs}}$ and $m/z_{\text{calc}}$ values is indicated in ppm. The GlcNAc residue is indicated in two moieties, glucosamine (GlcN) and the acetyl group (Ac), since these moieties are potentially differentially labeled. For the same reason, the reduced (Red) MurNAc residue is indicated in three moieties, reduced glucosamine ($\text{GlcN}^{\text{Red}}$), the acetyl group (Ac), and the D-lactoyl group. Labeled and unlabeled moieties are represented in red and purple, respectively. The peaks corresponding to the list of fragments appearing in the table are highlighted by orange dots in the mass spectrum. In the mass spectrum, peaks differing by the loss of $H_20$ are connected by a dashed line. The tables only contain the $m/z$ values for the fragments containing $H_20$. The arrows indicate the position of the 3→3 and 4→3 crosslinks. Residues in the acceptor are underlined.
An interactive report of the MS2 analysis is available in Supplementary File F4.1.

| Precursor ion (MS1) | $m/z_{obs}$ $[\text{M+2H}]^{2+}$ | $m/z_{calc}$ $[\text{M+2H}]^{2+}$ | ppm |
|---|---|---|---|
| GlcN(-Ac)-GlcN$^{\text{Red}}$(-Ac)-Lac-Ala-Glu-DAP-Ala→DAP-Glu-Ala-Lac-GlcN$^{\text{Red}}$(-Ac)-GlcN(-Ac) | 939.992 | 939.992 | 0.1 |

| Product ion | $m/z_{obs}$ $[\text{M+H}]^{1+}$ | $m/z_{calc}$ $[\text{M+H}]^{1+}$ | ppm | Intensity (a.u.) |
|---|---|---|---|---|
| GlcN(-Ac)-GlcN$^{\text{Red}}$(-Ac)-Lac-Ala-Glu-DAP-Ala→DAP-Glu-Ala-Lac-GlcN$^{\text{Red}}$(-Ac) | 1666.880 | 1666.872 | -4.9 | 113 |
| GlcN$^{\text{Red}}$(-Ac)-Lac-Ala-Glu-DAP-Ala→DAP-Glu-Ala-Lac-GlcN$^{\text{Red}}$(-Ac) | 1454.780 | 1454.769 | -7.6 | 462 |
| GlcN$^{\text{Red}}$(-Ac)-Lac-Ala-Glu-DAP-Ala→DAP-Glu-Ala | 1165.627 | 1165.619 | -6.9 | 103 |
| GlcN$^{\text{Red}}$(-Ac)-Lac-Ala-Glu-DAP-Ala→DAP-Glu | 1090.570 | 1090.574 | 3.6 | 240 |
| GlcN$^{\text{Red}}$(-Ac)-Lac-Ala-Glu-DAP-Ala→DAP | 955.520 | 955.518 | -2.5 | 170 |
| Ala-Glu-DAP-Ala→DAP-Glu | 801.426 | 801.424 | -2.3 | 43 |
| Ala→DAP-Glu-Ala-Lac-GlcN$^{\text{Red}}$(-Ac) | 774.416 | 774.416 | -0.7 | 342 |
| GlcN$^{\text{Red}}$(-Ac)-Lac-Ala-Glu-DAP-Ala | 756.413 | 756.405 | -11.1 | 181 |
| Glu-DAP-Ala→DAP-Glu | 726.390 | 726.380 | -13.5 | 50 |
| DAP-Glu-Ala-Lac-GlcN$^{\text{Red}}$(-Ac) | 699.369 | 699.371 | 2.9 | 166 |
| GlcN$^{\text{Red}}$(-Ac)-Lac-Ala-Glu-DAP | 681.366 | 681.361 | -6.8 | 54 |
| Ala-Glu-DAP | 392.216 | 392.211 | -12.7 | 44 |
| GlcN(-Ac) | 213.112 | 213.111 | -5.5 | 1238 |

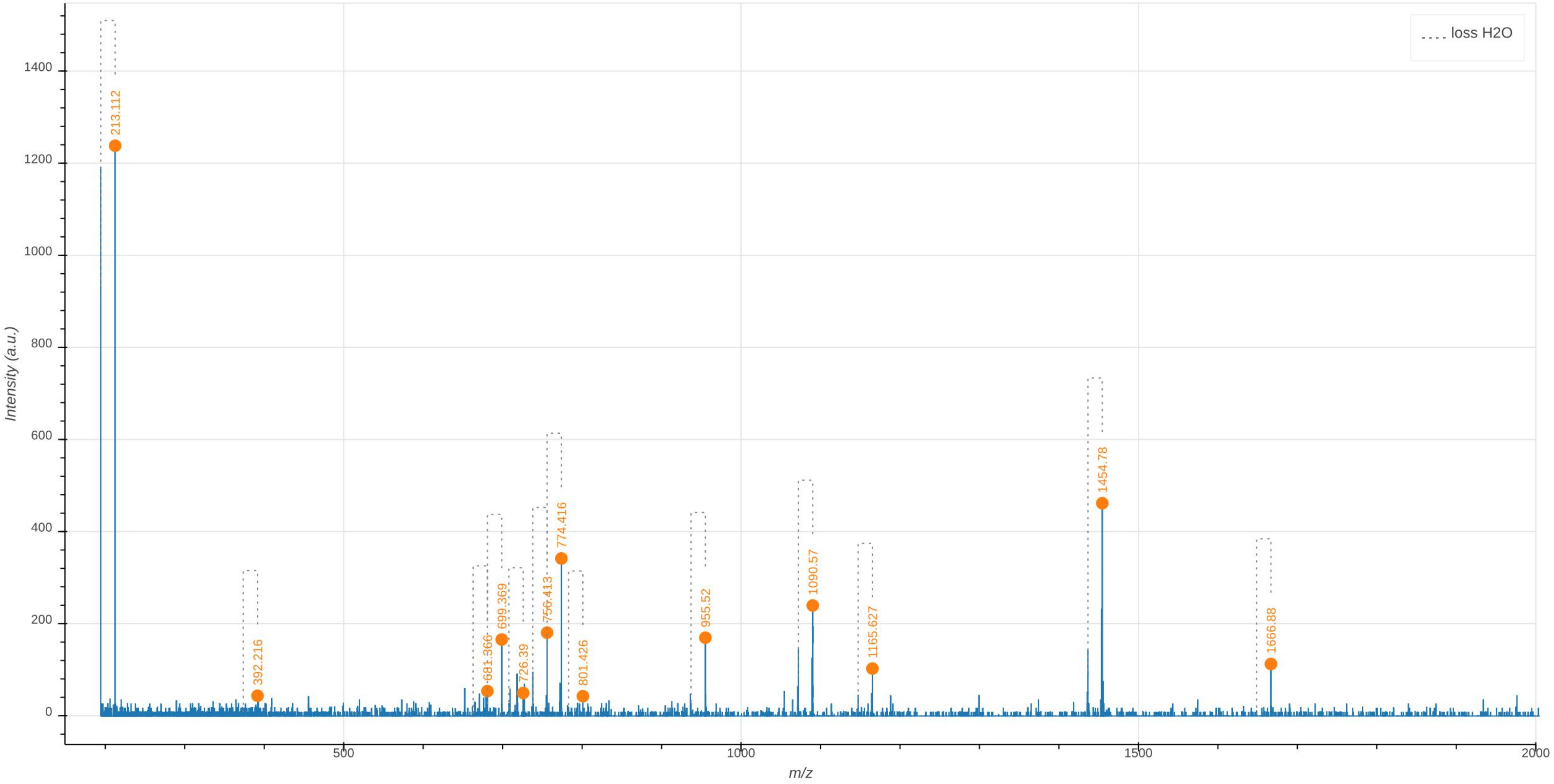

loss H2O
213.112
392.216
681.366
699.369
726.39
756.413
774.416
801.426
955.52
1090.57
1165.627
1454.78
1666.88
Intensity (a.u.)
m/z
0
200
400
600
800
1000
1200
1400
500
1000
1500
2000

**Supplementary Data 4.2: Tandem mass spectrometry analysis of the uniformly unlabeled Tri(4→3)Tri dimer.** The observed $m/z_{\text{obs}}$ and calculated $m/z_{\text{cal}}$ values of the parental ion, $[M+2H]^{2+}$, as determined in the absence of fragmentation (MS1), are indicated. The $m/z_{\text{calc}}$ value was used to select the ion for fragmentation. The difference between the $m/z_{\text{obs}}$ and $m/z_{\text{calc}}$ values is indicated in ppm. The GlcNAc residue is indicated in two moieties, glucosamine (GlcN) and the acetyl group (Ac), since these moieties are potentially differentially labeled. For the same reason, the reduced (Red) MurNAc residue is indicated in three moieties, reduced glucosamine ($GlcN^{Red}$), the acetyl group (Ac), and the D-lactoyl group. Labeled and unlabeled moieties are represented in red and purple, respectively. The peaks corresponding to the list of fragments appearing in the table are highlighted by orange dots in the mass spectrum. In the mass spectrum, peaks differing by the loss of $H_20$ are connected by a dashed line. The tables only contain the $m/z$ values for the fragments containing $H_20$.
The arrows indicate the position of the 3→3 and 4→3 crosslinks. Residues in the acceptor are underlined.
An interactive report of the MS2 analysis is available in Supplementary File F4.2.

| Precursor ion (MS1) | $m/z_{obs}$ $[M+2H]^{2+}$ | $m/z_{calc}$ $[M+2H]^{2+}$ | ppm |
|---|---|---|---|
| GlcN(-Ac)-GlcN$^{Red}$(-Ac)-Lac-Ala-Glu-DAP-Ala→DAP-Glu-Ala-Lac-GlcN$^{Red}$(-Ac)-GlcN(-Ac) | 897.889 | 897.892 | -3.3 |

| Product ion | $m/z_{obs}$ $[M+H]^{1+}$ | $m/z_{calc}$ $[M+H]^{1+}$ | ppm | Intensity (a.u.) |
|---|---|---|---|---|
| GlcN(-Ac)-GlcN$^{Red}$(-Ac)-Lac-Ala-Glu-DAP-Ala→DAP-Glu-Ala-Lac-GlcN$^{Red}$(-Ac) | 1591.693 | 1591.696 | 1.9 | 144 |
| GlcN$^{Red}$(-Ac)-Lac-Ala-Glu-DAP-Ala→DAP-Glu-Ala-Lac-GlcN$^{Red}$(-Ac) | 1388.622 | 1388.617 | -3.4 | 1802 |
| GlcN$^{Red}$(-Ac)-Lac-Ala-Glu-DAP-Ala→DAP-Glu-Ala | 1111.500 | 1111.501 | 0.4 | 395 |
| GlcN$^{Red}$(-Ac)-Lac-Ala-Glu-DAP-Ala→DAP-Glu | 1040.474 | 1040.464 | -10.3 | 1464 |
| GlcN$^{Red}$(-Ac)-Lac-Ala-Glu-DAP-Ala→DAP | 911.426 | 911.421 | -5.8 | 677 |
| Ala-Glu-DAP-Ala→DAP-Glu | 763.357 | 763.347 | -13.1 | 209 |
| Ala→DAP-Glu-Ala-Lac-GlcN$^{Red}$(-Ac) | 739.335 | 739.336 | 0.9 | 2041 |
| GlcN$^{Red}$(-Ac)-Lac-Ala-Glu-DAP-Ala | 721.325 | 721.326 | 0.5 | 1560 |
| Glu-DAP-Ala→DAP-Glu | 692.308 | 692.310 | 3.2 | 291 |
| DAP-Glu-Ala-Lac-GlcN$^{Red}$(-Ac) | 668.297 | 668.299 | 2.4 | 618 |
| GlcN$^{Red}$(-Ac)-Lac-Ala-Glu-DAP | 650.288 | 650.288 | 1.3 | 370 |
| Ala-Glu-DAP-Ala→DAP | 634.304 | 634.305 | 1.7 | 218 |
| Glu-DAP-Ala→DAP | 563.267 | 563.268 | 1.2 | 409 |
| Ala→DAP-Glu-Ala | 462.221 | 462.220 | -1.1 | 137 |
| DAP-Ala→DAP | 434.225 | 434.225 | -0.2 | 298 |
| Ala→DAP-Glu | 391.183 | 391.183 | 0.3 | 451 |
| Ala-Glu-DAP | 373.173 | 373.172 | -2.5 | 355 |
| GlcN$^{Red}$(-Ac)-Lac-Ala | 349.161 | 349.161 | -1.1 | 557 |
| DAP-Glu | 320.146 | 320.146 | -1.6 | 252 |
| Glu-DAP | 302.135 | 302.135 | -0.7 | 366 |
| GlcN$^{Red}$(-Ac)-Lac | 278.125 | 278.124 | -5.0 | 107 |

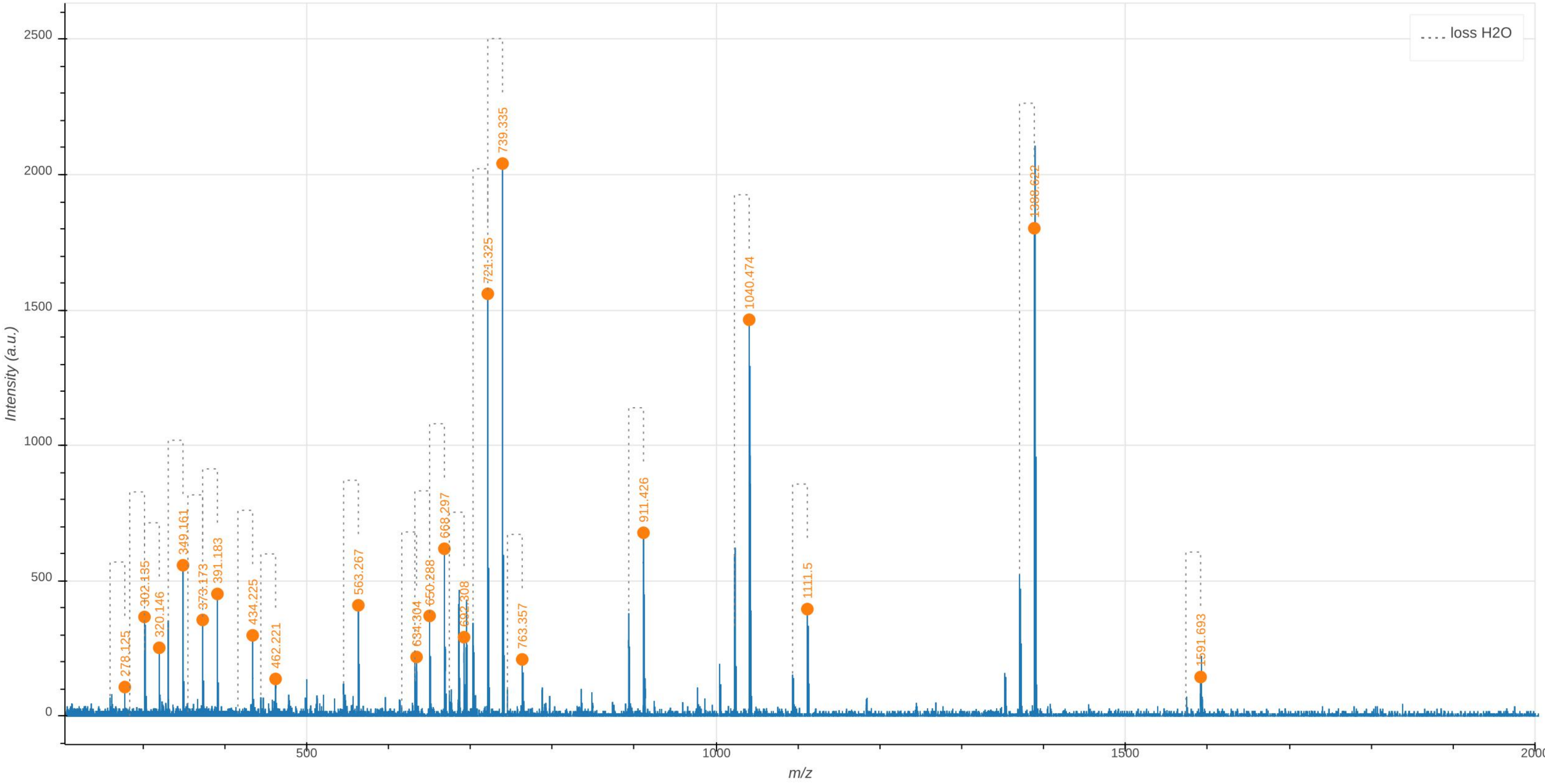

loss H2O
Intensity (a.u.)
m/z
0
500
1000
1500
2000
2500
500
1000
1500
2000
278.125
302.135
320.146
349.161
373.173
391.183
434.225
462.221
563.267
634.304
650.288
668.297
692.308
721.325
739.335
763.357
911.426
1040.474
1111.5
1388.622
1591.693

**Supplementary Data 4.3: Tandem mass spectrometry analysis of the Tetra→Tri dimer containing a uniformly labeled disaccharide-tetrapeptide cross-linked to a uniformly unlabeled disaccharide-tripeptide (4→3 cross-link).** The observed $m/z_{\text{obs}}$ and calculated $m/z_{\text{cal}}$ values of the parental ion, $[M+2H]^{2+}$, as determined in the absence of fragmentation (MS1), are indicated. The $m/z_{\text{calc}}$ value was used to select the ion for fragmentation. The difference between the $m/z_{\text{obs}}$ and $m/z_{\text{calc}}$ values is indicated in ppm. The GlcNAc residue is indicated in two moieties, glucosamine (GlcN) and the acetyl group (Ac), since these moieties are potentially differentially labeled. For the same reason, the reduced (Red) MurNAc residue is indicated in three moieties, reduced glucosamine ($\text{GlcN}^{\text{Red}}$), the acetyl group (Ac), and the D-lactoyl group. Labeled and unlabeled moieties are represented in red and purple, respectively.

In the mass spectrum, peaks differing by the loss of $H_20$ are connected by a dashed line. The arrows indicate the position of the 3→3 and 4→3 crosslinks. Residues in the acceptor are underlined.

An interactive report of the MS2 analysis is available in Supplementary File F4.3.

| Precursor ion (MS1) | $m/z_{obs}$ $[M+2H]^{2+}$ | $m/z_{calc}$ $[M+2H]^{2+}$ | ppm |
|---|---|---|---|
| GlcN(-Ac)-GlcN$^{\text{Red}}$(-Ac)-Lac-Ala-Glu-DAP-Ala→DAP-Glu-Ala-Lac-GlcN$^{\text{Red}}$(-Ac)-GlcN(-Ac) | 917.936 | 917.940 | -4.4 |

| Discriminatory product ion | $m/z_{obs}$ $[M+H]^{1+}$ | $m/z_{calc}$ $[M+H]^{1+}$ | ppm | Intensity (a.u.) |
|---|---|---|---|---|
| GlcN$^{\text{Red}}$(-Ac)-Lac-Ala-Glu-DAP-Ala→DAP-Glu-Ala-Lac-GlcN$^{\text{Red}}$(-Ac) | 1419.689 | 1419.689 | 0.3 | 345 |
| Glu-DAP-Ala→DAP-Glu-Ala-Lac-GlcN$^{\text{Red}}$(-Ac) | 1071.533 | 1071.536 | 2.7 | 71 |
| GlcN$^{\text{Red}}$(-Ac)-Lac-Ala-Glu-DAP-Ala→DAP-Glu | 1055.499 | 1055.495 | -3.4 | 63 |
| DAP-Ala→DAP-Glu-Ala-Lac-GlcN$^{\text{Red}}$(-Ac) | 942.496 | 942.493 | -3.2 | 172 |
| GlcN$^{\text{Red}}$(-Ac)-Lac-Ala-Glu-DAP-Ala→DAP | 920.433 | 920.439 | 5.9 | 44 |
| Glu-DAP-Ala→DAP-Glu-Ala | 782.389 | 782.386 | -3.8 | 26 |
| Ala→DAP-Glu-Ala-Lac-GlcN$^{\text{Red}}$(-Ac) | 770.407 | 770.409 | 1.4 | 190 |
| GlcN$^{\text{Red}}$(-Ac)-Lac-Ala-Glu-DAP-Ala | 721.325 | 721.326 | 0.7 | 148 |
| DAP-Glu-Ala-Lac-GlcN$^{\text{Red}}$(-Ac) | 699.381 | 699.371 | -14.3 | 71 |
| GlcN$^{\text{Red}}$(-Ac)-Lac-Ala-Glu-DAP | 650.294 | 650.288 | -8.5 | 23 |
| DAP-Ala→DAP-Glu | 578.299 | 578.299 | 0.2 | 27 |
| Glu-DAP-Ala→DAP | 572.284 | 572.285 | 2.7 | 51 |
| Ala→DAP-Glu-Ala-Lac | 556.295 | 556.290 | -9.4 | 37 |
| Ala-Glu-DAP-Ala | 444.207 | 444.209 | 6.0 | 42 |
| DAP-Ala→DAP | 443.242 | 443.243 | 1.9 | 81 |
| Ala→DAP-Glu | 406.212 | 406.214 | 5.0 | 159 |
| Ala-Glu-DAP | 373.173 | 373.172 | -0.5 | 63 |
| Ala-Lac-GlcN$^{\text{Red}}$(-Ac) | 365.199 | 365.202 | 8.0 | 97 |
| GlcN$^{\text{Red}}$(-Ac)-Lac-Ala | 349.162 | 349.161 | -1.4 | 40 |
| Glu-DAP | 302.134 | 302.135 | 4.8 | 51 |
| GlcN(-Ac) | 213.110 | 213.111 | 6.5 | 235 |
| GlcN(-Ac) | 204.086 | 204.087 | 7.7 | 502 |

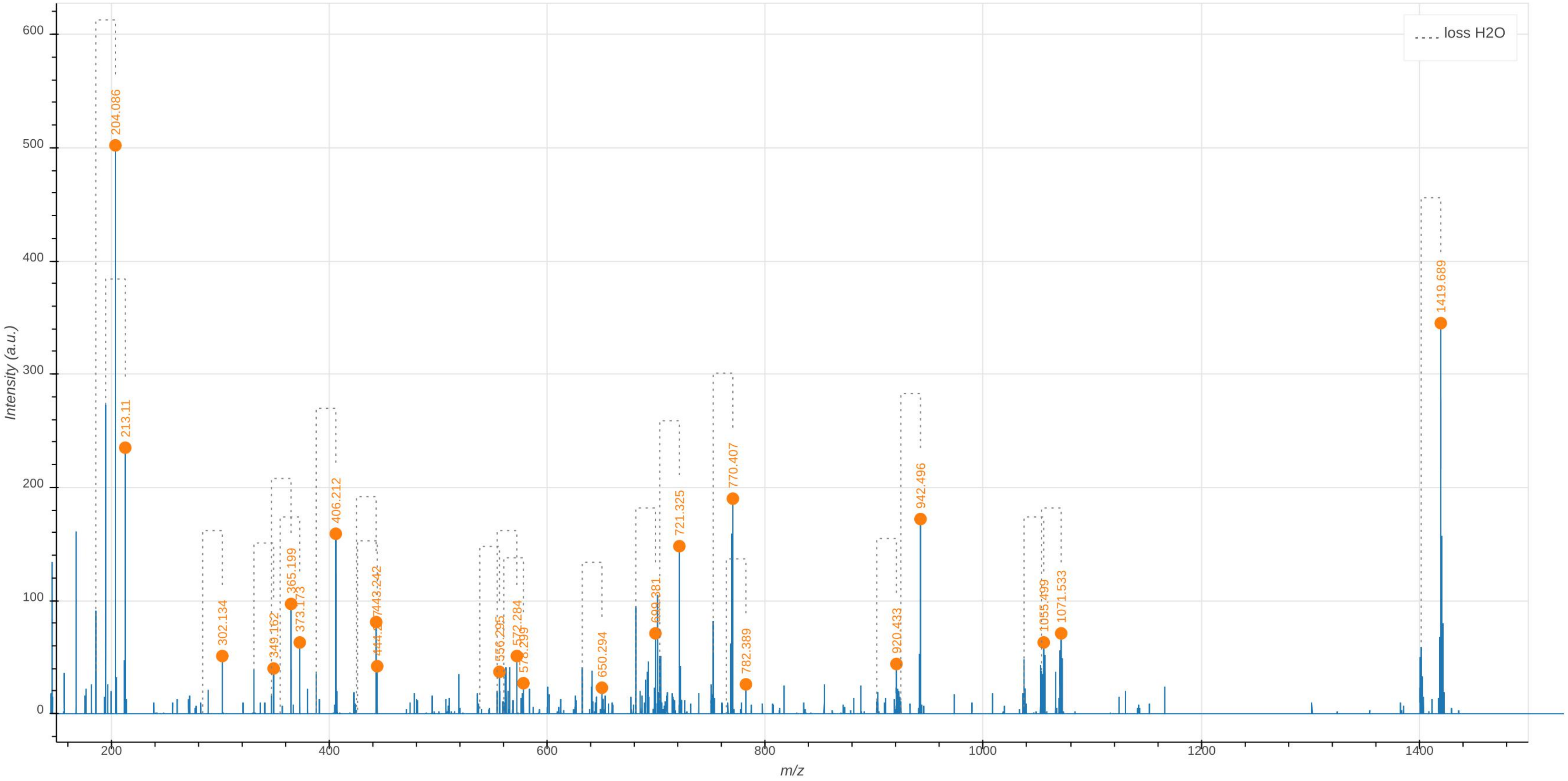

loss H2O
Intensity (a.u.)
m/z
204.086
213.11
302.134
349.162
365.199
373.173
406.212
443.242
444.2
556.295
572.284
578.299
650.294
699.381
721.325
770.407
782.389
920.433
942.496
1055.499
1071.533
1419.689

### Tri-Tetra

**Supplementary Data 5.1: Tandem mass spectrometry analysis of the uniformly labeled Tri(4→3)Tetra dimer.** The observed $m/z_{\text{obs}}$ and calculated $m/z_{\text{cal}}$ values of the parental ion, $[M+2H]^{2+}$, as determined in the absence of fragmentation (MS1), are indicated. The $m/z_{\text{calc}}$ value was used to select the ion for fragmentation. The difference between the $m/z_{\text{obs}}$ and $m/z_{\text{calc}}$ values is indicated in ppm. The GlcNAc residue is indicated in two moieties, glucosamine (GlcN) and the acetyl group (Ac), since these moieties are potentially differentially labeled. For the same reason, the reduced (Red) MurNAc residue is indicated in three moieties, reduced glucosamine ($GlcN^{Red}$), the acetyl group (Ac), and the D-lactoyl group. Labeled and unlabeled moieties are represented in red and purple, respectively. The peaks corresponding to the list of fragments appearing in the table are highlighted by orange dots in the mass spectrum. In the mass spectrum, peaks differing by the loss of $H_20$ are connected by a dashed line. The tables only contain the $m/z$ values for the fragments containing $H_20$. The arrows indicate the position of the 3→3 and 4→3 crosslinks. Residues in the acceptor are underlined.
An interactive report of the MS2 analysis is available in Supplementary File F5.1.

| Precursor ion (MS1) | $m/z_{obs}$ $[M+2H]^{2+}$ | $m/z_{calc}$ $[M+2H]^{2+}$ | ppm |
|---|---|---|---|
| GlcN(-Ac)-GlcN^Red(-Ac)-Lac-Ala-Glu-DAP→DAP(-Ala)-Glu-Ala-Lac-GlcN^Red(-Ac)-GlcN(-Ac) | 939.988 | 939.992 | -3.3 |

| Product ion | $m/z_{obs}$ $[M+H]^{1+}$ | $m/z_{calc}$ $[M+H]^{1+}$ | ppm | Intensity (a.u.) |
|---|---|---|---|---|
| GlcN^Red(-Ac)-Lac-Ala-Glu-DAP→DAP(-Ala)-Glu-Ala-Lac-GlcN^Red(-Ac) | 1454.757 | 1454.769 | 8.0 | 376 |
| GlcN^Red(-Ac)-Lac-Ala-Glu-DAP→DAP(-Ala)-Glu-Ala | 1165.632 | 1165.619 | -11.3 | 41 |
| GlcN^Red(-Ac)-Lac-Ala-Glu-DAP→DAP(-Ala)-Glu | 1090.586 | 1090.574 | -10.9 | 238 |
| GlcN^Red(-Ac)-Lac-Ala-Glu-DAP→DAP-Glu-Ala | 1072.561 | 1072.564 | 2.6 | 158 |
| GlcN^Red(-Ac)-Lac-Ala-Glu-DAP→DAP-Glu | 997.520 | 997.520 | -0.5 | 97 |
| GlcN^Red(-Ac)-Lac-Ala-Glu-DAP→DAP(-Ala) | 955.528 | 955.518 | -10.1 | 135 |
| DAP(-Ala)-Glu-Ala-Lac-GlcN^Red(-Ac) | 774.414 | 774.416 | 2.5 | 184 |
| Glu-DAP→DAP(-Ala) | 591.320 | 591.324 | 6.3 | 44 |
| GlcN^Red(-Ac)-Lac-Ala | 365.201 | 365.202 | 1.9 | 135 |
| Glu-DAP | 317.166 | 317.167 | 3.2 | 123 |

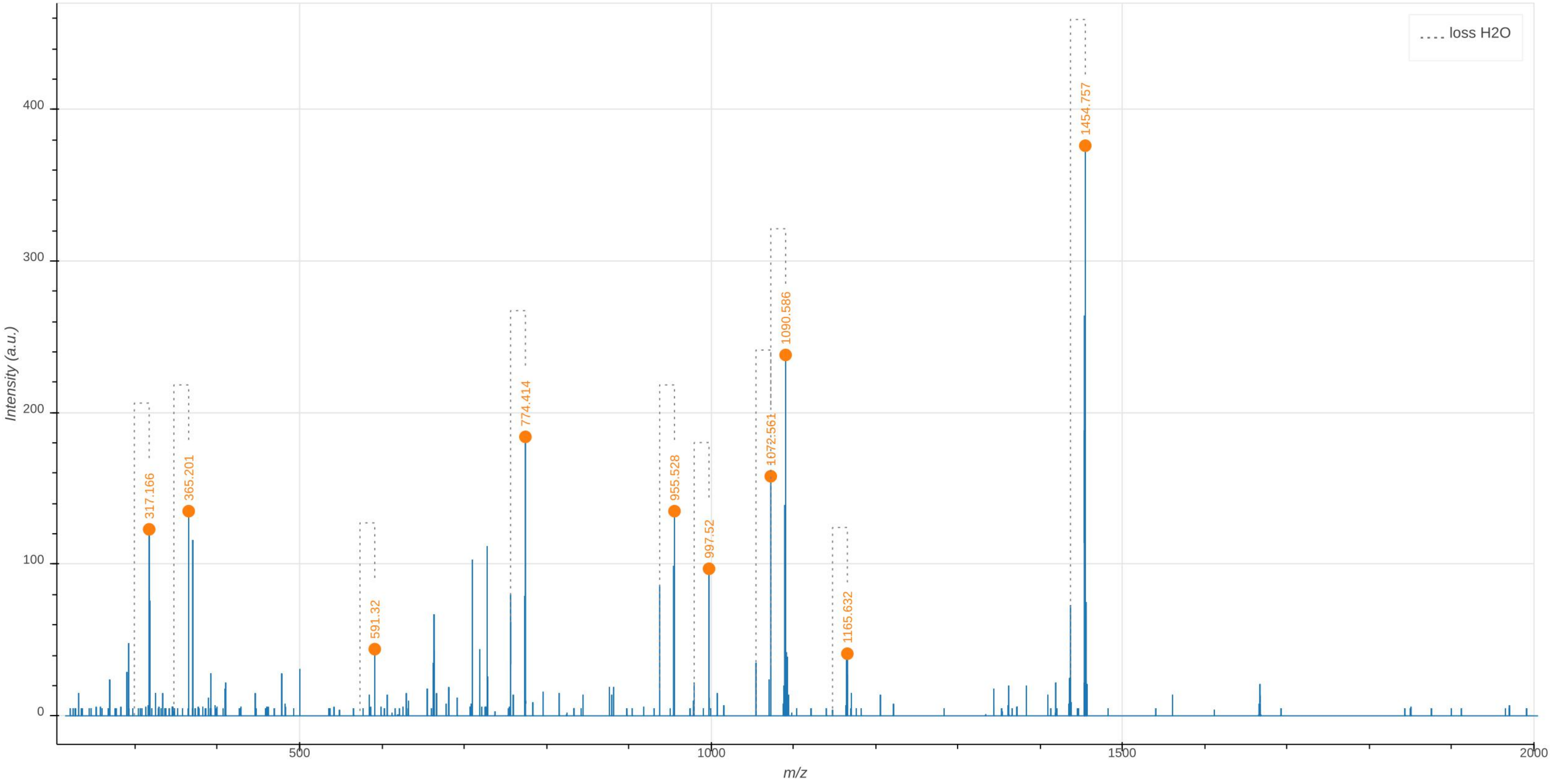

loss H2O
Intensity (a.u.)
m/z
0
100
200
300
400
500
1000
1500
2000
317.166
365.201
591.32
774.414
955.528
997.52
1072.561
1090.586
1165.632
1454.757

**Supplementary Data 5.2: Tandem mass spectrometry analysis of the uniformly unlabeled Tri(3→3)Tetra dimer.** The observed $m/z_{\text{obs}}$ and calculated $m/z_{\text{cal}}$ values of the parental ion, $[\text{M+2H}]^{2+}$, as determined in the absence of fragmentation (MS1), are indicated. The $m/z_{\text{calc}}$ value was used to select the ion for fragmentation. The difference between the $m/z_{\text{obs}}$ and $m/z_{\text{calc}}$ values is indicated in ppm. The GlcNAc residue is indicated in two moieties, glucosamine (GlcN) and the acetyl group (Ac), since these moieties are potentially differentially labeled. For the same reason, the reduced (Red) MurNAc residue is indicated in three moieties, reduced glucosamine (GlcN$^{\text{Red}}$), the acetyl group (Ac), and the D-lactoyl group. Labeled and unlabeled moieties are represented in red and purple, respectively. The peaks corresponding to the list of fragments appearing in the table are highlighted by orange dots in the mass spectrum. In the mass spectrum, peaks differing by the loss of $H_20$ are connected by a dashed line. The tables only contain the $m/z$ values for the fragments containing $H_20$.
The arrows indicate the position of the 3→3 and 4→3 crosslinks. Residues in the acceptor are underlined.
An interactive report of the MS2 analysis is available in Supplementary File F5.2.

| Precursor ion (MS1) | $m/z_{obs}$ [M+2H]$^{2+}$ | $m/z_{calc}$ [M+2H]$^{2+}$ | ppm |
|---|---|---|---|
| GlcN(-Ac)-GlcN$^{Red}$(-Ac)-Lac-Ala-Glu-DAP→DAP(-Ala)-Glu-Ala-Lac-GlcN$^{Red}$(-Ac)-GlcN(-Ac) | 897.891 | 897.892 | -0.9 |

| Product ion | $m/z_{obs}$ [M+H]$^{1+}$ | $m/z_{calc}$ [M+H]$^{1+}$ | ppm | Intensity (a.u.) |
|---|---|---|---|---|
| GlcN(-Ac)-GlcN$^{Red}$(-Ac)-Lac-Ala-Glu-DAP→DAP(-Ala)-Glu-Ala-Lac-GlcN$^{Red}$(-Ac) | 1591.703 | 1591.696 | -4.5 | 290 |
| GlcN$^{Red}$(-Ac)-Lac-Ala-Glu-DAP→DAP(-Ala)-Glu-Ala-Lac-GlcN$^{Red}$(-Ac) | 1388.618 | 1388.617 | -0.8 | 2966 |
| GlcN$^{Red}$(-Ac)-Lac-Ala-Glu-DAP→DAP-Glu-Ala-Lac-GlcN$^{Red}$(-Ac) | 1299.575 | 1299.569 | -4.2 | 311 |
| GlcN$^{Red}$(-Ac)-Lac-Ala-Glu-DAP→DAP(-Ala)-Glu-Ala-Lac | 1183.529 | 1183.522 | -5.7 | 139 |
| GlcN$^{Red}$(-Ac)-Lac-Ala-Glu-DAP→DAP(-Ala)-Glu-Ala | 1111.500 | 1111.501 | 0.4 | 614 |
| GlcN$^{Red}$(-Ac)-Lac-Ala-Glu-DAP→DAP-Glu-Ala-Lac | 1094.502 | 1094.474 | -25.7 | 132 |
| GlcN$^{Red}$(-Ac)-Lac-Ala-Glu-DAP→DAP(-Ala)-Glu | 1040.465 | 1040.464 | -1.8 | 1697 |
| GlcN$^{Red}$(-Ac)-Lac-Ala-Glu-DAP→DAP-Glu-Ala | 1022.450 | 1022.453 | 2.8 | 687 |
| GlcN$^{Red}$(-Ac)-Lac-Ala-Glu-DAP→DAP-Glu | 951.412 | 951.416 | 4.5 | 820 |
| GlcN$^{Red}$(-Ac)-Lac-Ala-Glu-DAP→DAP(-Ala) | 911.422 | 911.421 | -0.9 | 805 |
| Lac-Ala-Glu-DAP→DAP(-Ala)-Glu-Ala | 906.399 | 906.406 | 7.8 | 112 |
| GlcN$^{Red}$(-Ac)-Lac-Ala-Glu-DAP→DAP | 822.372 | 822.373 | 1.4 | 141 |
| Ala-Glu-DAP→DAP(-Ala)-Glu | 763.350 | 763.347 | -3.4 | 207 |
| DAP(-Ala)-Glu-Ala-Lac-GlcN$^{Red}$(-Ac) | 739.335 | 739.336 | 0.9 | 1449 |
| Glu-DAP→DAP(-Ala)-Glu | 692.308 | 692.310 | 3.2 | 351 |
| Ala-Glu-DAP→DAP-Glu | 674.295 | 674.300 | 6.8 | 209 |
| GlcN$^{Red}$(-Ac)-Lac-Ala-Glu-DAP | 650.291 | 650.288 | -3.1 | 639 |
| Ala-Glu-DAP→DAP(-Ala) | 634.304 | 634.305 | 1.8 | 231 |
| Glu-DAP→DAP-Glu | 603.267 | 603.263 | -6.7 | 231 |
| Glu-DAP→DAP(-Ala) | 563.270 | 563.268 | -4.0 | 605 |
| Ala-Glu-DAP→DAP | 545.260 | 545.257 | -4.6 | 275 |
| GlcN$^{Red}$(-Ac)-Lac-Ala-Glu | 478.201 | 478.204 | 5.9 | 120 |
| Glu-DAP→DAP | 474.220 | 474.220 | -1.0 | 198 |
| DAP→DAP(-Ala) | 434.225 | 434.225 | -0.2 | 182 |
| DAP(-Ala)-Glu | 391.180 | 391.183 | 7.9 | 360 |
| Ala-Glu-DAP | 373.173 | 373.172 | -2.4 | 360 |
| GlcN$^{Red}$(-Ac)-Lac-Ala | 349.160 | 349.161 | 3.2 | 653 |
| Glu-DAP | 302.135 | 302.135 | -0.7 | 1087 |
| GlcN$^{Red}$(-Ac)-Lac | 278.124 | 278.124 | 0.3 | 167 |

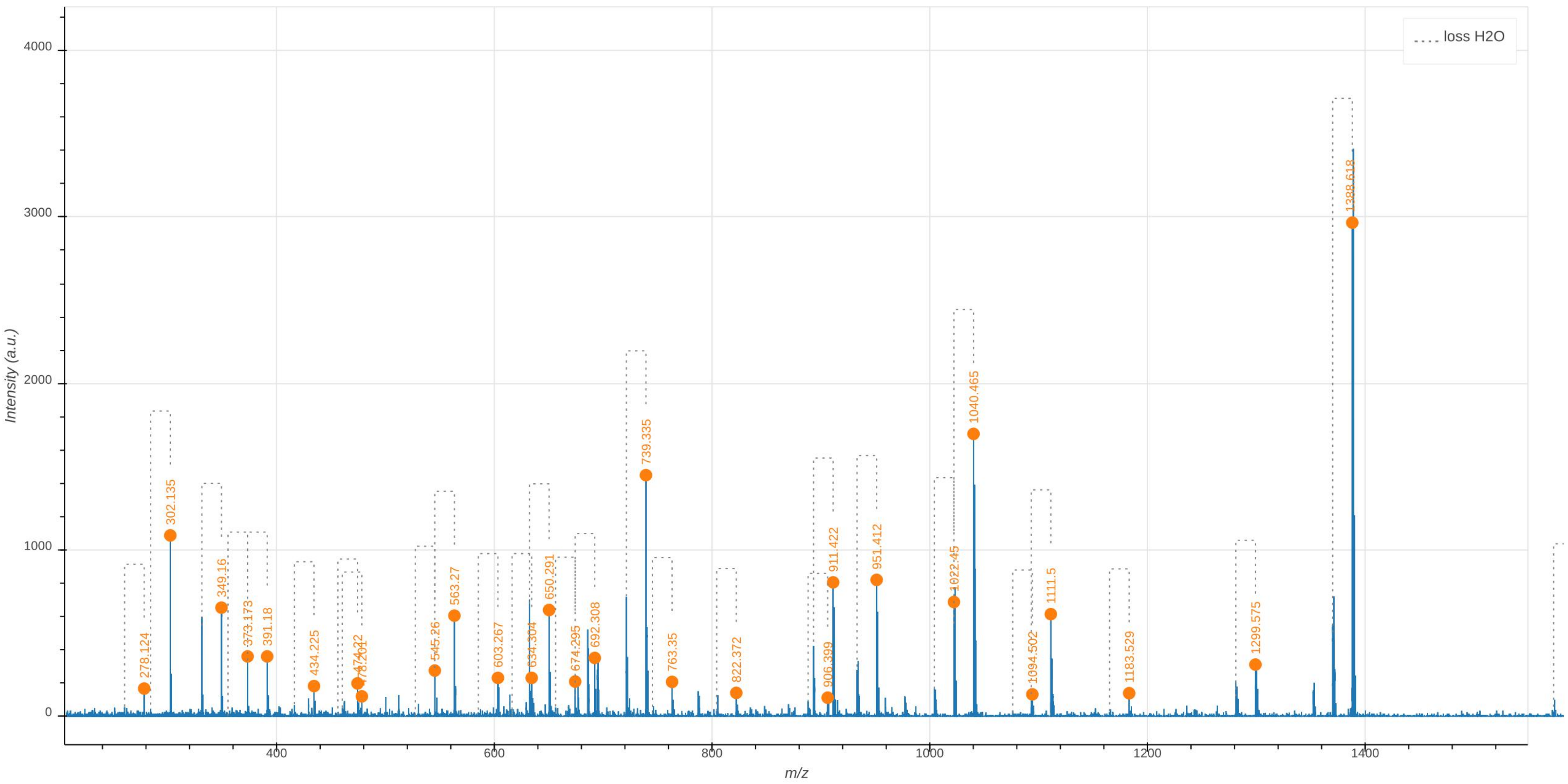
loss H2O
Intensity (a.u.)
m/z
0
1000
2000
3000
4000
400
600
800
1000
1200
1400
278.124
302.135
349.16
373.173
391.18
434.225
474.22
478.201
545.26
563.27
603.267
634.304
650.291
674.295
692.308
739.335
763.35
822.372
906.399
911.422
951.412
1022.45
1040.465
1094.502
1111.5
1183.529
1299.575
1388.618

**Supplementary Data 5.3: Tandem mass spectrometry analysis of the Tri→Tetra dimer containing a uniformly labeled disaccharide-tetrapeptide cross-linked to a uniformly unlabeled disaccharide-tripeptide (3→3 cross-link).** The observed $m/z_{\mathrm{obs}}$ and calculated $m/z_{\mathrm{cal}}$ values of the parental ion, $[M{+}2H]^{2+}$, as determined in the absence of fragmentation (MS1), are indicated. The $m/z_{\mathrm{calc}}$ value was used to select the ion for fragmentation. The difference between the $m/z_{\mathrm{obs}}$ and $m/z_{\mathrm{calc}}$ values is indicated in ppm. The GlcNAc residue is indicated in two moieties, glucosamine (GlcN) and the acetyl group (Ac), since these moieties are potentially differentially labeled. For the same reason, the reduced (Red) MurNAc residue is indicated in three moieties, reduced glucosamine ($\mathrm{GlcN^{Red}}$), the acetyl group (Ac), and the D-lactoyl group. Labeled and unlabeled moieties are represented in red and purple, respectively.

In the mass spectrum, peaks differing by the loss of $H_20$ are connected by a dashed line. The arrows indicate the position of the 3→3 and 4→3 crosslinks. Residues in the acceptor are underlined.

An interactive report of the MS2 analysis is available in Supplementary File F5.3.

| Precursor ion (MS1) | $m/z_{obs}$ $[M{+}2H]^{2+}$ | $m/z_{calc}$ $[M{+}2H]^{2+}$ | ppm |
|---|---|---|---|
| GlcN(-Ac)-$\mathrm{GlcN^{Red}}$(-Ac)-Lac-Ala-Glu-DAP→DAP(-Ala)-Glu-Ala-Lac-$\mathrm{GlcN^{Red}}$(-Ac)-GlcN(-Ac) | 919.942 | 919.943 | -1.0 |

| Product ion | $m/z_{obs}$ $[M{+}H]^{1+}$ | $m/z_{calc}$ $[M{+}H]^{1+}$ | ppm | Intensity (a.u.) |
|---|---|---|---|---|
| $\mathrm{GlcN^{Red}}$(-Ac)-Lac-Ala-Glu-DAP→DAP(-Ala)-Glu-Ala-Lac-$\mathrm{GlcN^{Red}}$(-Ac) | 1423.693 | 1423.696 | 2.4 | 167 |
| $\mathrm{GlcN^{Red}}$(-Ac)-Lac-Ala-Glu-DAP→DAP-Glu-Ala-Lac-$\mathrm{GlcN^{Red}}$(-Ac) | 1330.655 | 1330.642 | -10.1 | 30 |
| Glu-DAP→DAP(-Ala)-Glu-Ala-Lac-$\mathrm{GlcN^{Red}}$(-Ac) | 1075.541 | 1075.543 | 2.3 | 41 |
| $\mathrm{GlcN^{Red}}$(-Ac)-Lac-Ala-Glu-DAP→DAP(-Ala)-Glu | 1059.504 | 1059.502 | -1.7 | 58 |
| Glu-DAP→DAP-Glu-Ala-Lac-$\mathrm{GlcN^{Red}}$(-Ac) | 982.486 | 982.488 | 2.8 | 13 |
| DAP→DAP(-Ala)-Glu-Ala-Lac-$\mathrm{GlcN^{Red}}$(-Ac) | 946.496 | 946.500 | 5.1 | 37 |
| $\mathrm{GlcN^{Red}}$(-Ac)-Lac-Ala-Glu-DAP→DAP(-Ala) | 924.445 | 924.446 | 1.1 | 57 |
| DAP(-Ala)-Glu-Ala-Lac-$\mathrm{GlcN^{Red}}$(-Ac) | 774.421 | 774.416 | -7.2 | 105 |
| Glu-Ala-Lac-$\mathrm{GlcN^{Red}}$(-Ac)-GlcN(-Ac) | 712.348 | 712.362 | 19.6 | 17 |
| DAP-Glu-Ala-Lac-$\mathrm{GlcN^{Red}}$(-Ac) | 681.354 | 681.361 | 10.6 | 19 |
| Glu-DAP→DAP-Glu | 618.291 | 618.294 | 4.1 | 29 |
| DAP→DAP-Glu | 489.249 | 489.251 | 5.0 | 15 |
| DAP→DAP(-Ala) | 447.248 | 447.250 | 4.1 | 15 |
| DAP-Glu-Ala | 392.206 | 392.211 | 13.2 | 40 |
| $\mathrm{GlcN^{Red}}$(-Ac)-Lac-Ala | 349.157 | 349.161 | 10.7 | 18 |
| DAP-Glu | 317.170 | 317.167 | -9.7 | 15 |
| GlcN(-Ac) | 213.110 | 213.111 | 5.5 | 122 |
| GlcN(-Ac) | 204.086 | 204.087 | 6.8 | 471 |
| DAP | 182.110 | 182.110 | 0.8 | 24 |
| Ala-Lac | 151.083 | 151.083 | -1.5 | 40 |

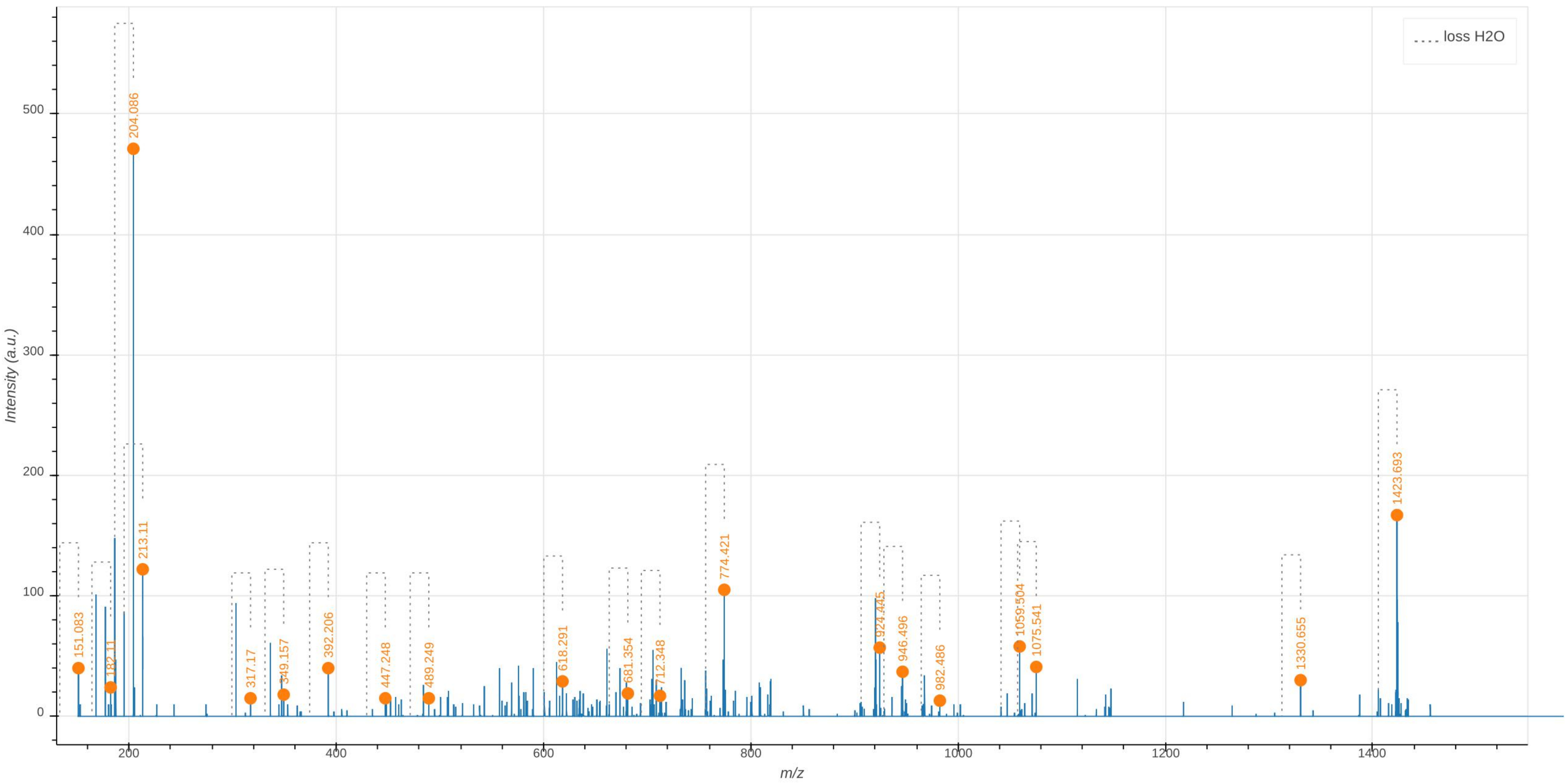

loss H2O
Intensity (a.u.)
m/z
204.086
213.11
151.083
182.11
317.17
349.157
392.206
447.248
489.249
618.291
681.354
712.348
774.421
924.445
946.496
982.486
1059.504
1075.541
1330.655
1423.693

**Tetra-Tetra**

**Supplementary Data 6.1: Tandem mass spectrometry analysis of the uniformly labeled Tetra(4→3)Tetra dimer.** The observed $m/z_{\text{obs}}$ and calculated $m/z_{\text{cal}}$ values of the parental ion, $[\text{M+2H}]^{2+}$, as determined in the absence of fragmentation (MS1), are indicated. The $m/z_{\text{calc}}$ value was used to select the ion for fragmentation. The difference between the $m/z_{\text{obs}}$ and $m/z_{\text{calc}}$ values is indicated in ppm. The GlcNAc residue is indicated in two moieties, glucosamine (GlcN) and the acetyl group (Ac), since these moieties are potentially differentially labeled. For the same reason, the reduced (Red) MurNAc residue is indicated in three moieties, reduced glucosamine ($\text{GlcN}^{\text{Red}}$), the acetyl group (Ac), and the D-lactoyl group. Labeled and unlabeled moieties are represented in red and purple, respectively. The peaks corresponding to the list of fragments appearing in the table are highlighted by orange dots in the mass spectrum. In the mass spectrum, peaks differing by the loss of $H_20$ are connected by a dashed line. The tables only contain the $m/z$ values for the fragments containing $H_20$. The arrows indicate the position of the 3→3 and 4→3 crosslinks. Residues in the acceptor are underlined.
An interactive report of the MS2 analysis is available in Supplementary File F6.1.

| Precursor ion (MS1) | $m/z_{obs}$ [M+2H]$^{2+}$ | $m/z_{calc}$ [M+2H]$^{2+}$ | ppm |
|---|---|---|---|
| GlcN(-Ac)-GlcN$^{\text{Red}}$(-Ac)-Lac-Ala-Glu-DAP-Ala→DAP(-Ala)-Glu-Ala-Lac-GlcN$^{\text{Red}}$(-Ac)-GlcN(-Ac) | 977.512 | 977.514 | -2.0 |

| Product ion | $m/z_{obs}$ [M+H]$^{1+}$ | $m/z_{calc}$ [M+H]$^{1+}$ | ppm | Intensity (a.u.) |
|---|---|---|---|---|
| GlcN$^{\text{Red}}$(-Ac)-Lac-Ala-Glu-DAP-Ala→DAP(-Ala)-Glu-Ala-Lac-GlcN$^{\text{Red}}$(-Ac) | 1529.815 | 1529.813 | -1.7 | 1017 |
| GlcN$^{\text{Red}}$(-Ac)-Lac-Ala-Glu-DAP-Ala→DAP(-Ala)-Glu-Ala | 1240.686 | 1240.663 | -18.5 | 172 |
| GlcN$^{\text{Red}}$(-Ac)-Lac-Ala-Glu-DAP-Ala→DAP(-Ala)-Glu | 1165.619 | 1165.619 | -0.4 | 576 |
| GlcN$^{\text{Red}}$(-Ac)-Lac-Ala-Glu-DAP-Ala→DAP-Glu-Ala | 1147.614 | 1147.608 | -5.1 | 229 |
| GlcN$^{\text{Red}}$(-Ac)-Lac-Ala-Glu-DAP-Ala→DAP-Glu | 1072.569 | 1072.564 | -4.8 | 156 |
| GlcN$^{\text{Red}}$(-Ac)-Lac-Ala-Glu-DAP-Ala→DAP(-Ala) | 1030.565 | 1030.562 | -2.9 | 194 |
| Ala→DAP(-Ala)-Glu-Ala-Lac-GlcN$^{\text{Red}}$(-Ac) | 849.462 | 849.460 | -2.4 | 405 |
| DAP(-Ala)-Glu-Ala-Lac-GlcN$^{\text{Red}}$(-Ac) | 774.421 | 774.416 | -7.4 | 274 |
| GlcN$^{\text{Red}}$(-Ac)-Lac-Ala-Glu-DAP-Ala | 756.411 | 756.405 | -8.2 | 533 |
| Glu-DAP-Ala→DAP-Glu | 708.373 | 708.370 | -5.4 | 106 |
| GlcN$^{\text{Red}}$(-Ac)-Lac-Ala | 365.203 | 365.202 | -1.1 | 320 |
| Glu-DAP | 317.168 | 317.167 | -4.8 | 197 |
| GlcN(-Ac) | 213.112 | 213.111 | -6.5 | 2182 |
| DAP | 182.113 | 182.110 | -14.4 | 253 |
| Lac-Ala | 151.086 | 151.083 | -19.2 | 414 |

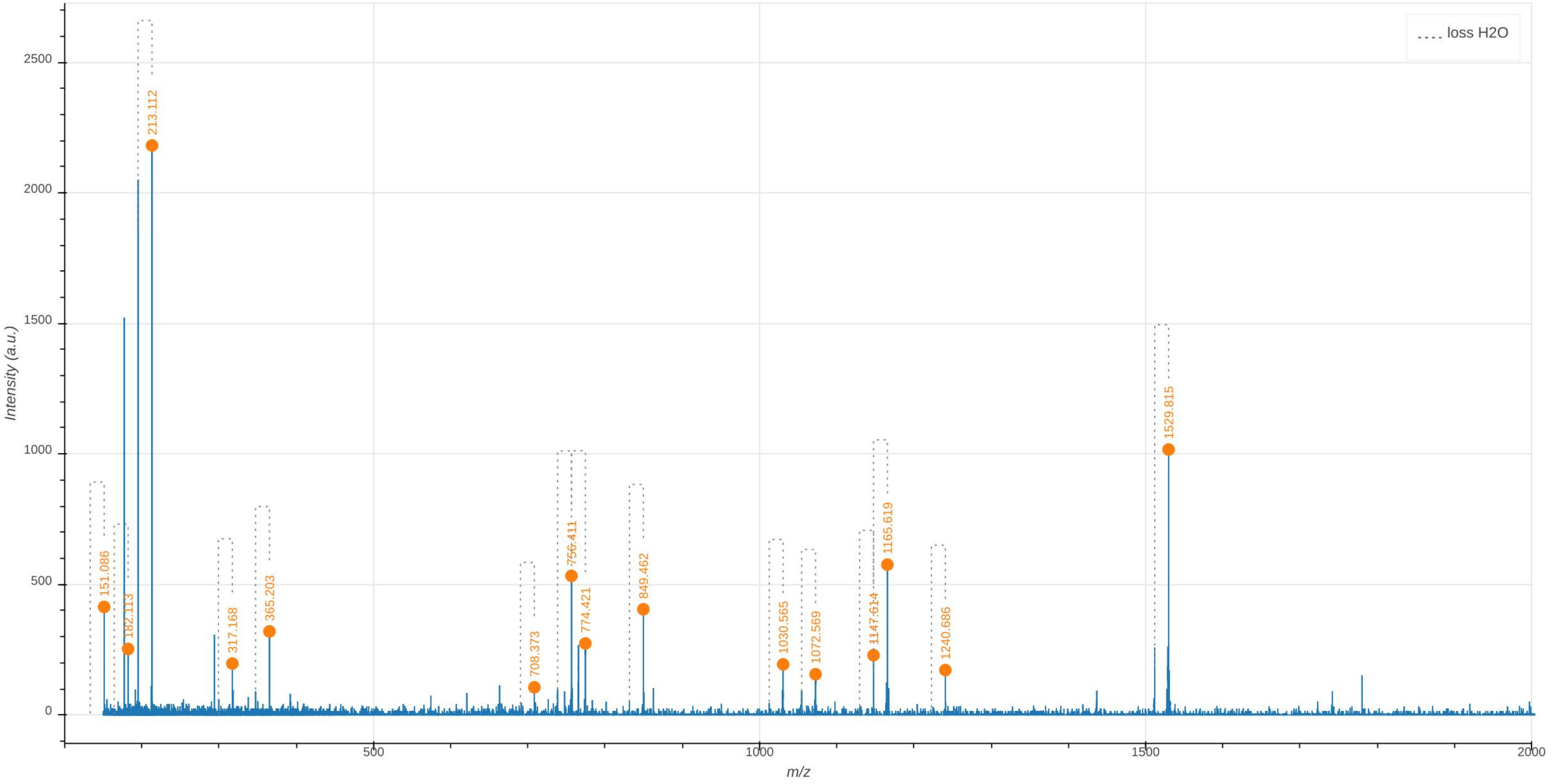

loss H2O
Intensity (a.u.)
m/z
0
500
1000
1500
2000
2500
151.086
182.113
213.112
317.168
365.203
708.373
756.411
774.421
849.462
1030.565
1072.569
1147.614
1165.619
1240.686
1529.815

**Supplementary Data 6.2: Tandem mass spectrometry analysis of the uniformly unlabeled Tetra(4→3)Tetra dimer.** The observed $m/z_{\text{obs}}$ and calculated $m/z_{\text{cal}}$ values of the parental ion, $[M+2H]^{2+}$, as determined in the absence of fragmentation (MS1), are indicated. The $m/z_{\text{calc}}$ value was used to select the ion for fragmentation. The difference between the $m/z_{\text{obs}}$ and $m/z_{\text{calc}}$ values is indicated in ppm. The GlcNAc residue is indicated in two moieties, glucosamine (GlcN) and the acetyl group (Ac), since these moieties are potentially differentially labeled. For the same reason, the reduced (Red) MurNAc residue is indicated in three moieties, reduced glucosamine (GlcN$^{\text{Red}}$), the acetyl group (Ac), and the D-lactoyl group. Labeled and unlabeled moieties are represented in red and purple, respectively. The peaks corresponding to the list of fragments appearing in the table are highlighted by orange dots in the mass spectrum. In the mass spectrum, peaks differing by the loss of $H_20$ are connected by a dashed line. The tables only contain the $m/z$ values for the fragments containing $H_20$.
The arrows indicate the position of the 3→3 and 4→3 crosslinks. Residues in the acceptor are underlined.
An interactive report of the MS2 analysis is available in Supplementary File F6.2.

| Precursor ion (MS1) | $m/z_{obs}$ $[M+2H]^{2+}$ | $m/z_{calc}$ $[M+2H]^{2+}$ | ppm |
|---|---|---|---|
| GlcN(-Ac)-GlcN$^{Red}$(-Ac)-Lac-Ala-Glu-DAP-Ala→DAP(-Ala)-Glu-Ala-Lac-GlcN$^{Red}$(-Ac)-GlcN(-Ac) | 933.409 | 933.410 | -1.7 |

| Product ion | $m/z_{obs}$ $[M+H]^{1+}$ | $m/z_{calc}$ $[M+H]^{1+}$ | ppm | Intensity (a.u.) |
|---|---|---|---|---|
| GlcN(-Ac)-GlcN$^{Red}$(-Ac)-Lac-Ala-Glu-DAP-Ala→DAP(-Ala)-Glu-Ala-Lac-GlcN$^{Red}$(-Ac) | 1662.731 | 1662.733 | 1.1 | 416 |
| GlcN$^{Red}$(-Ac)-Lac-Ala-Glu-DAP-Ala→DAP(-Ala)-Glu-Ala-Lac-GlcN$^{Red}$(-Ac) | 1459.655 | 1459.654 | -0.9 | 4041 |
| GlcN$^{Red}$(-Ac)-Lac-Ala-Glu-DAP-Ala→DAP-Glu-Ala-Lac-GlcN$^{Red}$(-Ac) | 1370.614 | 1370.606 | -5.8 | 419 |
| GlcN(-Ac)-GlcN$^{Red}$(-Ac)-Lac-Ala-Glu-DAP-Ala→DAP(-Ala)-Glu | 1314.586 | 1314.580 | -4.8 | 108 |
| GlcN$^{Red}$(-Ac)-Lac-Ala-Glu-DAP-Ala→DAP(-Ala)-Glu-Ala | 1182.533 | 1182.538 | 4.2 | 436 |
| GlcN$^{Red}$(-Ac)-Lac-Ala-Glu-DAP-Ala→DAP(-Ala)-Glu | 1111.500 | 1111.501 | 0.3 | 2489 |
| GlcN$^{Red}$(-Ac)-Lac-Ala-Glu-DAP-Ala→DAP-Glu-Ala | 1093.492 | 1093.490 | -1.8 | 1176 |
| GlcN$^{Red}$(-Ac)-Lac-Ala-Glu-DAP-Ala→DAP-Glu | 1022.450 | 1022.453 | 3.2 | 1094 |
| GlcN$^{Red}$(-Ac)-Lac-Ala-Glu-DAP-Ala→DAP(-Ala) | 982.461 | 982.458 | -2.7 | 1305 |
| GlcN$^{Red}$(-Ac)-Lac-Ala-Glu-DAP-Ala→DAP | 893.408 | 893.410 | 3.2 | 159 |
| Ala-Glu-DAP-Ala→DAP(-Ala)-Glu | 834.397 | 834.385 | -14.7 | 173 |
| Ala→DAP(-Ala)-Glu-Ala-Lac-GlcN$^{Red}$(-Ac) | 810.373 | 810.373 | 0.4 | 2603 |
| Glu-DAP-Ala→DAP(-Ala)-Glu | 763.353 | 763.347 | -7.3 | 355 |
| Ala-Glu-DAP-Ala→DAP-Glu | 745.337 | 745.337 | 0.4 | 285 |
| DAP(-Ala)-Glu-Ala-Lac-GlcN$^{Red}$(-Ac) | 739.336 | 739.336 | 0.8 | 1027 |
| GlcN$^{Red}$(-Ac)-Lac-Ala-Glu-DAP-Ala | 721.328 | 721.326 | -3.7 | 2676 |
| Ala-Glu-DAP-Ala→DAP(-Ala) | 705.345 | 705.342 | -3.8 | 134 |
| Glu-DAP-Ala→DAP-Glu | 674.300 | 674.300 | 0.1 | 133 |
| GlcN$^{Red}$(-Ac)-Lac-Ala-Glu-DAP | 650.291 | 650.288 | -3.3 | 234 |
| Glu-DAP-Ala→DAP(-Ala) | 634.304 | 634.305 | 1.6 | 324 |
| Ala-Glu-DAP-Ala→DAP | 616.294 | 616.294 | -0.2 | 193 |
| Glu-DAP-Ala→DAP | 545.257 | 545.257 | 0.6 | 279 |
| Ala→DAP(-Ala)-Glu-Ala | 533.257 | 533.257 | 0.6 | 121 |
| DAP-Ala→DAP(-Ala) | 505.261 | 505.262 | 2.2 | 228 |
| Ala→DAP(-Ala)-Glu | 462.221 | 462.220 | -1.2 | 549 |
| Ala-Glu-DAP-Ala | 444.209 | 444.209 | 0.1 | 198 |
| DAP(-Ala)-Glu | 391.183 | 391.183 | 0.2 | 328 |
| Ala-Glu-DAP | 373.173 | 373.172 | -2.6 | 787 |
| GlcN$^{Red}$(-Ac)-Lac-Ala | 349.162 | 349.161 | -1.2 | 1062 |
| Ala→DAP(-Ala) | 333.177 | 333.177 | 2.6 | 284 |
| Glu-DAP | 302.135 | 302.135 | -0.9 | 560 |
| GlcN$^{Red}$(-Ac)-Lac | 278.121 | 278.124 | 10.8 | 141 |

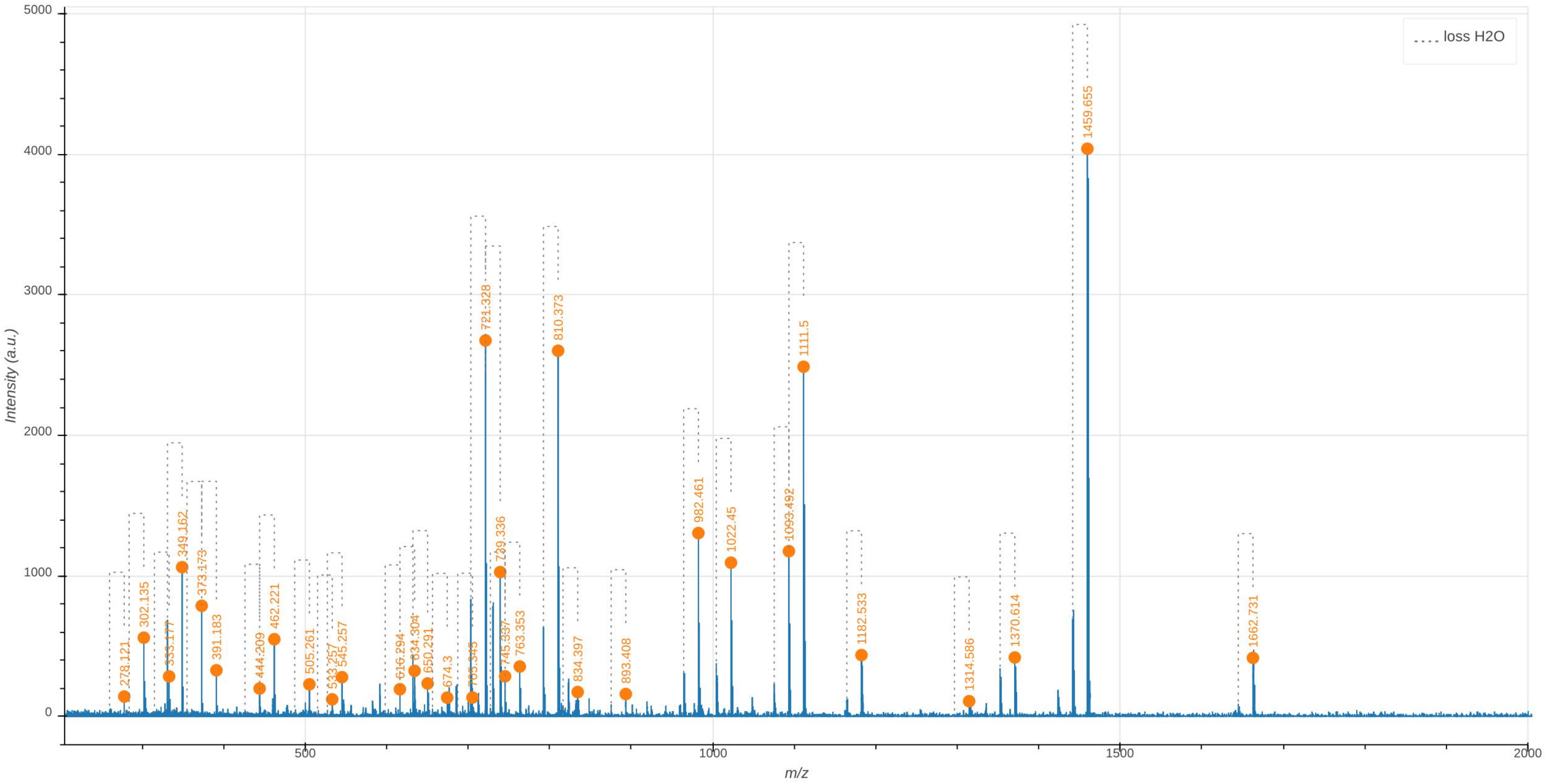

loss H2O
Intensity (a.u.)
m/z
0
1000
2000
3000
4000
5000
500
1000
1500
2000
278.121
302.135
333.177
349.162
373.173
391.183
444.209
462.221
505.261
533.257
545.257
616.294
634.304
650.291
674.3
705.343
721.328
739.336
745.337
763.353
810.373
834.397
893.408
982.461
1022.45
1093.492
1111.5
1182.533
1314.586
1370.614
1459.655
1662.731

**Supplementary Data 6.3: Tandem mass spectrometry analysis of the Tetra→Tetra dimer containing a uniformly labeled disaccharide-tripeptide cross-linked to a uniformly unlabeled disaccharide-tripeptide (4→3 cross-link).** The observed $m/z_{\mathrm{obs}}$ and calculated $m/z_{\mathrm{cal}}$ values of the parental ion, $[\mathrm{M+2H}]^{2+}$, as determined in the absence of fragmentation (MS1), are indicated. The $m/z_{\mathrm{calc}}$ value was used to select the ion for fragmentation. The difference between the $m/z_{\mathrm{obs}}$ and $m/z_{\mathrm{calc}}$ values is indicated in ppm. The GlcNAc residue is indicated in two moieties, glucosamine (GlcN) and the acetyl group (Ac), since these moieties are potentially differentially labeled. For the same reason, the reduced (Red) MurNAc residue is indicated in three moieties, reduced glucosamine ($\mathrm{GlcN}^{\mathrm{Red}}$), the acetyl group (Ac), and the D-lactoyl group. Labeled and unlabeled moieties are represented in red and purple, respectively. The fragmentation potentially leads to two series of fragments (discriminatory fragments) specific of the isotopomers containing the fully labeled disaccharide subunit in the donor (all heavy-all light) or acceptor position (all light-all heavy). These fragments are highlighted by green and orange dots in the mass spectrum, respectively. The other fragments are common to the fragmentation of the two isotopomers (highlighted by blue dots in the mass spectrum). The mass spectrum indicates the presence of the all light-all heavy isotopomers (orange dots). The peak at $m/z_{obs}$ 1128.568 (green dot) can also be accounted for by the loss of $H_20$ from the peak at 1142.567 indicating that all peaks may be accounted for by the presence of the all light-all heavy isotopomer.

In the mass spectrum, peaks differing by the loss of $H_20$ are connected by a dashed line. The arrows indicate the position of the 3→3 and 4→3 crosslinks. Residues in the acceptor are underlined.

An interactive report of the MS2 analysis is available in Supplementary File F6.3.

| Precursor ion (MS1) | $m/z_{obs}$ [M+2H]$^{2+}$ | $m/z_{calc}$ [M+2H]$^{2+}$ | ppm |
|---|---|---|---|
| GlcN(-Ac)-GlcN$^{Red}$(-Ac)-Lac-Ala-Glu-DAP-Ala→DAP(-Ala)-Glu-Ala-Lac-GlcN$^{Red}$(-Ac)-GlcN(-Ac) | 955.459 | 955.462 | -2.6 |
| GlcN(-Ac)-GlcN$^{Red}$(-Ac)-Lac-Ala-Glu-DAP-Ala→DAP(-Ala)-Glu-Ala-Lac-GlcN$^{Red}$(-Ac)-GlcN(-Ac) | 955.459 | 955.462 | -2.6 |

| Discriminatory product ion | $m/z_{obs}$ [M+H]$^{1+}$ | $m/z_{calc}$ [M+H]$^{1+}$ | ppm | Intensity (a.u.) | Isotopologue |
|---|---|---|---|---|---|
| GlcN$^{Red}$(-Ac)-Lac-Ala-Glu-DAP-Ala→DAP-Glu-Ala | 1128.568 | 1128.570 | 1.6 | 215 | all heavy-all light |
| Glu-DAP-Ala→DAP-Glu-Ala-Lac-GlcN$^{Red}$(-Ac) | 1053.516 | 1053.525 | 9.2 | 97 | all light-all heavy |
| GlcN$^{Red}$(-Ac)-Lac-Ala-Glu-DAP-Ala→DAP-Glu | 1037.486 | 1037.484 | -1.5 | 153 | all light-all heavy |
| Ala→DAP(-Ala)-Glu-Ala-Lac-GlcN$^{Red}$(-Ac) | 845.449 | 845.453 | 4.3 | 304 | all light-all heavy |
| DAP(-Ala)-Glu-Ala-Lac-GlcN$^{Red}$(-Ac) | 774.410 | 774.416 | 7.5 | 459 | all light-all heavy |
| Ala→DAP-Glu-Ala-Lac-GlcN$^{Red}$(-Ac) | 752.394 | 752.398 | 5.1 | 307 | all light-all heavy |
| GlcN$^{Red}$(-Ac)-Lac-Ala-Glu-DAP-Ala | 721.322 | 721.326 | 4.4 | 186 | all light-all heavy |
| Glu-DAP-Ala→DAP-Glu | 689.334 | 689.331 | -3.8 | 139 | all light-all heavy |
| DAP(-Ala)-Glu | 410.225 | 410.221 | -9.5 | 158 | all light-all heavy |
| Ala→DAP-Glu | 388.202 | 388.204 | 4.3 | 128 | all light-all heavy |
| Ala→DAP(-Ala) | 346.197 | 346.202 | 14.6 | 106 | all light-all heavy |

| Common product ion | $m/z_{obs}$ [M+H]$^{1+}$ | $m/z_{calc}$ [M+H]$^{1+}$ | ppm | Intensity (a.u.) |
|---|---|---|---|---|
| GlcN$^{Red}$(-Ac)-Lac-Ala-Glu-DAP-Ala→DAP(-Ala)-Glu-Ala-Lac-GlcN$^{Red}$(-Ac) | 1494.729 | 1494.733 | 2.8 | 1109 |
| GlcN$^{Red}$(-Ac)-Lac-Ala-Glu-DAP-Ala→DAP(-Ala)-Glu-Ala-Lac-GlcN$^{Red}$(-Ac) | | | | |
| GlcN$^{Red}$(-Ac)-Lac-Ala-Glu-DAP-Ala→DAP(-Ala)-Glu-Ala | 1205.576 | 1205.583 | 6.2 | 113 |
| Ala-Glu-DAP-Ala→DAP(-Ala)-Glu-Ala-Lac-GlcN$^{Red}$(-Ac) | | | | |
| Glu-DAP-Ala→DAP(-Ala)-Glu-Ala-Lac-GlcN$^{Red}$(-Ac) | 1146.576 | 1146.580 | 3.8 | 295 |
| GlcN$^{Red}$(-Ac)-Lac-Ala-Glu-DAP-Ala→DAP(-Ala)-Glu | | | | |
| Glu-DAP-Ala→DAP(-Ala)-Glu-Ala-Lac-GlcN$^{Red}$(-Ac) | 1130.533 | 1130.539 | 5.7 | 219 |
| GlcN$^{Red}$(-Ac)-Lac-Ala-Glu-DAP-Ala→DAP(-Ala)-Glu | | | | |
| DAP-Ala→DAP(-Ala)-Glu-Ala-Lac-GlcN$^{Red}$(-Ac) | 1017.530 | 1017.538 | 7.3 | 126 |
| GlcN$^{Red}$(-Ac)-Lac-Ala-Glu-DAP-Ala→DAP(-Ala) | | | | |
| DAP-Ala→DAP(-Ala)-Glu-Ala-Lac-GlcN$^{Red}$(-Ac) | 995.475 | 995.483 | 7.4 | 176 |
| GlcN$^{Red}$(-Ac)-Lac-Ala-Glu-DAP-Ala→DAP(-Ala) | | | | |
| DAP-Ala→DAP(-Ala)-Glu | 647.322 | 647.329 | 11.3 | 149 |
| Glu-DAP-Ala→DAP(-Ala) | | | | |
| DAP-Glu | 302.130 | 302.135 | 15.7 | 91 |
| Glu-DAP | | | | |
| GlcN(-Ac) | 204.089 | 204.087 | -9.0 | 2307 |
| GlcN(-Ac) | | | | |
| DAP | 182.108 | 182.110 | 12.0 | 134 |

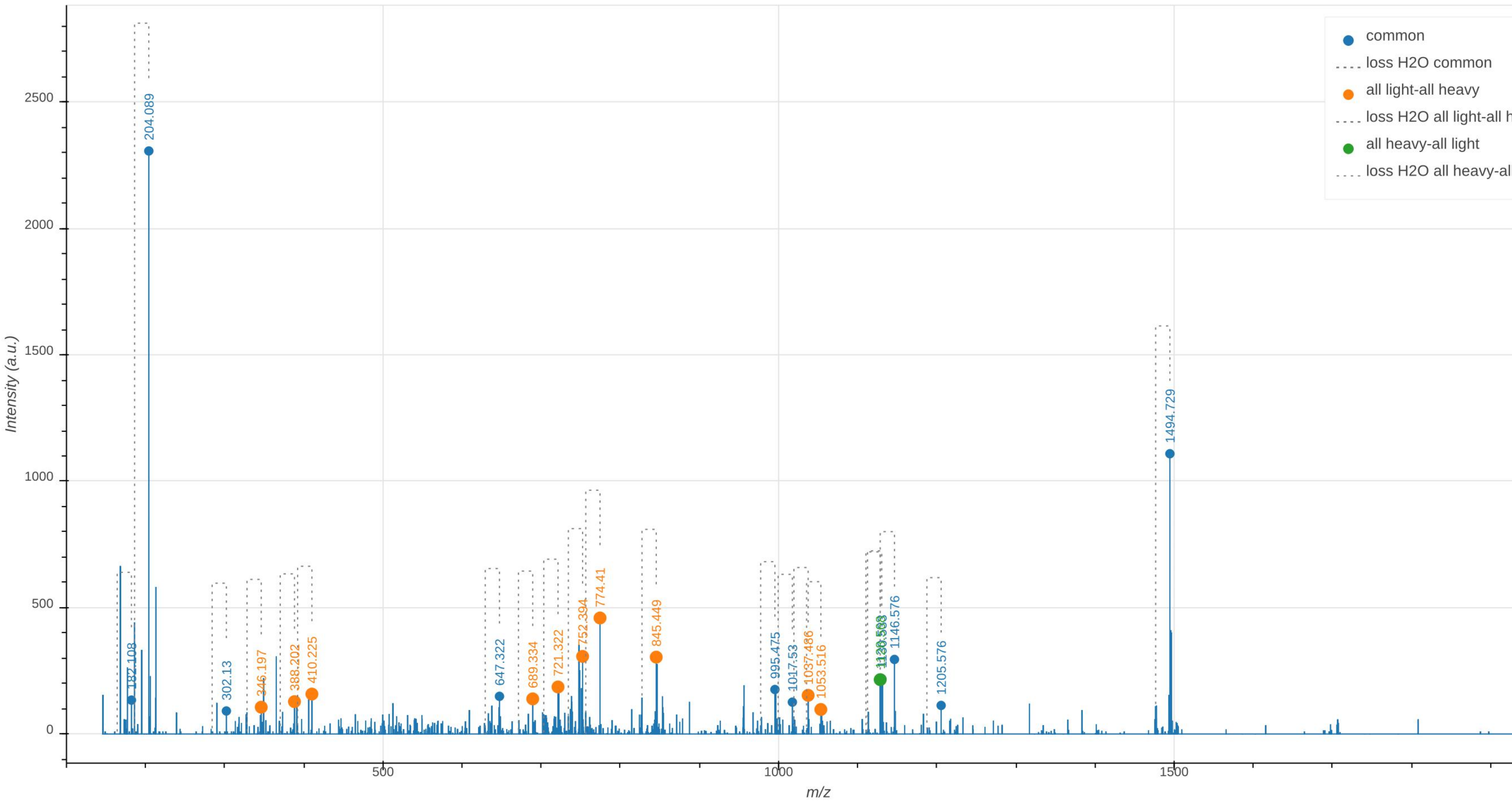

common
loss H2O common
all light-all heavy
loss H2O all light-all h
all heavy-all light
loss H2O all heavy-al
Intensity (a.u.)
m/z
0
500
1000
1500
2000
2500
500
1000
1500
187.108
204.089
302.13
346.197
388.202
410.225
647.322
689.334
721.322
752.394
774.41
845.449
995.475
1017.53
1037.486
1053.516
1146.576
1205.576
1494.729
