## Supplementary Material and Methods for "Heavy isotope labeling and mass spectrometry reveal unexpected remodeling of bacterial cell wall expansion in response to drugs"

### Mass spectrometric data analysis

**Simulation of high resolution mass spectra to determine the isotopic composition of muropeptides.** Mass spectral data analyses were carried over by performing matches between simulated and observed isotopic clusters for muropeptides recovered from bacteria grown in labeled and unlabeled media. The approach used to generate the simulated spectra was adapted from a recent publication (Lacki et al., 2017), as detailed below for the reduced disaccharide-tripeptide (GlcNAc-MurNAc-L-Ala-γ-D-Glu-DAP), which was taken as an example.

**Table 1 of Supplementary Material and Methods**

**Mass and relative abundance of H, C, N, and O isotopes used for isotopic cluster simulations**

| | | Relative abundance in the growth media | |
|---|---|---|---|
| **Isotope** | **Mass** [a] | **Unlabeled** | **Labeled** |
| $^{12}C$ | 12.000 000 000 000 | 0.9893 [b] | 0.01 [c] |
| $^{13}C$ | 13.003 354 835 071 | 0.0107 [b] | 0.99 [c] |
| $^{1}H$ | 1.007 825 032 231 | 0.999885 [b] | 0.999885 [b] |
| $^{2}H$ | 2.014 101 778 120 | 0.000115 [b] | 0.000115 [b] |
| $^{14}N$ | 14.003 074 004 426 | 0.99636 [b] | 0.01 [c] |
| $^{15}N$ | 15.000 108 898 884 | 0.00364 [b] | 0.99 [c] |
| $^{16}O$ | 15.994 914 619 566 | 0.99757 [b] | 0.99757 [b] |
| $^{17}O$ | 16.999 131 756 500 | 0.00038 [b] | 0.00038 [b] |
| $^{18}O$ | 17.999 159 612 858 | 0.00205 [b] | 0.00205 [b] |

[a] Mass of atoms (u) as reported by Wang *et al.* (Wang et al., 2012).
[b] Isotopic abundance in natural materials as reported by Wieser *et al.* (Wieser et al., 2013).
[c] Isotopic abundance reported by the manufacturer of [$^{13}C$]glucose and [$^{15}N$]$NH_4Cl$ (Cambridge Isotope Laboratories).

**Isotopologue abundance.** The first step in obtaining simulations of high resolution mass spectra is the calculation of the abundance of all possible isotopomers, *i.e.* molecules defined by the presence of a specific isotope at each position. Table 1 provides the mass and abundance of the C, H, N, and O isotopes that were used for these calculations. There is a total of nine isotopes: two for carbon, hydrogen, and nitrogen and three for oxygen. The elemental composition of the $[M+H]^+$ pseudo molecular ion is $C_{34}H_{58}N_6O_{20}{\bullet}H^+$ for the reduced disaccharide. According to the elemental composition of the mono-charged disaccharide-tripeptide (*i.e.* containing 59 hydrogens), there are 2.21 x $10^{39}$ isotopomers ($2^{34} * 2^{59} * 2^{6} * 3^{20}$) but 3,395,700 (35 * 60 * 7 * 231) distinct isotopologues (*i.e.* isotopomers with the same mass) since isotopomers with the same isotopic composition have the same mass irrespective of the position of the isotopes in the molecules. For the disaccharide-tripeptide, the isotopic composition of the isotopologues is defined by five numbers ($n_1$ to $n_5$) according to the number of $^{12}$C ($n_1$) and $^{13}$C (34-$n_1$) nuclei for carbon, the number of $^{1}$H ($n_2$) and $^{2}$H (59-$n_2$) nuclei for hydrogen, the number of $^{14}$N ($n_3$) and $^{15}$N (6-$n_3$) nuclei for nitrogen, and the number of $^{16}$O ($n_4$), $^{17}$O ($n_5$), and $^{18}$O (20-$n_4$-$n_5$) nuclei for oxygen. The abundance of the isotopologues with a defined isotopic composition depends upon the abundance of the isotopes listed in Table 1 and is the product of four relative abundances ($RA_C$ * $RA_H$ * $RA_N$ * $RA_O$) independently calculated for the four elements [For carbon: $RA_C = 0.9893^{n1} * 0.0107^{(35\text{-}n1)}$ * the number of combinations of $n_1$ objects among 35 ($C_{35,n1}$); For hydrogen: $RA_H = 0.999885^{n2} * 0.000115^{(59\text{-}n2)} * (C_{59,n2})$; For nitrogen: $RA_N = 0.99636^{n3} * 0.00364^{(6\text{-}n3)} * (C_{6,n3})$; For oxygen: $RA_O = 0.99757^{n4} * 0.00038^{n5} * 0.00205^{(20\text{-}n4\text{-}n5)} * (C_{20,n4,n5})$]. Figure 1A provides the general formula for molecules composed of any combination of elements and isotopes. Fig. 1B provides, as an example, the application of this formula to the calculation of the abundance of the disaccharide-tripeptide isotopologue containing light isotopes except for two $^{13}$C and one $^{15}$N atoms. The formula is valid under the assumption that the incorporation of an isotope at a specific position of a muropeptide solely depends upon the average abundance of this isotope in the culture medium. This requires, but is not guaranteed by, a uniform distribution of $^{12}$C and $^{13}$C nuclei at the six positions of labeled glucose used as the sole source of carbon in the labeled medium. Minor deviations from this assumption may occur since the rate of enzymatic reactions is known to be marginally affected by the isotopic composition of substrates and products.

Figure 1C shows the mass and abundance of the 25 most abundant isotopologues calculated for the unlabeled medium using the natural abundance of the isotopes (Table 1). The cumulative abundance of these isotopologues was 99.9008% indicating that the remaining isotopologues have minor contributions to the intensity of the peaks in the actual mass spectra. In practice, computing the relative abundance of the full complement of the 3,395,700 distinct isotopologues is time-consuming but the algorithm described by Lacki *et al.* (Lacki et al., 2017) provides an efficient means to limit calculations to the most abundant isotopologues by setting a limit for the cumulative abundance (a limit of 99.99% was used in the current study). Note that the most abundant isotopologue (64.2053%) exclusively contained the light isotopes of the four elements ($^{12}$C, $^{1}$H, $^{14}$N, and $^{16}$O). The second and third most abundant isotopologues (23.6105% and 4.2135%) contained light isotopes of the four elements except for one and two $^{13}$C carbon atoms, respectively. This was accounted for by the relatively high natural abundance of the $^{13}$C isotope (1.07%) and the important contribution of carbon to the total number of atoms present in the disaccharide-tripeptide (34 out of 119 atoms). In comparison, nitrogen only contributed to 6 of the 119 atoms accounting for the modest contribution of isotopologues containing a single $^{15}$N nucleus to the isotopologue pool (1.4074%) despite the relatively high natural abundance of $^{15}$N (0.364%).

Figure 1D shows the mass and abundance of the 25 most abundant isotopologues calculated for the labeled medium (Table 1), which contains the hydrogen and oxygen isotopes in natural abundance and predominantly the $^{13}$C (99%) and $^{15}$N (99%) isotopes of carbon and nitrogen according to the data sheets from the manufacturer of [$^{13}$C]glucose and [$^{15}$N]$NH_4Cl$, respectively. The cumulative abundance of these isotopologues was 99.8327% indicating that they should be the major contributors to isotopic clusters. The most abundant isotopologue only contained $^{13}$C and $^{15}$N nuclei. The next most abundant isotopologues contained a single unlabeled nucleus, $^{12}$C (21.7356%) or $^{14}$N (3.8357%), reflecting the similar relative abundance of these isotopes (1%) and the elemental composition of the disaccharide tripeptide (34 carbons *versus* 6 nitrogens), respectively.

**Shaping the isotopic cluster peaks using the calculated abundance of isotopologues**. The calculated isotopologue abundances cannot be easily compared to experimental isotopic cluster data because the former only provide relative abundances for each discrete (centroid) *m*/*z* value of the isotopic cluster peaks whereas the latter provide fully profiled cluster peaks, the shape of which is defined by the resolution of the mass spectrometer, the treatment of the signal, and the convolution of signals from isotopologues with almost identical masses. The effect of signal convolution is illustrated by the distribution of the masses of isotopologues in representative cluster peaks, calculated as described in Fig. 1 but sorted, this time, by increasing mass (Fig. 2A and 2B). This analysis shows that the masses expected to generate the peaks of the isotopic clusters differ by approximately, but not exactly, one unit. This is due to the fact that the gain of a neutron by different nuclei does not lead to exactly the same mass increment as the gain in energy in the resulting nuclei is not exactly the same. For example, the gain of a neutron by the $^{12}$C and $^{14}$N nuclei results in mass increments of 1.00335 and 0.99703, respectively. Thus, the distribution of the relative abundances of the isotopologues can only be visualized as centroids (Figure 2C). To enable the comparison of the calculated isotopic cluster peaks with experimental mass spectra, a Gaussian peak was modeled for each isotopic composition according to equations 1 and 2 (Fig. 2D). Each peak was subsequently scaled by the relative abundance of the isotopologues and overlaid (Fig. 2E). In order to obtain simulated mass spectra that could effectively be compared to experimental data, the modeled peaks were binned with a bin size of 0.002 *m/z* corresponding to the bin size deduced from the experimental mass spectra (Fig. 2F).

**Estimation of the relative increase in the amount of peptidoglycan and muropeptides upon bacterial growth**. Since the amount of peptidoglycan per cell remains constant during exponential growth, the amount of peptidoglycan $f_{PG}(t)$ was considered to increase with time according to equation $f_{PG}(t) = PG_0 * 2^{t/g}$ (equation 3) in which $PG_0$ is the amount of peptidoglycan at t=0 (normalized to 1), *g* is the generation time, and t=0 corresponds to the medium switch time. Since the relative amount of the muropeptides did not vary (above) the normalized molar amount of muropeptide *m*, $f_m(t)$, was considered to increase with time according to equation 4, $f_m(t) = m_0 * 2^{t/g}$ , in which $m_0$ is the relative molar amount of muropeptide *m* at t=0. This approach enables monitoring the increase in the amount of muropeptides during exponential growth relative to the amount present at t=0.

**Estimates of the molar abundance of muropeptide isotopologues.** The relative molar abundance of muropeptide isotopologues detected in the same mass spectrum was deduced from the relative maximal current intensity ($A_{mi}$) for *m/z* values corresponding to the $[M+H]^+$ and $[M+H]^{2+}$ ions of isotopologues for the monomers and dimers, respectively. These estimates are robust since that they are based on the detection of chemically identical molecular ions in the same injection, which is standard practice in quantitative mass spectrometry. The increase in the normalized amount of isotopologue *i* of muropeptide *m*, $f_{mi}(t)$, was deduced from the normalized molar amount of muropeptide *m* [equation 4, $f_m(t) = m_0 * 2^{t/g}$; above] and from $A_{mi}$, the relative current intensity of the isotopologue i (above) according to equation 5 [$f_{mi}(t) = A_{mi} * m_0 * 2^{t/g}$].

**A**

$$P = \prod_{e \in E} X\left((n_e(i_1) = k_{i_1}) \cap (n_e(i_2) = k_{i_2}) \cap \ldots \cap (n_e(i_m) = k_{i_m})\right)$$

$$X\left((n_e(i_1) = k_{i_1}) \cap (n_e(i_2) = k_{i_2}) \cap \ldots \cap (n_e(i_m) = k_{i_m})\right) = \frac{t_e!}{k_{i_1}!\, k_{i_2}! \ldots k_{i_m}!} * p_e(i_1)^{k_{i_1}} * \ldots * p_e(i_m)^{k_{i_m}}$$

| | |
|---|---|
| $P$: | Relative abundance of isotopologue |
| $E$: | Set of elements |
| $X$: | Joint probability of finding defined number of atoms of each isotope of element $e$ |
| $n_e(i_m)$: | Number of atoms of isotope $m$ of element $e$ |
| $p_e(i_m)$: | Relative abundance of isotope $m$ within element $e$ |
| $t_e$: | Total number of atoms of element $e$ (=$\sum_1^m n\,(i_m)$) |

**B**

**$C_{34}H_{58}N_6O_{20}$•$H^+$ All $^{12}C$ $^{1}H$ $^{14}N$ $^{16}O$, except 2 $^{13}C$ and 1 $^{15}N$**

| **Isotopic composition** | **Contribution of each element** |
|---|---|
| $^{12}C$ n=32 $^{13}C$ n=2 | $0.9893^{32}$ * $0.0107^{2}$ * 34! / (32! * 2!)<br>0.708755 * 0.000114 * 561 = **0.045523** |
| $^{1}H$ n=59 | $0.999885^{59}$ = **0.993238** |
| $^{14}N$ n=5 $^{15}N$ n=1 | $0.99636^{5}$ * $0.00364^{1}$ * 6! / (5! * 1!)<br>0.98193201 * 0.00364 * 6 = **0.021445** |
| $^{16}O$ n=20 | $0.99757^{20}$ = **0.952506** |

**Relative abundance of the isotopologue**

0.045523 * 0.993238 * 0.021445 * 0.952506 = **0.000924 or 0.0924%**

**C**

| **Isotopic composition**<br>**All $^{12}C$ $^{1}H$ $^{14}N$ $^{16}O$ except** | **Exact mass** | **Abundance %** |
|---|---|---|
| None | 871.3779 | 64.2053 |
| 1 $^{13}C$ | 872.3812 | 23.6105 |
| 2 $^{13}C$ | 873.3846 | 4.2135 |
| 1 $^{18}O$ | 873.3821 | 2.6388 |
| 1 $^{15}N$ | 872.3749 | 1.4074 |
| 1 $^{13}C$ 1 $^{18}O$ | 874.3855 | 0.9704 |
| 1 $^{13}C$ 1 $^{15}N$ | 873.3783 | 0.5175 |
| 1 $^{17}O$ | 872.3821 | 0.4891 |
| 3 $^{13}C$ | 874.3879 | 0.4861 |
| 1 $^{2}H$ | 872.3841 | 0.4357 |
| 1 $^{13}C$ 1 $^{17}O$ | 873.3854 | 0.1799 |
| 2 $^{13}C$ 1 $^{18}O$ | 875.3888 | 0.1732 |
| 1 $^{2}H$ 1 $^{13}C$ | 873.3875 | 0.1602 |
| 2 $^{13}C$ 1 $^{15}N$ | 874.3816 | 0.0924 ⟵ |
| 1 $^{15}N$ 1 $^{18}O$ | 874.3791 | 0.0578 |
| 2 $^{18}O$ | 875.3864 | 0.0515 |
| 4 $^{13}C$ | 875.3913 | 0.0407 |
| 2 $^{13}C$ 1 $^{17}O$ | 874.3888 | 0.0321 |
| 1 $^{2}H$ 2 $^{13}C$ | 874.3909 | 0.0286 |
| 1 $^{13}C$ 1 $^{15}N$ 1 $^{18}O$ | 875.3825 | 0.0213 |
| 3 $^{13}C$ 1 $^{18}O$ | 876.3922 | 0.0200 |
| 1 $^{17}O$ 1 $^{18}O$ | 874.3863 | 0.0191 |
| 1 $^{13}C$ 2 $^{18}O$ | 876.3897 | 0.0189 |
| 1 $^{2}H$ 1 $^{18}O$ | 874.3884 | 0.0179 |
| 2 $^{15}N$ | 873.3719 | 0.0129 |

Cumulative frequency 99.9008 %

**D**

| **Isotopic composition**<br>**All $^{13}C$ $^{1}H$ $^{15}N$ $^{16}O$ except** | **Exact mass** | **Abundance %** |
|---|---|---|
| None | 911.4741 | 63.2890 |
| 1 $^{12}C$ | 910.4708 | 21.7356 |
| 1 $^{14}N$ | 910.4771 | 3.8357 |
| 2 $^{12}C$ | 909.4674 | 3.6226 |
| 1 $^{18}O$ | 913.4784 | 2.6012 |
| 1 $^{12}C$ 1 $^{14}N$ | 909.4737 | 1.3173 |
| 1 $^{12}C$ 1 $^{18}O$ | 912.4750 | 0.8933 |
| 1 $^{17}O$ | 912.4784 | 0.4822 |
| 1 $^{2}H$ | 912.4804 | 0.4295 |
| 3 $^{12}C$ | 908.4641 | 0.3903 |
| 2 $^{12}C$ 1 $^{14}N$ | 908.4704 | 0.2196 |
| 1 $^{12}C$ 1 $^{17}O$ | 911.4750 | 0.1656 |
| 1 $^{14}N$ 1 $^{18}O$ | 912.4813 | 0.1576 |
| 2 $^{12}C$ 1 $^{18}O$ | 911.4717 | 0.1489 |
| 1 $^{2}H$ 1 $^{12}C$ | 911.4771 | 0.1475 |
| 2 $^{14}N$ | 909.4801 | 0.0969 |
| 1 $^{12}C$ 1 $^{14}N$ 1 $^{18}O$ | 911.4780 | 0.0541 |
| 2 $^{18}O$ | 915.4826 | 0.0508 |
| 1 $^{12}C$ 2 $^{14}N$ | 908.4767 | 0.0333 |
| 4 $^{12}C$ | 907.4607 | 0.0306 |
| 1 $^{14}N$ 1 $^{17}O$ | 911.4813 | 0.0292 |
| 2 $^{12}C$ 1 $^{17}O$ | 910.4716 | 0.0276 |
| 1 $^{2}H$ 1 $^{14}N$ | 911.4834 | 0.0260 |
| 1 $^{2}H$ 2 $^{12}C$ | 910.4737 | 0.0246 |
| 3 $^{12}C$ 1 $^{14}N$ | 907.4670 | 0.0237 |

Cumulative frequency 99.8327%

**Figure 1 of Supplementary Material and Methods.**

**Calculation of the relative abundance of isotopologues.** (**A**) General formula for molecules composed of any combination of elements and isotopes. (**B**) Example of the calculation of the abundance of a mono-charged ($[M+H]^+$) GlcNAc-MurNAc-tripeptide molecular ion ($C_{34}H_{58}N_6O_{20}$•$H^+$;) exclusively containing light isotopes ($^{12}C$, $^{1}H$, $^{14}N$, $^{16}O$) except for two $^{13}C$ and one $^{15}N$ heavy isotopes. (**C** and **D**) Mass and relative abundance of the 25 most abundant GlcNAc-MurNAc-tripeptide isotopologues ($C_{34}H_{58}N_6O_{20}$•$H^+$), calculated for the unlabeled and labeled media, respectively, and sorted by decreasing abundance. The arrow points to the example of calculation detailed in panel B.

**Contribution of isotopologues to cluster peaks** A

| All $^{12}C$ $^{1}H$ $^{14}N$ $^{16}O$ | Exact mass | Abundance % |
|---|---|---|
| **Peak A (872.3809; 13.9922)** | ← | |
| 1 $^{2}H$ | 872.3841 | 0.4357 |
| 1 $^{17}O$ | 872.3821 | 0.4891 |
| 1 $^{13}C$ | 872.3812 | 23.6105 |
| 1 $^{15}N$ | 872.3749 | 1.4074 |
| **Peak B (873.3834; 4.1290)** | ← | |
| 1 $^{2}H$ 1 $^{13}C$ | 873.3875 | 0.1602 |
| 1 $^{13}C$ 1 $^{17}O$ | 873.3854 | 0.1799 |
| 2 $^{13}C$ | 873.3846 | 4.2135 |
| 1 $^{18}O$ | 873.3821 | 2.6388 |
| 1 $^{13}C$ 1 $^{15}N$ | 873.3783 | 0.5175 |
| **Peak C (874.3859; 0.9102)** | | |
| 1 $^{2}H$ 2 $^{13}C$ | 874.3909 | 0.0286 |
| 2 $^{13}C$ 1 $^{17}O$ | 874.3888 | 0.0321 |
| 3 $^{13}C$ | 874.3879 | 0.4861 |
| 1 $^{13}C$ 1 $^{18}O$ | 874.3855 | 0.9704 |
| 2 $^{13}C$ 1 $^{15}N$ | 874.3816 | 0.0924 |
| 1 $^{15}N$ 1 $^{18}O$ | 874.3791 | 0.0578 |
| **Peak D (875.3883; 0.1682)** | | |
| 4 $^{13}C$ | 875.3913 | 0.0407 |
| 2 $^{13}C$ 1 $^{18}O$ | 875.3888 | 0.1732 |
| 2 $^{18}O$ | 875.3864 | 0.0515 |
| 1 $^{13}C$ 1 $^{15}N$ 1 $^{18}O$ | 875.3825 | 0.0213 |

**Contribution of isotopologues to cluster peaks** B

| All $^{13}C$ $^{1}H$ $^{15}N$ $^{16}O$ except | Exact mass | Abundance % |
|---|---|---|
| **Peak A (908.4669; 0.3316)** | | |
| 1 $^{12}C$ 2 $^{14}N$ | 908.4767 | 0.0333 |
| 2 $^{12}C$ 1 $^{14}N$ | 908.4704 | 0.2196 |
| 3 $^{12}C$ | 908.4641 | 0.3903 |
| **Peak B (909.4693; 2.6356)** | | |
| 2 $^{14}N$ | 909.4801 | 0.0969 |
| 1 $^{12}C$ 1 $^{14}N$ | 909.4737 | 1.3173 |
| 2 $^{12}C$ | 909.4674 | 3.6226 |
| **Peak C (910.4717; 13.7225)** | | |
| 1 $^{14}N$ | 910.4771 | 3.8357 |
| 1 $^{12}C$ | 910.4708 | 21.7356 |
| **Peak D (911.4742; 35.0000)** | | |
| 1 $^{12}C$ 1 $^{14}N$ 1 $^{18}O$ | 911.4780 | 0.0541 |
| 1 $^{2}H$ 1 $^{12}C$ | 911.4771 | 0.1475 |
| 1 $^{12}C$ 1 $^{17}O$ | 911.4750 | 0.1656 |
| *All $^{13}C$ $^{1}H$ $^{15}N$ $^{16}O$* | 911.4741 | 63.2890 |
| 2 $^{12}C$ 1 $^{18}O$ | 911.4717 | 0.1489 |
| **Peak E (912.4775; 1.0413)** | | |
| 1 $^{14}N$ 1 $^{18}O$ | 912.4813 | 0.1576 |
| 1 $^{2}H$ | 912.4804 | 0.4295 |
| 1 $^{17}O$ | 912.4784 | 0.4822 |
| 1 $^{12}C$ 1 $^{18}O$ | 912.4750 | 0.8933 |

D Equation 1 $\mu = \frac{m_{iso} + z * m_H}{z}$ Equation 2 $\sigma = \frac{FWHM}{2.355}$

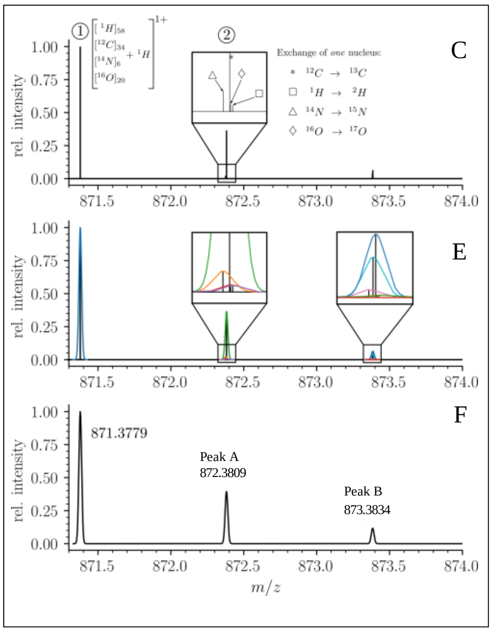


**Figure 2 of Supplementary Material and Methods.**

**Shaping mass spectra.** (**A**) and (**B**) Mass and abundance of GlcNAc-MurNAc-tripeptide isotopologues ($C_{34}H_{58}N_6O_{20}$•$H^+$) calculated for the unlabeled and labeled media, respectively. Data are sorted by increasing mass to highlight isotopologues that fall within very narrow mass intervals. The mass and the relative abundance are shown for isotopologues with major quantitative contributions to representative cluster peaks taken as examples. Arrows point to peaks analyzed in details in panels C, E, and F. (**C**) Abundance of isotopologues (for the unlabeled medium) visualized as a centroid spectrum. The magnification shows individual peaks within the cluster. Peak A consists of four isotopologues exclusively containing light isotopes except for one $^{2}H$, $^{13}C$, $^{15}N$, or $^{17}O$ nucleus. Peak B consists of seven isotopologues exclusively containing light isotopes except for one $^{18}O$ nucleus or combinations of two nuclei among $^{2}H$, $^{13}C$, $^{15}N$, and $^{17}O$. (**D**) Equations 1 and 2 used to model Gaussian peaks. $m_H$ is the mass of a proton ($H^+$), $z$ is the net charge of the analyte, and FWHM is the full width at half maximum. The value of FWHM used in this study (40,000) was chosen to match the resolving power of the mass spectrometer deduced from experimental spectra. (**E**) Modeling of Gaussian peaks for each cluster peaks according to equations 1 and 2. (**F**) Simulated mass spectrum. The mass at the apex of the peaks, which was used for comparison of simulated and experimental mass spectra, is indicated. These values are reported in panels A and B for each cluster along with the maximum intensity of the simulated peaks (arbitrary unit, shown in parenthesis).
