## Supplementary Tables and Figures for "Heavy isotope labeling and mass spectrometry reveal unexpected remodeling of bacterial cell wall expansion in response to drugs"

**Table S1. Muropeptide composition of the peptidoglycan of strain BW25113 Δ6*ldt* and M1.5**

| **Strain** | **Muropetide (%)[a]** | | | | | | | |
|---|---|---|---|---|---|---|---|---|
| **β-lactam[b]** | **Tri** | **Tetra** | **Penta** | **Tri→Tri** | **Tri→Tetra** | **Tri→Penta** | **Tetra→Tri** | **Tetra→Tetra** |
| BW25113 Δ6*ldt* | | | | | | | | |
| None (9) [c] | ND | 74 ± 2 | ND | ND | ND | ND | ND | 26 ± 2 |
| AZT (5) [c] | ND | 77 ± 1 | ND | ND | ND | ND | ND | 23 ± 1 |
| MEL (5) [c] | ND | 70 ± 2 | ND | ND | ND | ND | ND | 30 ± 2 |
| M1.5 | | | | | | | | |
| None (12) [c] | 31 ± 3 | 36 ± 4 | ND | 7 ± 1 | 8 ± 1 | ND | 7 ± 1 | 12 ± 1 |
| AMP (10) [c] | 29 ± 2 | 21 ± 3 | 6 ± 2 | 13 ± 2 | 24 ± 1 | 6 ± 0.4 | 1 ± 0.3 | ND |

[a] Data are the mean ± standard deviation from independent experiments (see note c).

[b] β-lactam added to the growth medium. AZT, aztreonam at 12 µg/ml; MEL, mecillinam at 2.5 µg/ml; AMP, ampicillin at 16 µg/ml.

[c] The number of independent peptidoglycan analyses is indicated in parenthesis.

**Abbreviations:** ND, not detected; Tri, tripeptide monomer; Tetra, tetrapeptide monomer; Penta, pentapeptide monomer; Tri→Tri, Tri→Tetra, and Tri→Penta, 3→3 cross-linked dimers; Tetra→Tri and Tetra→Tetra, 4→3 cross-linked dimers. The inter-peptide cross-link direction is indicated using the donor→acceptor conventional notation.

**Table S2. Generation time for growth of BW25113 derivatives in M1 minimal medium**

| Strain | Generation time (min) | |
|---|---|---|
| β-lactam [a] | Experiment 1 | Experiment 2 |
| M1.5 | | |
| None | 90 ± 2 | 85 ± 3 |
| AMP | 163 ± 9 | 165 ± 13 |
| BW25113 Δ6*ldt* | | |
| None | 67 ± 1 | 66 ± 1 |
| AZT [b] | 64 ± 1 | 63 ± 1 |
| MEL [b] | 63 ± 1 | 65 ± 1 |

[a] β-lactam added to the growth medium. AZT, aztreonam at 12 µg/ml; MEL, mecillinam at 2.5 µg/ml; AMP, ampicillin at 16 µg/ml.

[b] For growth in the presence of aztreonam and mecillinam, the values were deduced from the variation in $OD_{600}$ during 90 min.

### SUPPLEMENTARY FIGURES

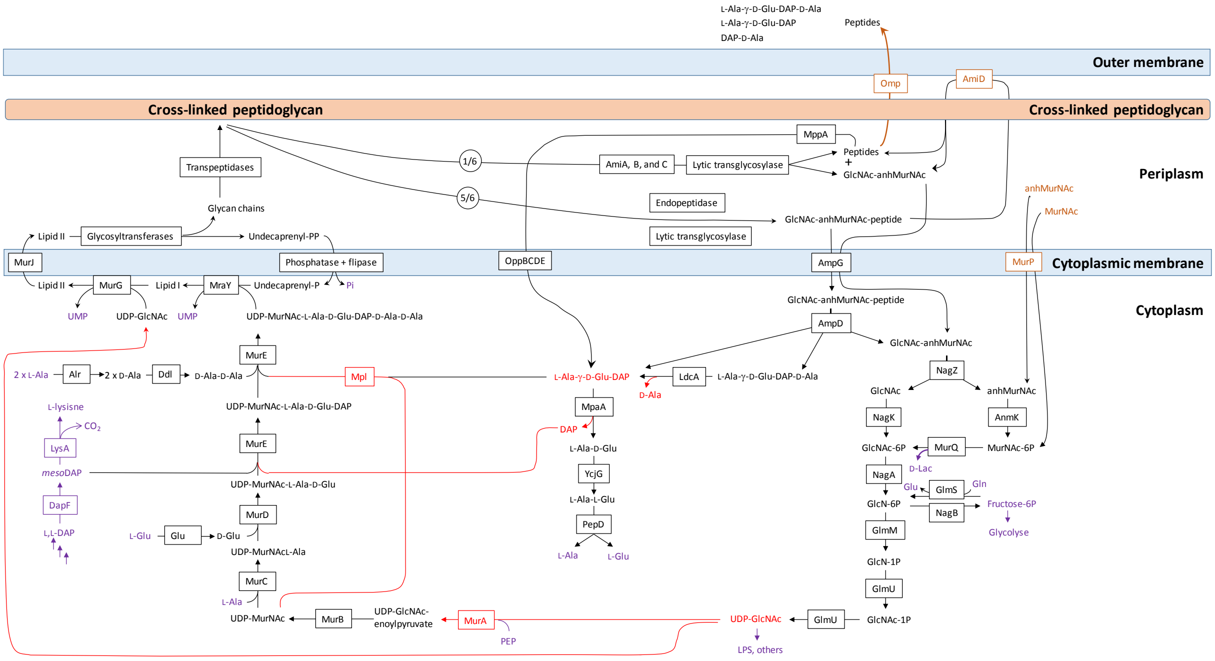


**Supplementary Fig. S1. Synthesis and recycling of peptidoglycan in *E. coli*.** Key intermediates connecting synthesis (on the left) to recycling (on the right) are figured in red. Metabolites common to peptidoglycan metabolism and to other metabolic pathways are figured in purple. Effective incorporation of DAP from the culture medium requires inactivation of the genes encoding DapF and LysA generating lysine auxotrophy. Peptidoglycan turnover refers to the release of peptidoglycan fragments (<600 Da) into the culture medium, which cross the outer membrane through porins.

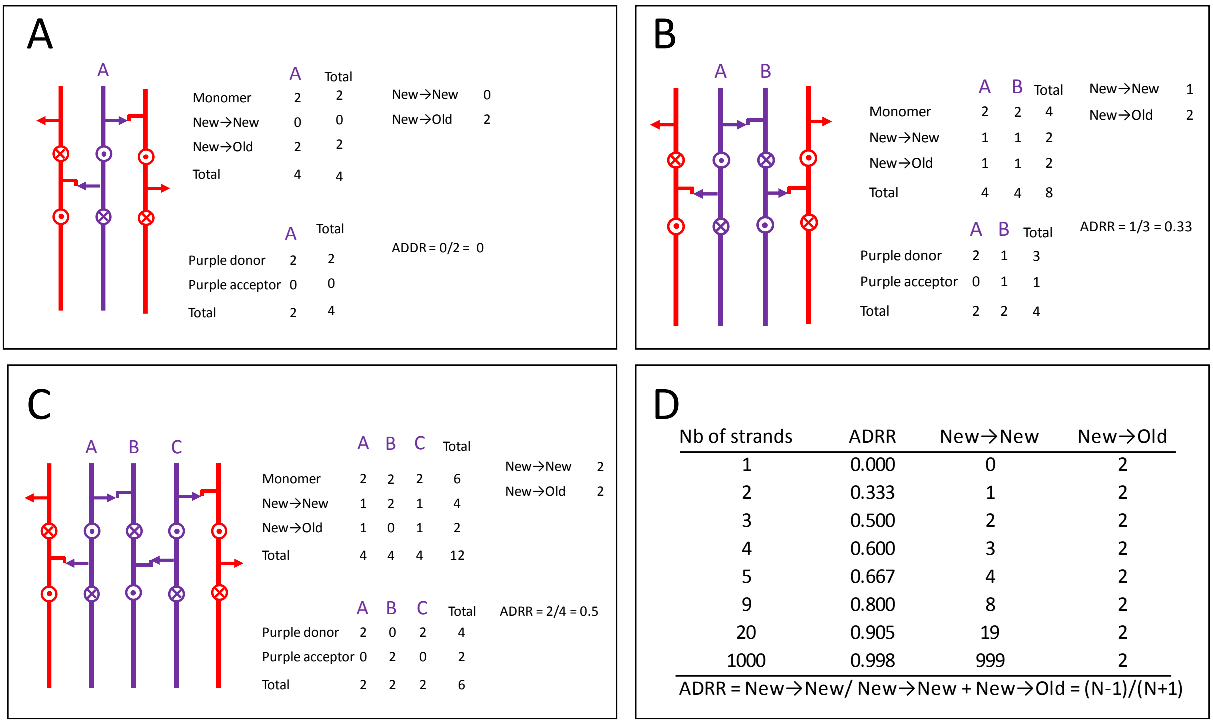


**Supplementary Fig. S2. Acceptor-to-donor radioactivity ratio (ADRR) as a function of the number of glycan strands inserted at the same time in the peptidoglycan layer.** (**A**) One strand at a time. (**B**) Two strands at a time. (**C**) Three stands at a time (applies to the three for one model). (**D**) Calculation for various number of strands. Existing and neo-synthesized strands are figures in red and in purple, respectively. Donor and acceptor stems are represented by an arrow and by an L, respectively. Uncross-linked peptide stems that protrude above and below the plan are indicated by circles with a dot or an X, respectively.

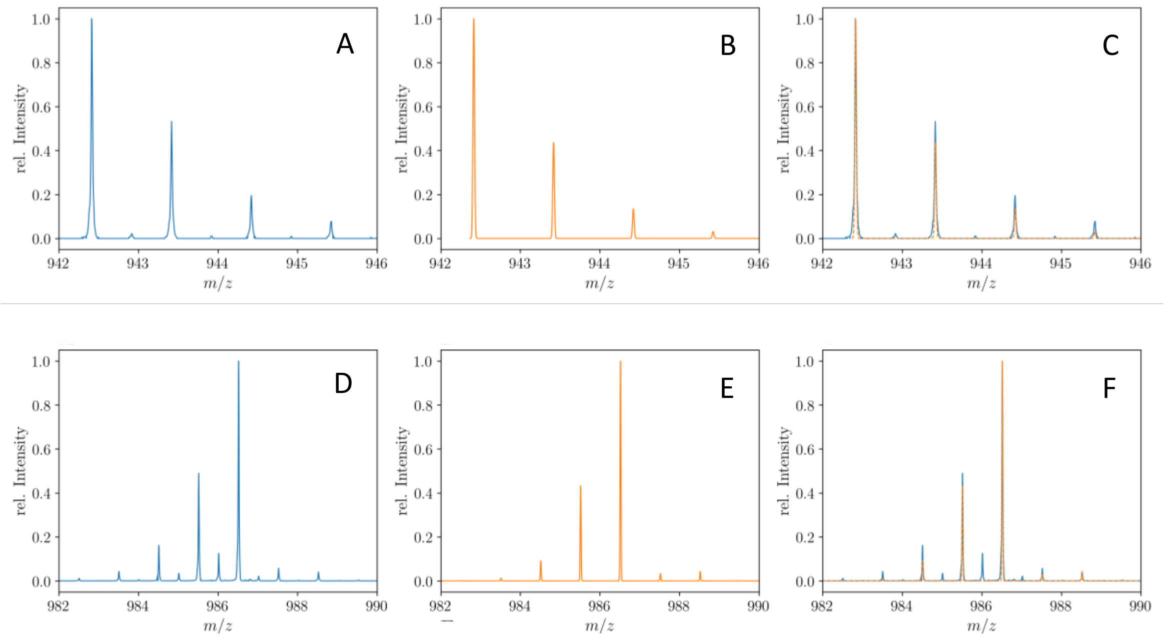


**Supplementary Fig. S3. Mass spectra of the unlabeled and fully labeled disaccharide-tetrapeptide monomers.** (**A**) Experimental mass spectrum of the mono-protonated ($[M+H]^{1+}$) disaccharide-tetrapeptide ($C_{37}H_{63}N_7O_{21}$•$H^+$) extracted from *E. coli* M1.5 grown in unlabeled M9 minimal medium. (**B**) Simulated spectrum obtained for the natural abundance of carbon and nitrogen isotopes. (**C**) Overlay of the spectra in A and B. (**D**) Observed mass spectrum of the GlcNAc-MurNAc-tetrapeptide purified from the peptidoglycan of *E. coli* M1.5 grown in the labeled M9 minimal medium. (**E**) Simulated mass spectrum obtained for 99% $^{13}C$ and $^{15}N$ labeling. (**F**) Overlay of spectra in D and E. The additional peaks of low intensity in the experimental spectra, in particular for the labeled disaccharide-tetrapeptide, correspond to the $[2M+2H]^{2+}$ ion.

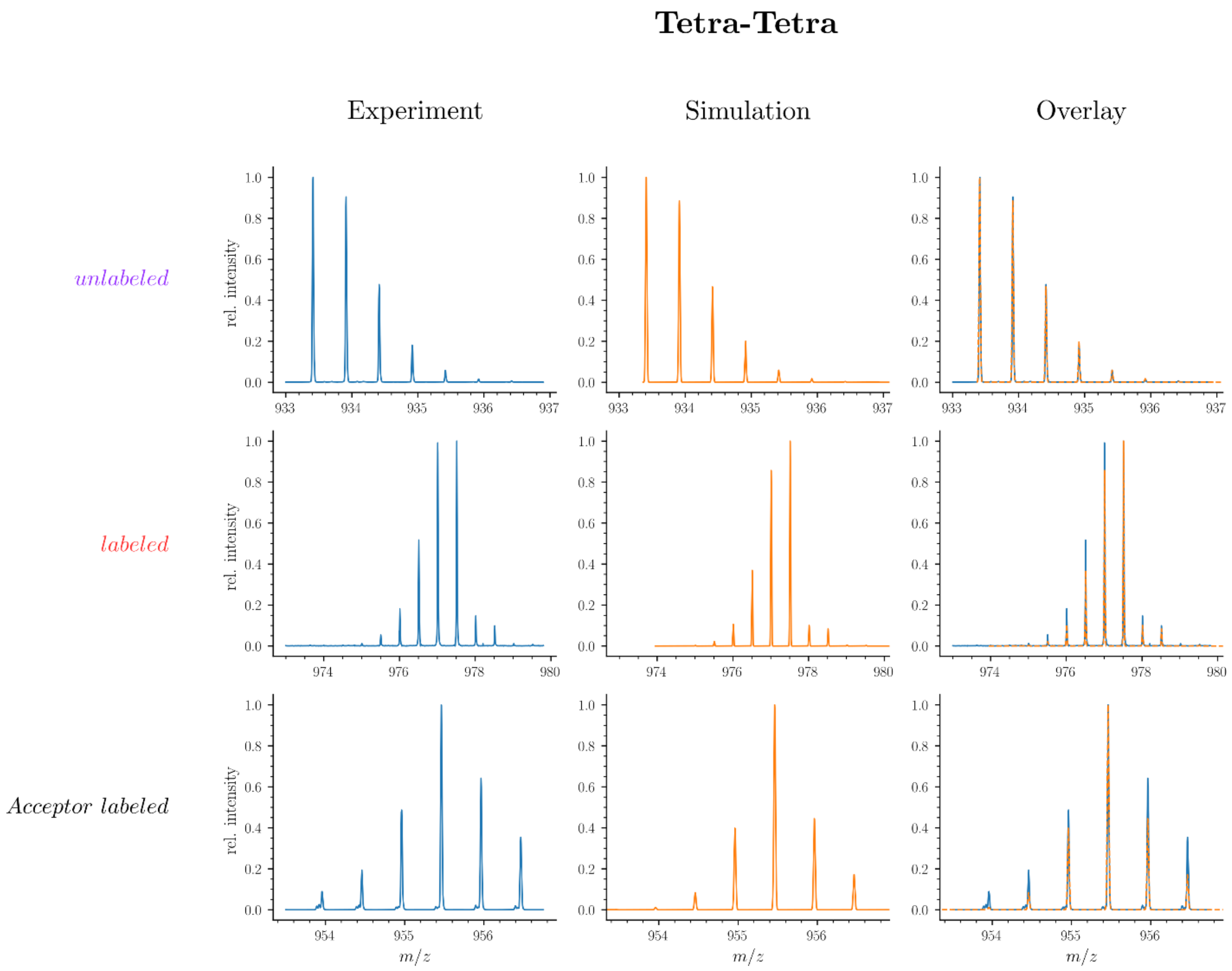


**Supplementary Fig. S4. Mass spectra of unlabeled and fully labeled Tetra→Tetra dimer.**

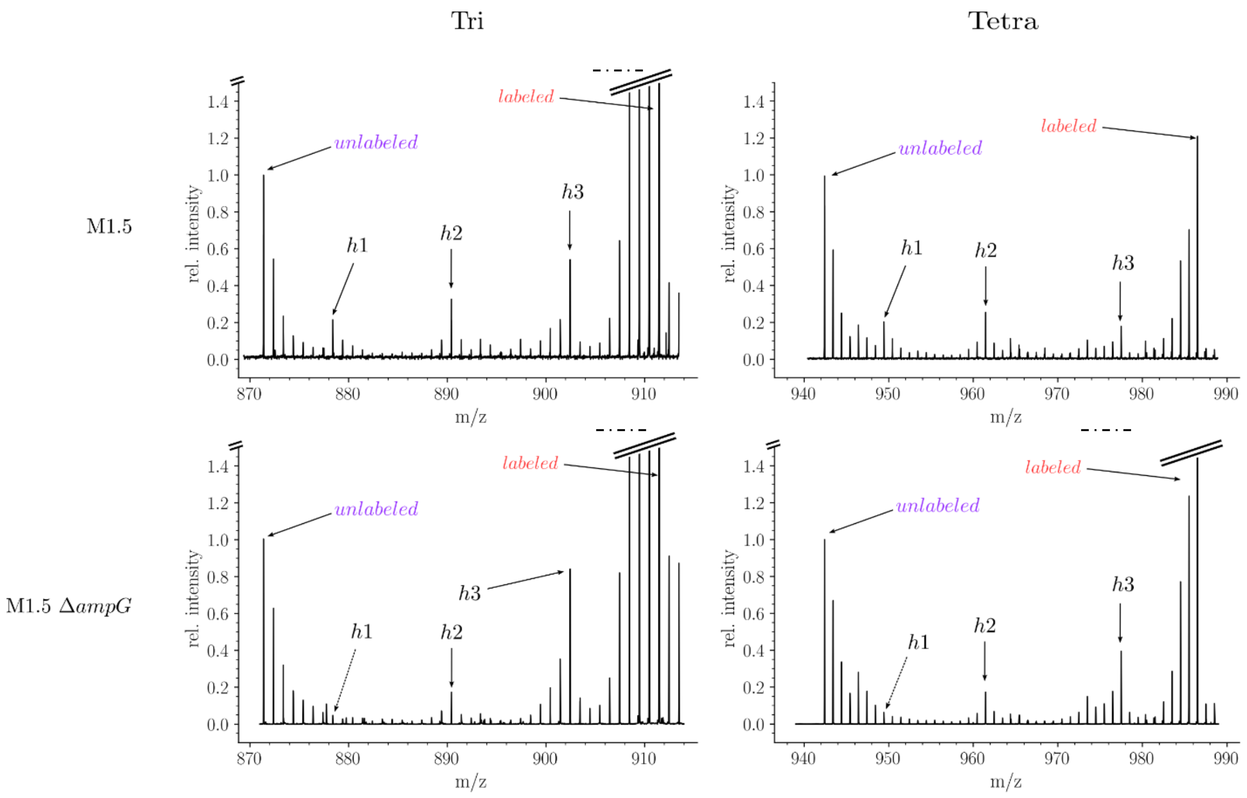


**Supplementary Fig. S5. Isotopic composition of monomers isolated from the peptidoglycan of *E. coli* M1.5 and a recycling-deficient derivative obtained by deletion of the *ampG* permease gene.**

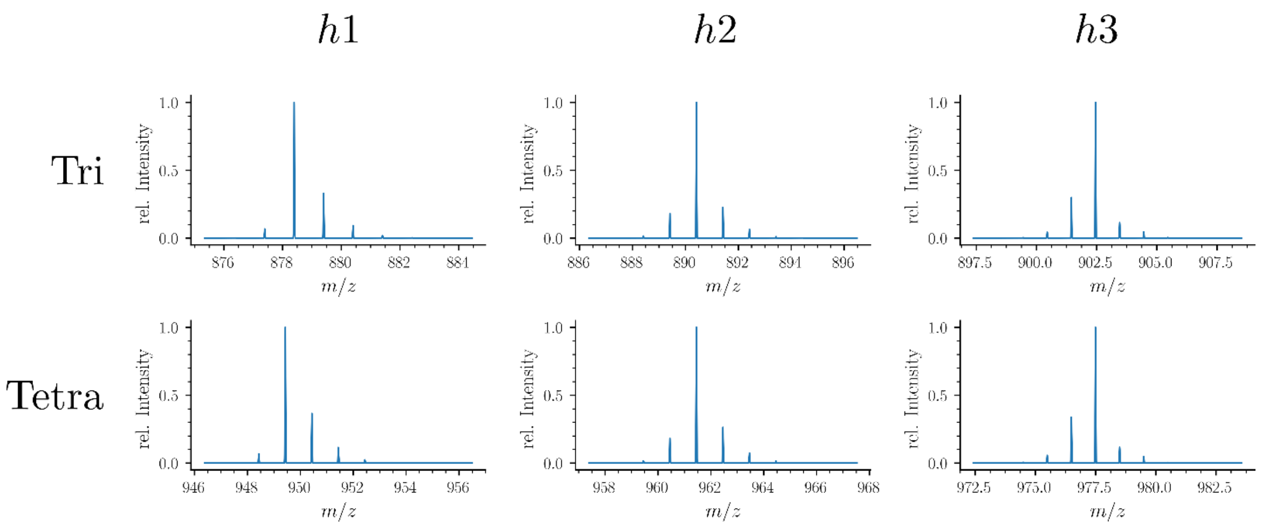


**Supplementary Fig. S6. Simulated spectra of hybrid monomers.**

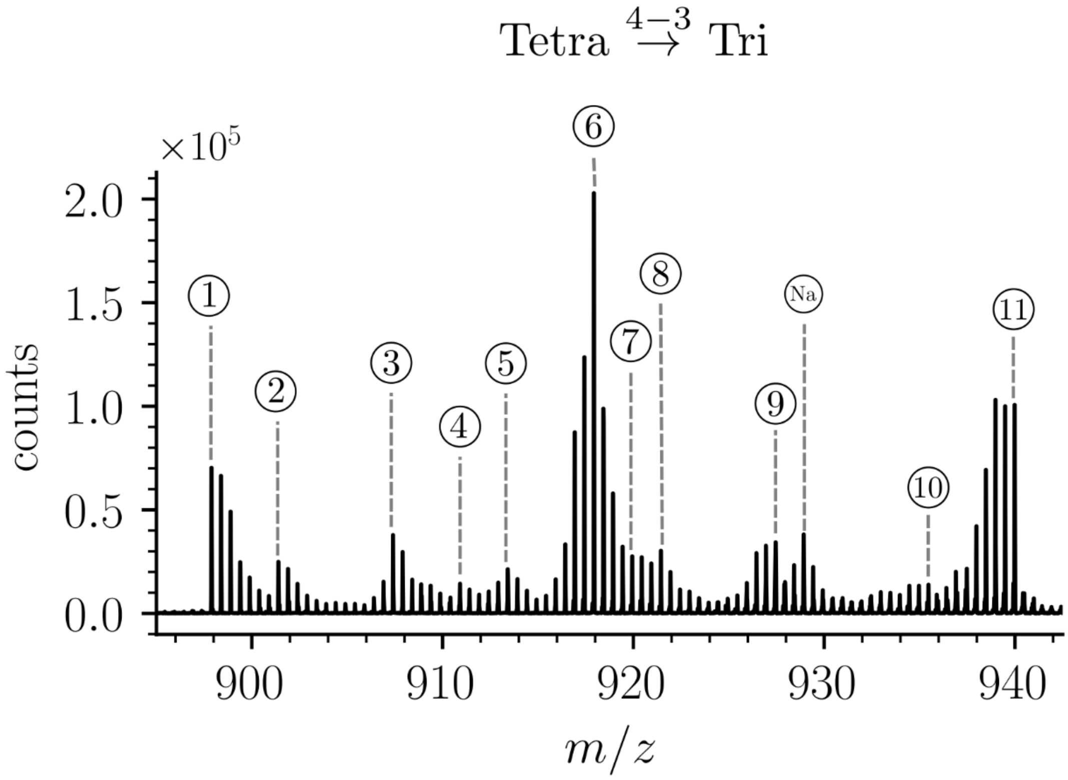


| | m/z | | | | Isotopologue | Origin (MS) | |
|---|---|---|---|---|---|---|---|
| | obs | calc | Ion | ppm | Donor→Acceptor | Donor→Acceptor | % |
| ① | 897.8935 | 897.8917 | $[M+2H]^{2+}$ | 2.1 | D→A | new→new | 11.0 |
| ② | 901.3974 | 901.4003 | $[M+2H]^{2+}$ | -3.2 | $D_{h1}$→A, D→$A_{h1}$ | new→new | 4.2 |
| ③ | 907.4109 | 907.4109 | $[M+2H]^{2+}$ | 0.1 | $D_{h2}$→A, D→$A_{h2}$ | new→new | 6.4 |
| ④ | 910.9203 | 910.9195 | $[M+2H]^{2+}$ | 0.9 | $D_{h1}$→$A_{h2}$, $D_{h2}$→$A_{h1}$<br>$D_{h2Hex}$→A, D→$A_{h2Hex}$ | new→new | 2.5 |
| ⑤ | 913.4263 | 913.4279 | $[M+2H]^{2+}$ | -1.8 | $D_{h3}$→A, D→$A_{h3}$ | new→new | 4.3 |
| ⑥ | 917.9383 | 917.9398 | $[M+2H]^{2+}$ | -1.7 | D→A | new→old | 35.8 |
| ⑦ | 919.9430 | 919.9434 | $[M+2H]^{2+}$ | -0.4 | $D_{hAla}$→A | new→old | 4.9 |
| ⑧ | 921.4472 | 921.4484 | $[M+2H]^{2+}$ | -1.3 | $D_{h1}$→A | new→old | 5.6 |
| ⑨ | 927.4563 | 927.4591 | $[M+2H]^{2+}$ | -3.0 | $D_{h2}$→A | new→old | 6.7 |
| (Na) | 928.9263 | 928.9308 | $[M+H+Na]^{2+}$ | 5.2 | D→A | new→new | NA |
| ⑩ | 935.4768 | 935.4796 | $[M+2H]^{2+}$ | -3.0 | $D_{h3}$→A | new→old | 2.4 |
| ⑪ | 939.9917 | 939.9915 | $[M+2H]^{2+}$ | 0.2 | D→A | old→old | 16.2 |

**Supplementary Fig. S7. Structure of Tetra→Tri isotopologues in the peptidoglycan of strain M1.4 grown in the absence of β-lactams.**

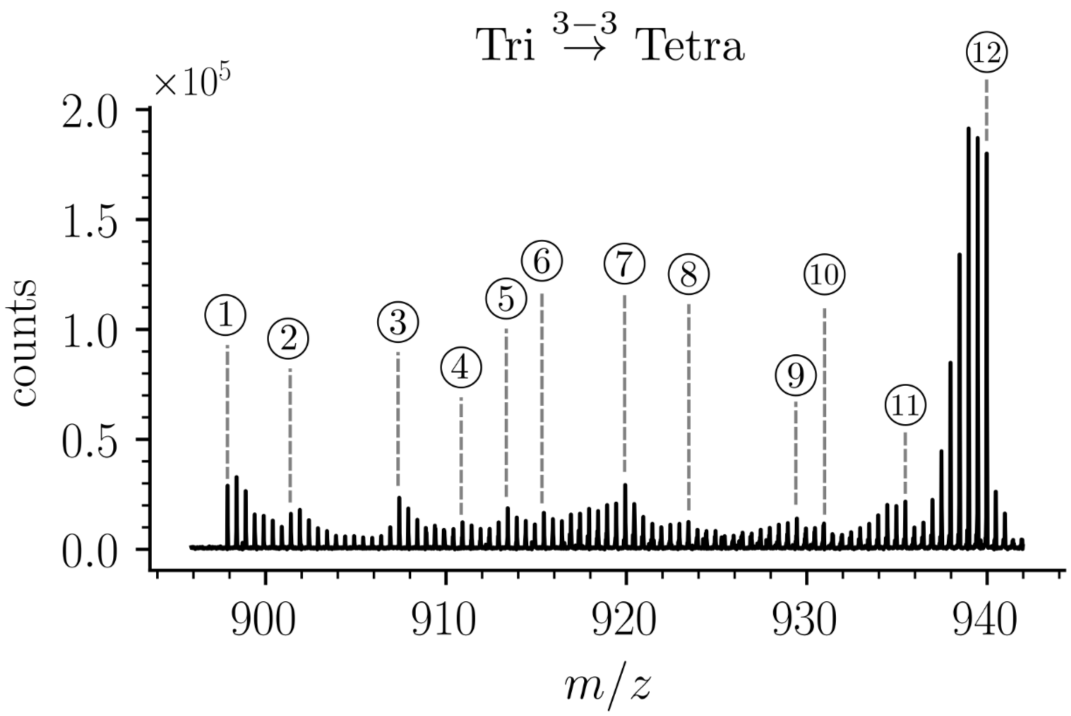


| | m/z | | | | Isotopologue | Origin (MS) | |
|---|---|---|---|---|---|---|---|
| | obs | calc | Ion | ppm | Donor→Acceptor | Donor→Acceptor | % |
| ① | 897.8935 | 897.8917 | $[M+2H]^{2+}$ | 2.5 | D→A | new→new | 8.2 |
| ② | 901.3974 | 901.4003 | $[M+2H]^{2+}$ | 0.4 | $D_{h1}$→A, D→$A_{h1}$ | new→new | 4.7 |
| ③ | 907.4109 | 907.4109 | $[M+2H]^{2+}$ | 2.1 | $D_{h2}$→A, D→$A_{h2}$ | new→new | 6.3 |
| ④ | 910.9203 | 910.9195 | $[M+2H]^{2+}$ | 0.4 | $D_{h1}$→$A_{h2}$, $D_{h2}$→$A_{h1}$ | new→new | 3.1 |
| | | | | | $D_{h2Hex}$→A, D→$A_{h2Hex}$ | | |
| ⑤ | 913.4263 | 913.4279 | $[M+2H]^{2+}$ | 1.1 | $D_{h3}$→A | new→new | 4.8 |
| ⑥ | 915.4293 | 915.4314 | $[M+2H]^{2+}$ | 1.7 | $D_{h3}$→$A_{hAla}$, D→$A_{h3}$ | new→new | 4.9 |
| ⑦ | 919.9430 | 919.9434 | $[M+2H]^{2+}$ | 3.1 | D→A | new→old | 7.0 |
| ⑧ | 923.4492 | 923.4520 | $[M+2H]^{2+}$ | 2.9 | $D_{h1}$→A | new→old | 3.2 |
| ⑨ | 929.4594 | 929.4626 | $[M+2H]^{2+}$ | 3.5 | $D_{h2}$→A | new→old | 3.6 |
| ⑩ | 930.9651 | 930.9676 | $[M+2H]^{2+}$ | 2.7 | $D_{h3}$→$A_{h3}$ | new→new | 2.8 |
| ⑪ | 935.4768 | 935.4796 | $[M+2H]^{2+}$ | 2.5 | $D_{h3}$→A | new→old | 5.6 |
| ⑫ | 939.9917 | 939.9915 | $[M+2H]^{2+}$ | 0.8 | D→A | old→old | 45.8 |

**Supplementary Fig. S8. Structure of Tri→Tetra isotopologues in the peptidoglycan of strain M1.4 grown in the absence of β-lactams.**

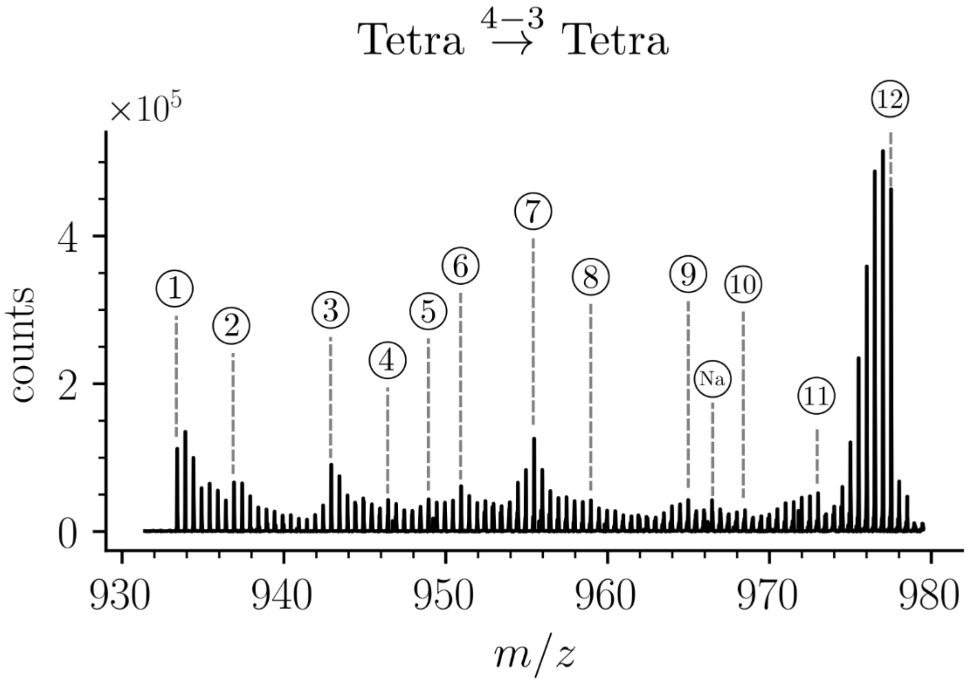


| | m/z | | | | Isotopologue | Origin (MS) | |
|---|---|---|---|---|---|---|---|
| | obs | calc | Ion | ppm | Donor→Acceptor | Donor→Acceptor | % |
| ① | 933.4097 | 933.4102 | $[M+2H]^{2+}$ | 0.6 | D→A | new→new | 8.2 |
| ② | 936.9174 | 936.9188 | $[M+2H]^{2+}$ | 1.5 | $D_{h1}\to A$, $D\to A_{h1}$ | new→new | 4.7 |
| ③ | 942.9261 | 942.9295 | $[M+2H]^{2+}$ | 3.6 | $D_{h2}\to A$, $D\to A_{h2}$ | new→new | 6.3 |
| ④ | 946.4369 | 946.4381 | $[M+2H]^{2+}$ | 1.2 | $D_{h1}\to A_{h2}$, $D_{h2}\to A_{h1}$<br>$D_{h2Hex}\to A$, $D\to A_{h2Hex}$ | new→new | 3.1 |
| ⑤ | 948.9410 | 948.9464 | $[M+2H]^{2+}$ | 5.8 | $D_{h3}\to A$, $D\to A_{h3}$ | new→new | 4.8 |
| ⑥ | 950.9477 | 950.9500 | $[M+2H]^{2+}$ | 2.5 | $D_{h3}\to A_{hAla}$, $D_{hAla}\to A_{h3}$ | new→new | 4.9 |
| ⑦ | 955.4594 | 955.4619 | $[M+2H]^{2+}$ | 2.6 | D→A | new→old | 7.0 |
| ⑧ | 958.9658 | 958.9705 | $[M+2H]^{2+}$ | 4.9 | $D_{h1}\to A$ | new→old | 3.2 |
| ⑨ | 964.9772 | 964.9812 | $[M+2H]^{2+}$ | 4.1 | $D_{h2}\to A$ | new→old | 3.6 |
| (Na) | 966.4509 | 966.4529 | $[M+H+Na]^{2+}$ | 1.8 | D→A | new→new | NA |
| ⑩ | 968.4868 | 968.4897 | $[M+2H]^{2+}$ | 3.0 | $D_{h3}\to A_{h3}$ | new→new | 2.8 |
| ⑪ | 973.0004 | 973.0017 | $[M+2H]^{2+}$ | 1.3 | $D_{h3}\to A$ | new→old | 5.6 |
| ⑫ | 977.5101 | 977.5136 | $[M+2H]^{2+}$ | 3.6 | D→A | old→old | 45.8 |

**Supplementary Fig. S9. Structure of Tetra→Tetra isotopologues in the peptidoglycan of strain M1.4 grown in the absence of β-lactams.**

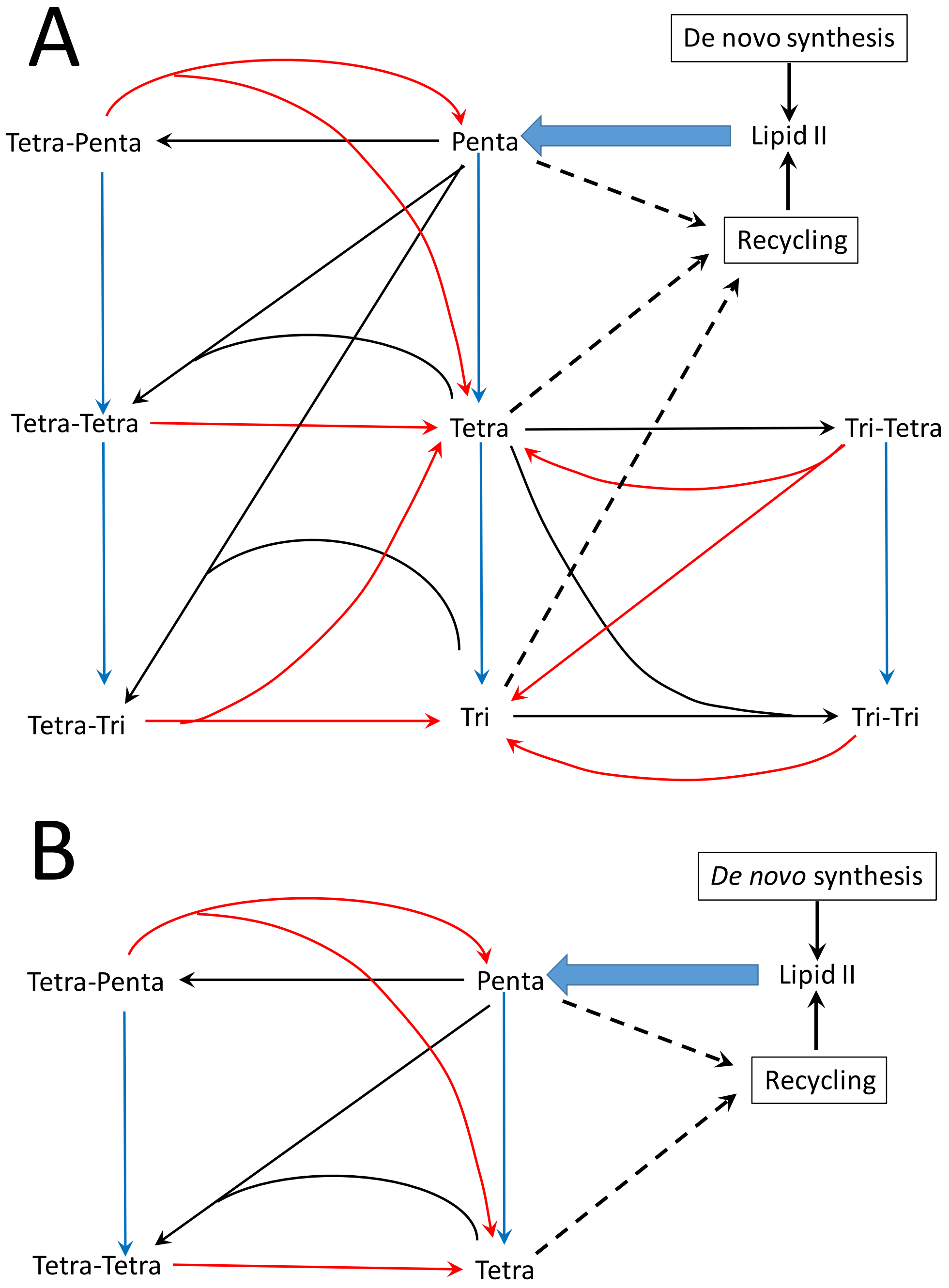


**Supplementary Fig. S10. Peptidoglycan metabolism in *E. coli* M1.5 (A) and Δ6*ldts* (B).** The reactions catalyzed by transpeptidases (formation of cross-links), endopeptidases (hydrolysis of cross-links), and carboxypeptidases (hydrolysis of C-terminal D-Ala$^4$ and D-Ala$^5$ are indicated by black, red, and blue arrows, respectively. PG is polymerized from a single precursor, Lipid II originates form *de novo* synthesis and from recycling. This lipid consists of the disaccharide-pentapeptide subunit linked to the lipid carrier (undecaprenyl) by a pyrophosphate bond.

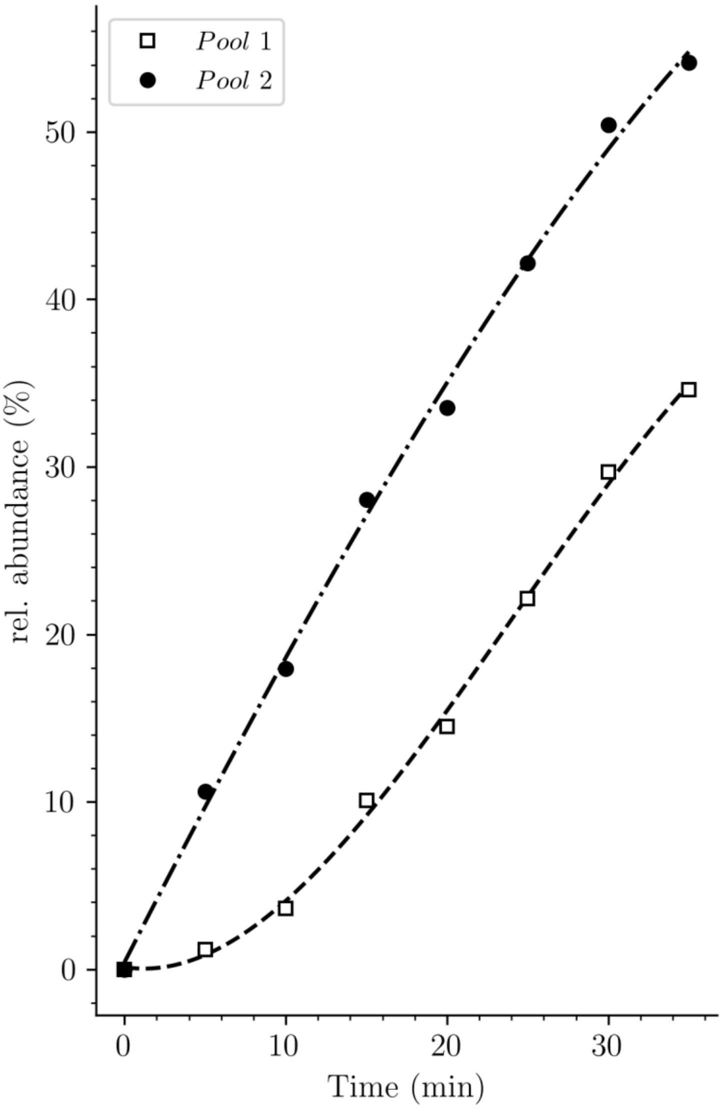


**Supplementary Fig. S11. Potential origin of the lag phase previously observed in experiments based on incorporation of radioactive DAP.** Kinetics of accumulation of neo-synthesized disaccharide-tetrapeptide isotopologues containing unlabeled DAP (pool 1, squares) was compared to kinetics of accumulation all neo-synthesized disaccharide-tetrapeptide isotopologues, including h2 and h3 hybrids (pool 2, solid circles) (Data from strain Δ6*ldt* grown in the absence of β-lactam).

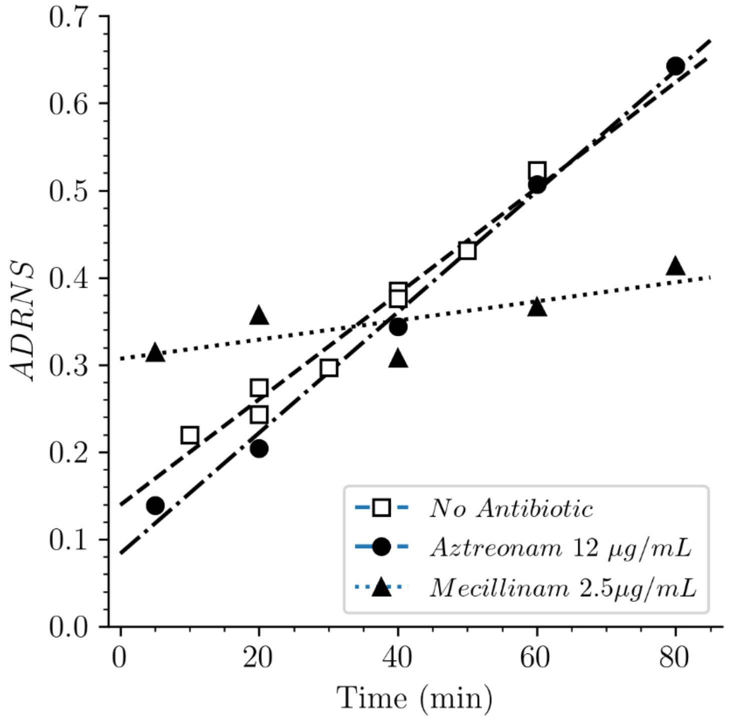


**Supplementary Fig. S12. Acceptor-to-donor ratio of neo-synthesized stems (ADRNS) of *E. coli* Δ6*ldt*.** Bacteria were grown in the presence of aztreonam, mecillinam, or in the absence of drug.

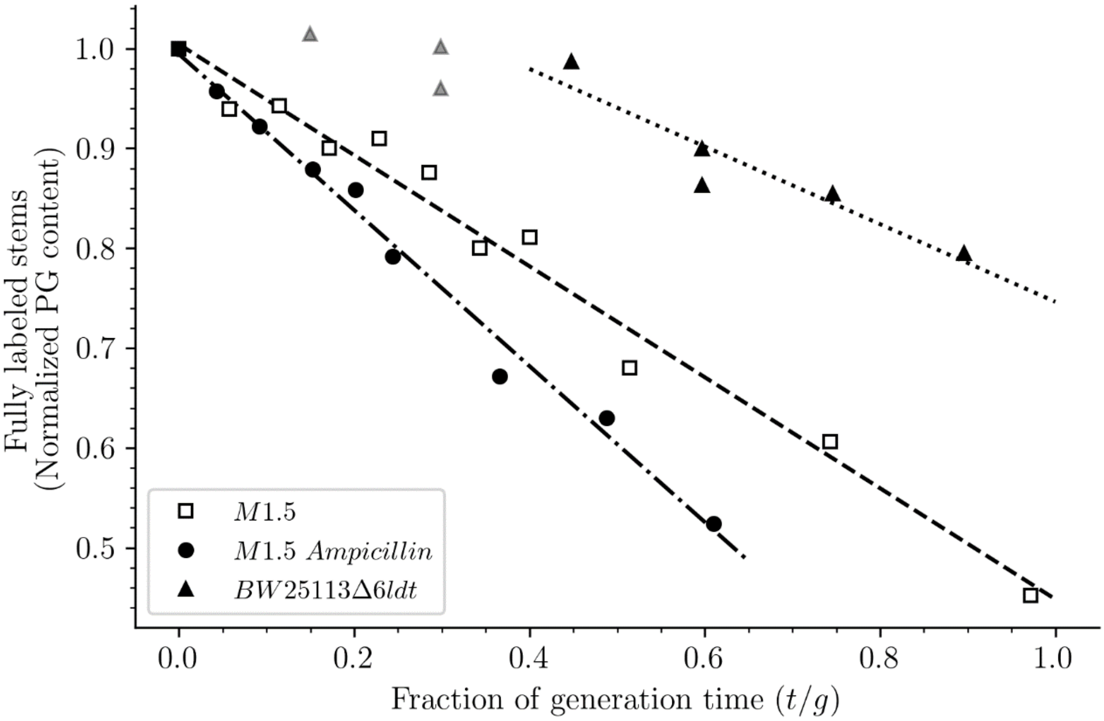


**Supplementary Fig. S13. Release of peptidoglycan fragments from sacculi**. Kinetics of the isotopologue composition of sacculi was used to determine the extent of the release of peptidoglycan fragments from sacculi. For this purpose, we considered that the decrease in the number of fully labeled disaccharide-peptide subunits is equal to the number of subunits released from the peptidoglycan. This is legitimate since peptidoglycan subunits are recycled in a large number of moieties including two glucosamine residues, two acetyl groups, one D-lactoyl residue, the L-Ala-D-iGlu-DAP tripeptide or its constitutive amino acids, and two D-Ala residues (Supplementary Fig. S1). Disaccharide-peptide units issued from recycling are therefore mixtures of labeled and unlabeled moieties that can be easily distinguished from fully labeled subunits. Of note, release of peptidoglycan fragments in the culture medium, corresponding to a small proportion of peptidoglycan degradation products (Goodell and Schwarz, 1985), also results in a reduction in the content of fully labeled disaccharide-peptides. Thus, the decrease in the absolute number of fully labeled disaccharide-peptide subunits in one generation directly provides an estimate of the fraction of the preexisting peptidoglycan that is degraded during each generation. For this reason, the decrease in the relative amount of fully labeled peptidoglycan disaccharide-peptides (normalized to 1.0 at the medium switch) was computed as a function of time (normalized for the generation time; Supplementary Table S2). The decrease was linear for M1.5 grown in the presence or absence of ampicillin. For BW25113Δ6*ldt*, a decrease in the fully labeled stems was only observed after 0.45 generation time. For this reason, regression analysis was performed for a portion of the kinetics (from 0.45 to 0.85 generation time; data points excluded from regression analysis are shown as grey triangles). The estimates of combined peptidoglycan turnover and recycling per generation were 39% for strain Δ6*ldt* and 56% or 78% for M1.5 grown in the absence or presence of ampicillin, respectively.
